# Role of autophagy in sepsis-induced skeletal muscle dysfunction, whole-body metabolism, and survival

**DOI:** 10.1101/2021.08.05.455081

**Authors:** Jean-Philippe Leduc-Gaudet, Kayla Miguez, Marina Cefis, Alaa Moamer, Tomer Jordi Chaffer, Julie Faitg, Olivier Reynaud, Felipe E Broering, Anwar Shams, Dominique Mayaki, Laurent Huck, Marco Sandri, Gilles Gouspillou, Sabah NA Hussain

**Affiliations:** Department of Critical Care and Translational Research in Respiratory Diseases Program, Research Institute of the McGill University Health Centre (MUHC), Montréal, QC, Canada; Meakins-Christie Laboratories, Department of Medicine, Faculty of Medicine, McGill University, Montréal, QC, Canada; Département des sciences de l’activité physique, Faculté des sciences, Université du Québec à Montréal (UQAM), Montréal, QC, Canada; Veneto Institute of Molecular Medicine (VIMM) and Department of Biomedical Science, Università di Padova, Padova, Italy; Department of Pharmacology, Faculty of Medicine, Taif University, Saudi Arabia

**Keywords:** autophagy, sepsis, skeletal muscle, atrophy, mitochondria, proteasome

## Abstract

Septic patients frequently develop skeletal muscle wasting and weakness, resulting in severe clinical consequences and adverse outcomes. Autophagy is a stress-induced degradative process essential to cell survival. Recent studies have demonstrated that sepsis triggers sustained induction of autophagy in skeletal muscles, although the impact of this enhanced autophagy on sepsis-induced muscle dysfunction remains unclear. *Atg7* is an autophagy gene that plays a major role in autophagosome formation. Using an inducible and muscle-specific *Atg7* knockout mouse model (Atg7^iSkM-KO^), we investigated the functional importance of skeletal muscle autophagy in sepsis. Sepsis was induced using cecal ligation and perforation (CLP) with a sham operation serving as a control. Atg7^iSkM-KO^ mice exhibited a more severe phenotype in response to sepsis, marked by severe muscle wasting and contractile dysfunction, hypoglycemia, higher ketone levels and a decreased in survival as compared to mice with intact *Atg7*. Several genes that encode 26S proteasome subunits were upregulated, suggesting that activation of the ubiquitin-proteasome system is responsible for the severe muscle atrophy that was seen in these mice. Sepsis and Atg7 deletion resulted in the accumulation of mitochondrial dysfunction, although sepsis did not further worsen mitochondrial dysfunction in Atg7^iSkM-KO^ mice. Overall, our study demonstrates that autophagy inactivation in skeletal muscles triggers significant worsening of sepsis-induced contractile and metabolic dysfunctions and negatively impacts survival. Induction of autophagy in skeletal muscles in response to sepsis thus represents a protective mechanism.

## Introduction

Sepsis is a leading cause of death in intensive care units. Several prospective and retrospective studies have confirmed that ventilatory and limb muscle dysfunction occurs in 70-100% of septic patients (1, 2). Ventilatory muscle dysfunction leads to difficult weaning of patients from mechanical ventilation and recurrence of respiratory failure after weaning (3, 4). Long-term ramifications of limb muscle dysfunction include functional limitations and poor quality of life (5). Sepsis-induced multiple organ failure and muscle inactivity are among the risk factors for development of severe muscle dysfunction. Primary manifestations include impaired contractility and atrophy of muscle fibers.

Sepsis-induced atrophy has been primarily attributed to increased protein degradation. Indeed, it has been shown in septic muscles of rats and mice that total and myofibrillar protein degradation increases by 50% and 440%, respectively (6), and that myofibrillar protein breakdown by the ubiquitin system is enhanced (7). However, it is now recognized that increased protein degradation, decreased contractility, and the development of atrophy that result from sepsis cannot be explained *in vivo* by proteasomal-mediated protein loss (7, 8).

Proteolysis in skeletal muscles is regulated by four pathways: the calpain, caspase-3, ubiquitin-proteasome, and autophagy pathways. The first three are responsible for degradation of myofilament proteins. Calpains and caspase-3 are strongly activated in limb and ventilatory muscle fibers during sepsis (9–11). Several components of the ubiquitin-proteasome system, including the muscle-specific ubiquitin E3 ligases *Fbxo32* (Atrogin-1) and *Trim63* (MuRF1) and 20S subunits, are upregulated in the ventilatory and limb muscles in animal models of sepsis (12–15), although there is no evidence that the proteasome *per se* is responsible for sepsis-induced skeletal muscle dysfunction.

The autophagy pathway is primarily responsible for the degradation of cytoplasmic proteins and organelles, including the mitochondria and peroxisomes. Autophagy plays an essential role in cell development and is generally thought of as a pro-survival process. Although it has been described in all cells, its role in skeletal muscle protein degradation has largely been ignored. Recently, however, several studies have demonstrated that autophagy itself is a critical regulator of protein homeostasis and mitochondrial quality in skeletal muscles, (16, 17) and that its contribution to total muscle protein degradation can be just as high as that from the proteasome pathway (18). Several groups have reported that autophagy is induced in skeletal muscles of mice and rats injected with bacterial lipopolysaccharide (LPS) and that its induction coincides with significant mitochondrial dysfunction, depressed muscle contractility, and the development of fiber atrophy (19–22). In a recent study, we demonstrated that cecal ligation and perforation (CLP)-induced sepsis triggers prolonged activation of autophagy and fiber atrophy in limb muscles (15). So, although there has been some progress in documenting autophagy in septic skeletal muscles, its contributions to sepsis-induced muscle atrophy remain largely unknown.

In this study, we generated conditional muscle-specific knockout of the autophagy related 7 (Atg7) gene in mice to address functional roles of autophagy in sepsis-induced skeletal muscle dysfunction. Our main hypotheses were that augmented autophagy serves as a protective mechanism designed to increase the recycling of dysfunctional organelles and protein aggregates inside septic muscles and that inactivation of autophagy exacerbates muscle loss during sepsis.

## Results

### Muscle-specific inactivation of autophagy

The physiological function of autophagy in septic skeletal muscles was investigated by crossing *Atg7*-floxed mice (*Atg7*^f/f^) with a transgenic line of mice in which Cre-ER^T2^ is expressed under the control of a tamoxifen-inducible human skeletal actin (HSA) promoter. The muscle-specific *Atg7* knockout mice generated from this cross are hereafter referred to as Atg7^iSkM-KO^ (Supplementary Fig. S1). Immunoblotting and PCR analyses showed that 48 h after undergoing a sham surgical operation, *Atg7* mRNA and ATG7 protein levels were markedly lower in the limb muscles of Atg7^iSkM-KO^ mice relative to Atg7^f/f^ mice, thereby confirming the efficacy of *Atg7* deletion (Fig. 1A-B). Lower LC3B-II (lipidated form of LC3B) protein levels and higher p62/SQSTM1 protein levels (autophagosome substrate) indicate that autophagosome formation was also inhibited, further confirming that *Atg7* deletion exerts an inhibitory effect on basal autophagy (Fig. 1A&C-D, Supplementary Fig. S2A-C). Heart ATG7 protein levels were unaffected, indicating that autophagy inactivation in this model is specific to skeletal muscles (Supplementary Fig. S2D). Any remaining traces of Atg7 (protein or mRNA) in the limb muscles likely came from endothelial cells, fibroblasts, macrophages, or blood cells, as *Atg7* is normally expressed in a variety of different tissues.

**Figure 1:**
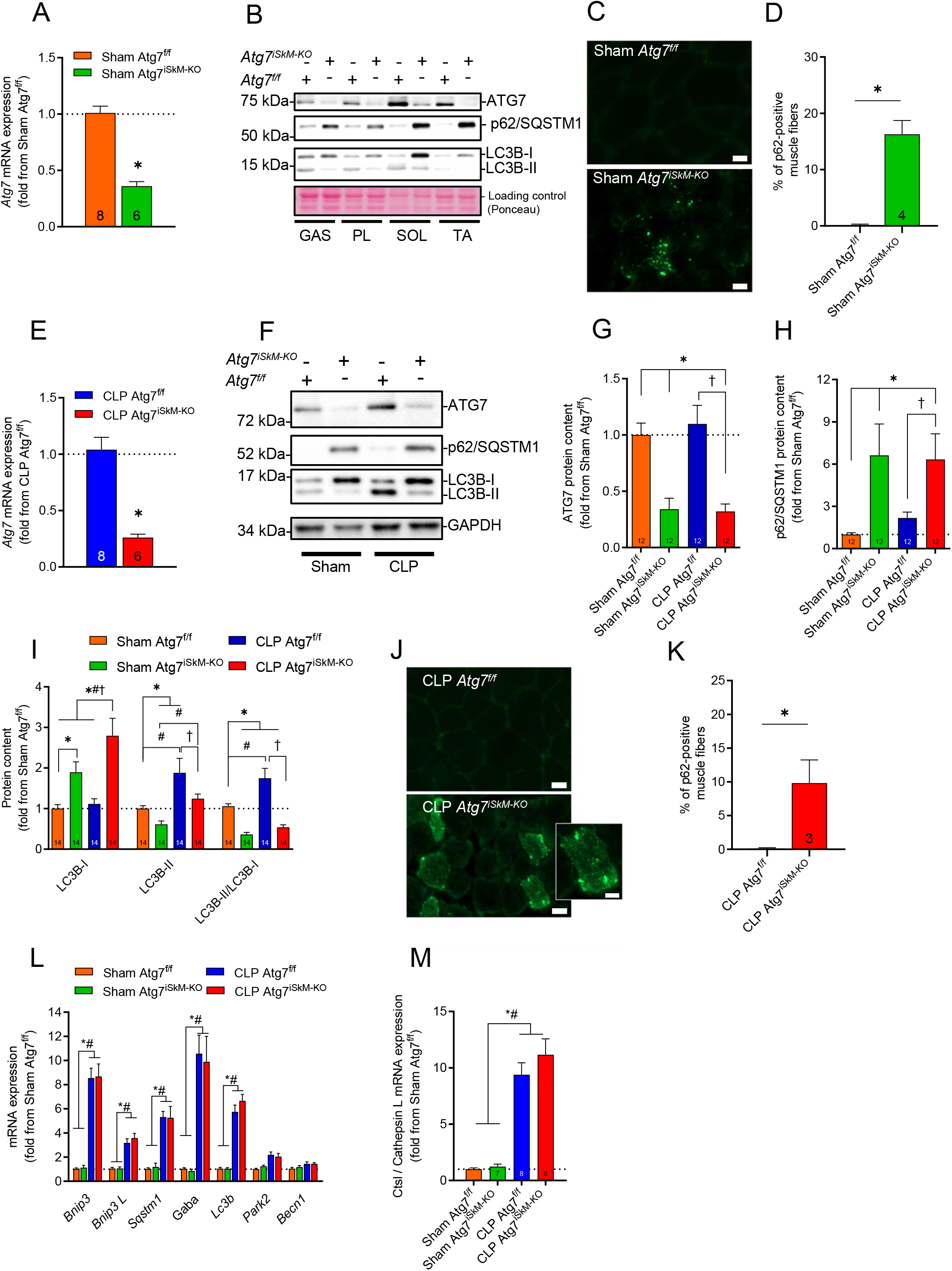
Muscle-specific deletion of Atg7 inhibits basal autophagy in skeletal muscles. A) mRNA expressions of Atg7 in TA muscles of female Atg7^f/f^ and Atg7^iSkM-KO^ mice 48 h after sham surgery. Data presented as fold change relative to sham Atg7^f/f^ mice. B) Representative immunoblots of Atg7, p62/SQSTM1, and LC3B proteins. GAS = gastrocnemius, PL = plantaris, SOL = soleus, TA = tibialis anterior. C-D) Immunostaining and quantification of the proportion of p62/SQSTM1 positive fibers in TA muscles of female Atg7^f/f^ and Atg7^iSkM-KO^ mice 48 h after sham surgery. Scale bars = 20µm. E) mRNA expressions of Atg7 in TA muscles of female Atg7^f/f^ and Atg7^iSkM-KO^ mice 48 h after CLP. F) Representative immunoblots of ATG7, p62/SQSTM1, and LC3B proteins in TA muscles of female Atg7^f/f^ and Atg7^iSkM-KO^ mice 48 h after sham surgery or CLP. G-I) Densitometric analyses of ATG7, p62/SQSTM1 and LC3B protein levels in TA muscles of female Atg7^f/f^ and Atg7^iSkM-KO^ mice 48 h after sham surgery or CLP. J-K) Immunostaining for p62/SQSTM1 in TA muscles of female Atg7^f/f^ and Atg7^iSkM-KO^ mice 48 h after CLP surgery. Scale bars = 20 µm, inset scale bar = 10 µm. L) mRNA expressions of various autophagy-related genes (M) and lysosomal cathepsin in TA muscles of female Atg7^f/f^ and Atg7^iSkM-KO^ mice 48 h after sham surgery or CLP. Gaba refers to Gabarapl1 and Becn1 refers to Beclin1. N = 6-8 group. Data in panels A, D, E, G, H, I, K, L and M presented as mean ± SEM. Number of animals indicated within bars, where applicable. * p <0.05 vs sham Atg7^f/f^; # p <0.05 for sepsis effect (i.e. sham Atg7^f/f^ vs. CLP Atg7^f/f^ or sham Atg7^iSkM-KO^ vs. CLP Atg7^iSkM-KO^); † p < 0.05 for sepsis plus knockout effect (i.e. CLP Atg7^f/f^ vs. CLP Atg7^iSkM-KO^).

We have previously reported that autophagy increases in the limb muscles of mice 48 h after sepsis is induced by CLP (15, 23). In line with our previous observations, we found that sepsis resulted in a significant increase in LC3BII levels and LC3B-II/LC3B-I ratio in various skeletal muscles (Supplementary Fig. 3A-D). To further define the impact of sepsis on autophagy, a leupeptin-based assay of the autophagic flux was conducted (Supplementary Fig. 3E). In septic animals, leupeptin treatment resulted in a further increase in the LC3B-II/I ratio (Supplementary Fig. 3F), indicating that autophagy flux increases in skeletal muscles upon sepsis induction.

To assess the functional importance of skeletal muscle autophagy in sepsis, sham operated and CLP-operated Atg7^f/f^ and Atg7^iSkM-KO^ mice were compared. Under septic conditions, *Atg7* mRNA expression, ATG7 protein level and LC3B-II/LC3B-I protein ratio was significantly lower in the limb muscles of Atg7^iSkM-KO^ mice as compared to Atg7^f/f^ mice (Fig. 1E-I). The higher levels of p62/SQSTM1 and LC3-I associated with the lower LC3B-II protein levels in muscles of Atg7^iSkM-KO^ mice suggests that autophagosome formation is blocked in this mouse model in response to sepsis (Fig. 1F-I). Importantly, p62/SQSTM1 protein accumulated in muscles of Atg7^iSkM-KO^ mice but not in those of Atg7^f/f^ mice (Fig. 1J&K), providing further evidence of selective inactivation of autophagy via deletion of *Atg7*.

Further confirming that autophagy is induced in the tibialis anterior (TA) and gastrocnemius (GAS) muscles of Atg7^f/f^ mice 48 h after CLP, protein levels of LC3BII, LC3B-II/LC3B-I ratio and Bnip3, as well as mRNA expression of several autophagy-related genes (*Lc3b*, *Sqstm1*, *Bnip3*, *Bnip3L*), and the lysosomal gene *Ctsl* all increased in septic Atg7^f/f^ muscles (Fig. 1F-M; Supplementary Fig. S4A-F). Expressions of two autophagy-related genes, *Gabarapl1* and *Park2*, were differentially regulated, depending on the muscle; in the TA, *Gabarapl1* expression increased while *Park2* did not change (Fig. 1L), whereas in the GAS, *Park2* increased while *Gabarapl1* did not change (Supplementary Fig. S4E). LC3B-II/LC3B-I ratio did not increase in the limb muscles of Atg7^iSkM-KO^ mice in response to sepsis, but p62/SQSTM1 protein level did (Fig. 1F-I and Supplementary Fig. S4C&D). In response to sepsis, p62/SQSTM1 protein aggregates were observed in limb muscles of Atg7^iSkM-KO^ mice (Fig. 1J&K), suggesting that it accumulates because of autophagy inhibition. Interestingly, in response to sepsis, autophagy-related gene expressions in the TA and GAS muscles of Atg7^f/f^ and Atg7^iSkM-KO^ mice increased to a similar extent (Fig. 1L&M, and Supplementary Fig. S2E&F), suggesting that deletion of *Atg7* does not interfere with the transcriptional upregulation of these genes that occurs in severe sepsis.

### The impact of sepsis and muscle-specific inactivation of autophagy on body mass, disease severity, and mortality

Although female and male Atg7^iSkM-KO^ mice appeared phenotypically normal prior to sham and CLP surgical procedures, their body mass were lower compared to age-matched Atg7^f/f^ mice (Supplementary Fig. S5A&B). These lower body mass cannot be attributed to the tamoxifen diet alone since wild-type mice that had been fed the same diet were relatively heavier (Supplementary Fig. S5A). The effect is mainly due to a decrease in lean body mass (Supplementary Fig. S5C), yet despite having less lean body mass, the Atg7^iSkM-KO^ mice had similar *in vivo* muscle grip strength as compared to Atg7^f/f^ mice (Supplementary Fig. S5D&E). Both female and male mice lost body mass in response to 48 h of sepsis, an effect which was more pronounced in Atg7^iSkM-KO^ mice than in Atg7^f/f^ mice (Fig. 2A).

**Figure 2:**
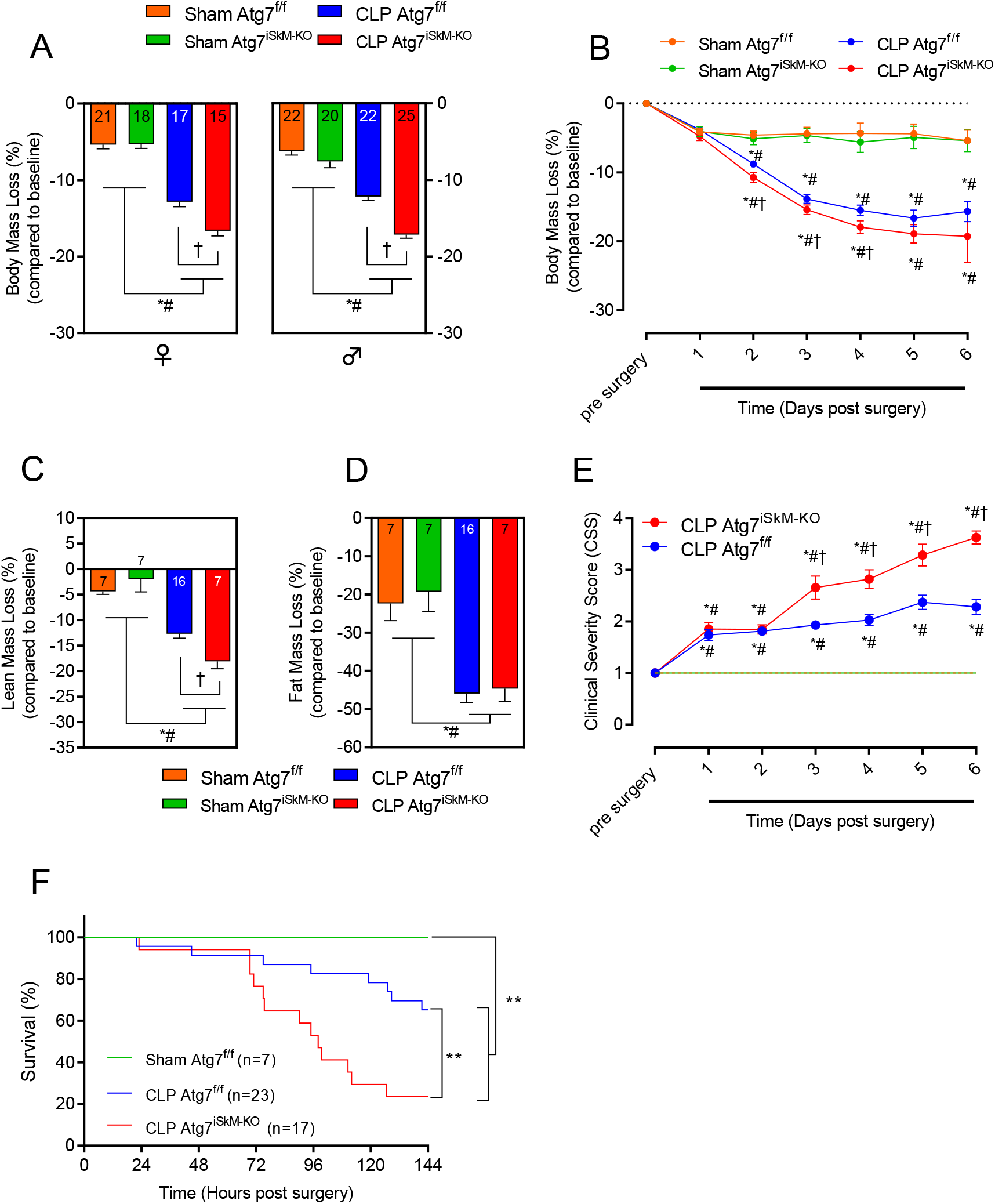
Autophagy inactivation exacerbates disease severity. A) Percent change in body mass of female (left panel) and male (right panel) Atg7^f/f^ and Atg7^iSkM-KO^ mice in response to acute sepsis. Body mass measured prior to sham surgery or CLP then 48 hours later. B) Percent change in body mass of female Atg7^f/f^ and Atg7^iSkM-KO^ mice in response to prolonged sepsis. Body mass measured prior to sham surgery or CLP then daily over a 6-day period. C-D) Percent change in lean (C) and fat (D) muscle mass of female Atg7^f/f^ and Atg7^iSkM-KO^ mice in response to prolonged sepsis. Mass measured prior to sham surgery or CLP then daily over a 4-day period. E) Clinical severity score (CSS) for female Atg7^f/f^ and Atg7^iSkM-KO^ mice in response to prolonged sepsis. CSS measured prior to sham surgery or CLP then daily over a 6-day period. F) Percent survival of female Atg7^f/f^ and Atg7^iSkM-KO^ mice in response to prolonged sepsis. Survival was monitored over a 6-day period following sham surgery or CLP. No mortality of sham Atg7^iSkM-KO^ mice was observed. Comparisons of curves indicated by **p < 0.001. Data in panels A-E presented as mean ± SEM. Number of animals indicated in bars, where applicable. * p <0.05 vs sham Atg7^f/f^; # p <0.05 for sepsis effect (i.e. sham Atg7^f/f^ vs. CLP Atg7^f/f^ or sham Atg7^iSkM-KO^ vs. CLP Atg7^iSkM-KO^); † p < 0.05 for sepsis plus knockout effect (i.e. CLP Atg7^f/f^ vs. CLP Atg7^iSkM-^KO_)._

We next determine the effects of sustained autophagy inactivation on sepsis-induced changes in body composition, clinical signs of sepsis, and survival rates over six days following the sham surgery or CLP procedure. Sham Atg7^f/f^ and Atg7^iSkM-KO^ mice showed relatively mild body mass loss one day post-surgery, but body mass remained unchanged thereafter (Fig. 2B). Both Atg7^f/f^ and Atg7^iSkM-KO^ mice progressively lost body mass in response to sepsis. However, body mass loss was more pronounced in septic Atg7^iSkM-KO^ mice because they experienced greater losses in lean body mass as compared to Atg7^f/f^ mice (Fig. 2C). Sepsis-induced losses of whole-body fat were similar in both groups of mice (Fig. 2D). Sham Atg7^f/f^ and Atg7^iSkM-KO^ mice showed no signs of sickness following surgery (Fig. 2E). In response to sepsis, Atg7^iSkM-KO^ mice exhibited more clinical signs of illness as compared to Atg7^f/f^ mice. These differences became more apparent 72 h post-CLP (Fig. 2E). No mortality was observed in sham Atg7^f/f^ or Atg7^iSkM-KO^ mice. Survival rates in the 6-day period post-CLP were lower for Atg7^iSkM-KO^ mice than for Atg7^f/f^ mice (65% vs 23%, respectively) (Fig. 2F). These data collectively indicate that autophagy inactivation in skeletal muscles leads to losses of body mass and lean muscle mass, exacerbates sepsis, and increases sepsis-induced mortality.

### The impact of sepsis and muscle-specific inactivation of autophagy on fiber atrophy

The effects of autophagy inactivation and sepsis on muscle atrophy and contractile performance were quantified by measuring muscle mass, fiber type composition, and fiber size of the TA and GAS muscles and by measuring *in-situ* contractility of the TA. TA and GAS mass were lower in sham Atg7^iSkM-KO^ mice as compared to sham Atg7^f/f^ mice; this difference was independent of sex (Fig. 3A&B). In response to sepsis, TA and GAS masses were lower in Atg7^f/f^ and Atg7^iSkM-KO^ mice as compared to corresponding sham mice and lower in septic Atg7^iSkM-KO^ mice relative to septic Atg7^f/f^ mice (Fig. 3A&B). Neither autophagy inactivation nor sepsis had any effect on muscle fiber type composition (Supplementary Fig. S5F-H). TA fiber diameter in sham Atg7^iSkM-KO^ mice was smaller vs sham Atg7^f/f^ mice (Fig. 2C-E and Supplementary Fig. 6A-D). In response to sepsis, TA fiber diameters in Atg7^f/f^ and Atg7^iSkM-KO^ mice were lower relative to sham Atg7^f/f^ mice and lower in septic Atg7^iSkM-KO^ mice relative to septic Atg7^f/f^ mice (Fig. 3C-D). TA fiber diameter measured 6 days post-surgery indicated that sepsis-induced fiber atrophy is more severe in Atg7^iSkM-KO^ mice vs Atg7^f/f^ mice (Fig. 3E and Supplementary Fig. 4D). These results collectively demonstrate that autophagy inactivation results in worsening of sepsis-induced fiber atrophy in skeletal muscles and that this effect is not a consequence of changes in fiber type composition.

**Figure 3:**
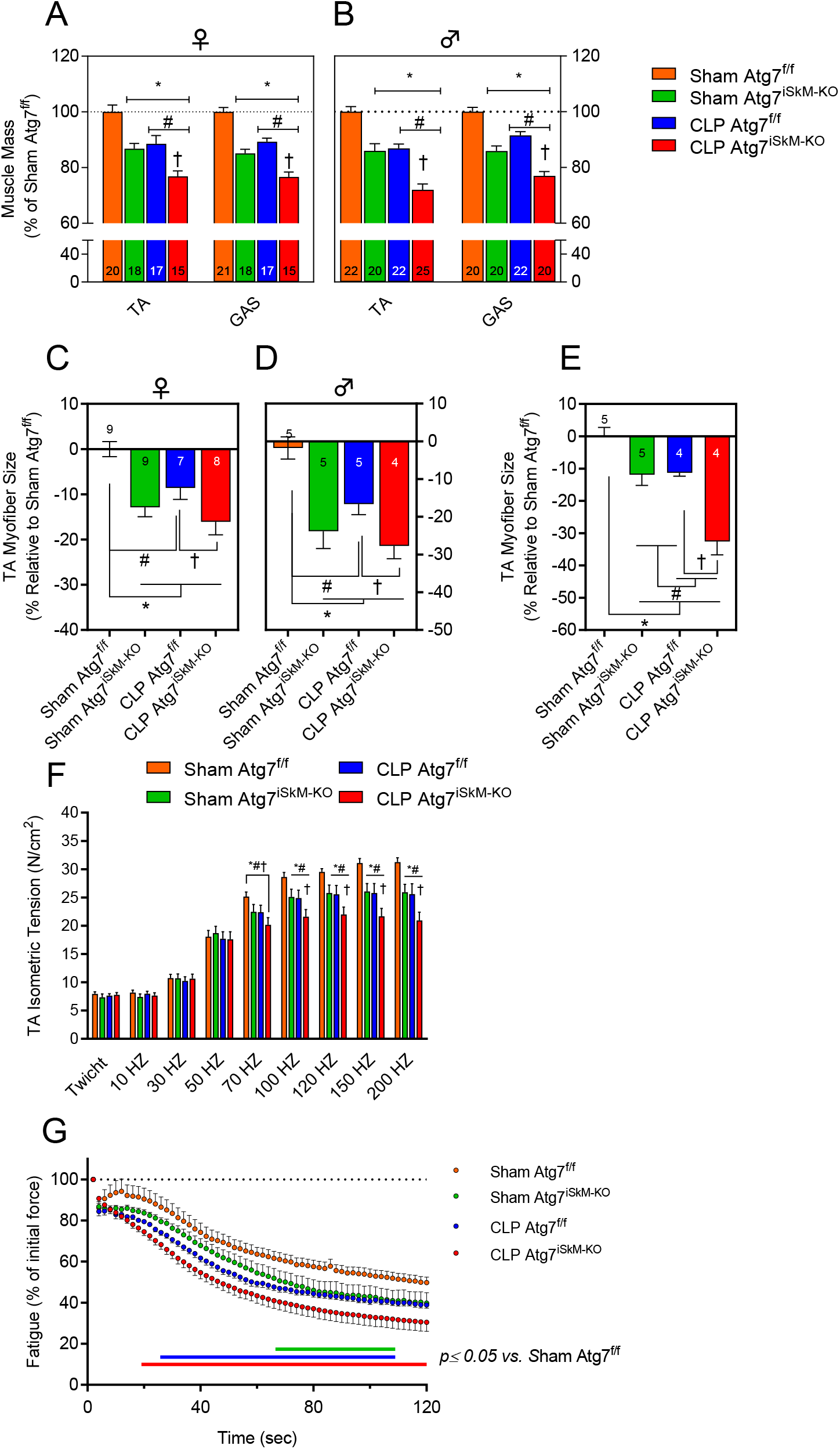
Autophagy inactivation exacerbates muscle wasting and weakness. A-B) Muscle mass loss in TA and GAS muscles of female (A) and male (B) Atg7^f/f^ and Atg7^iSkM-KO^ mice 48 h after sham surgery or CLP. Data presented as percent change relative to sham Atg7^f/f^ mice. C-E) Myofiber size of TA muscles of female (C&E) and male (D) Atg7^f/f^ and Atg7^iSkM-KO^ mice 48 h (C&D) or 6 days (E) after sham surgery or CLP. F) Isometric tension of TA muscles of female and male Atg7^f/f^ and Atg7^iSkM-KO^ mice 48 h after sham surgery or CLP. N = 20-23 per group). (G) Fatigue in TA muscles of female Atg7^f/f^ and Atg7^iSkM-KO^ mice 48 h after sham surgery or CLP. Fatigue curves presented as percent change from tetanus 1 (initial force) to tetanus 60, n = 17, sham Atg7^f/f^; n = 14, sham Atg7^iSkM-KO^; n = 11, CLP Atg7^f/f^; n = 12, CLP Atg7^iSkM-KO^. Data in panels A-G presented as mean ± SEM. Number of animals indicated in bars, where applicable. * p <0.05 vs sham Atg7^f/f^; # p <0.05 for sepsis effect (i.e. sham Atg7^f/f^ vs. CLP Atg7^f/f^ or sham Atg7^iSkM-KO^ vs. CLP Atg7^iSkM-KO^); † p < 0.05 for sepsis plus knockout effect (i.e. CLP Atg7^f/f^ vs. CLP Atg7^iSkM-KO^).

Peak isometric tension was lower in the TA muscles of sham Atg7^iSkM-KO^ mice than it was in sham Atg7^f/f^ mice (Fig. 3F and Supplementary Fig. 7A-D). In septic muscles, peak isometric tension was lower in Atg7^f/f^ and Atg7^iSkM-KO^ mice than it was in corresponding sham mice, and lower in Atg7^iSkM-KO^ mice than it was in Atg7^f/f^ mice (Fig. 3F). These differences in contractility were not a consequence of the tamoxifen diet per se, as peak isometric tension was lower in septic muscles of Atg7^iSkM-KO^ mice than in septic muscles of WT tamoxifen-fed mice (Supplementary Fig. S7E). TA fatigability was assessed by subjecting muscles to 60 repeated isometric contractions with 2 seconds between contractions (120s total duration). TA muscle tension progressively decreased in all mice. The fatigue effect was greater in sham Atg7^iSkM-KO^ mice than it was in sham Atg7^f/f^ mice (60 and 50% of initial tension at 120s, respectively) (Fig. 3G), and greater in Atg7^f/f^ and Atg7^iSkM-KO^ mice than it was in corresponding sham mice. In response to sepsis, fatiguability was greater in Atg7^iSkM-KO^ mice vs Atg7^f/f^ mice (Fig. 3G). These results demonstrate that autophagy is essential to the maintenance of skeletal muscle contractility and that its inactivation exacerbates sepsis-induced muscle contractile dysfunction.

### The impact of sepsis and muscle-specific inactivation of autophagy on mitochondrial integrity and function

No differences in state II (basal, ADP-restricted respiration) mitochondrial respiration were detected across groups (Fig. 4A). State III mitochondrial respiration (maximal ADP-stimulated) driven by complex I substrates (glutamate plus malate) was 14% lower in septic muscles of Atg7^f/f^ mice as compared to corresponding sham mice, and lower in sham and septic Atg7^iSkM-KO^ mice as compared to corresponding Atg7^f/f^ mice (Fig. 4A). Mitochondrial coupling efficiency was unaffected by Atg7 deletion or sepsis since no differences were observed in acceptor control ratios (ARC) (State III/State II respiration) (Fig. 4B). Immunoblotting for representative subunits of the oxidative phosphorylation (OXPHOS) pathway showed that there was an overall increase in OXPHOS proteins in Atg7^iSkM-KO^ mice as compared to Atg7^f/f^ mice (Supplementary Fig. S8A). This effect is unlikely to be a result of an increase in mitochondrial biogenesis since PGC1α expressions were similar in all groups (Supplementary Fig. S8B).

**Figure 4:**
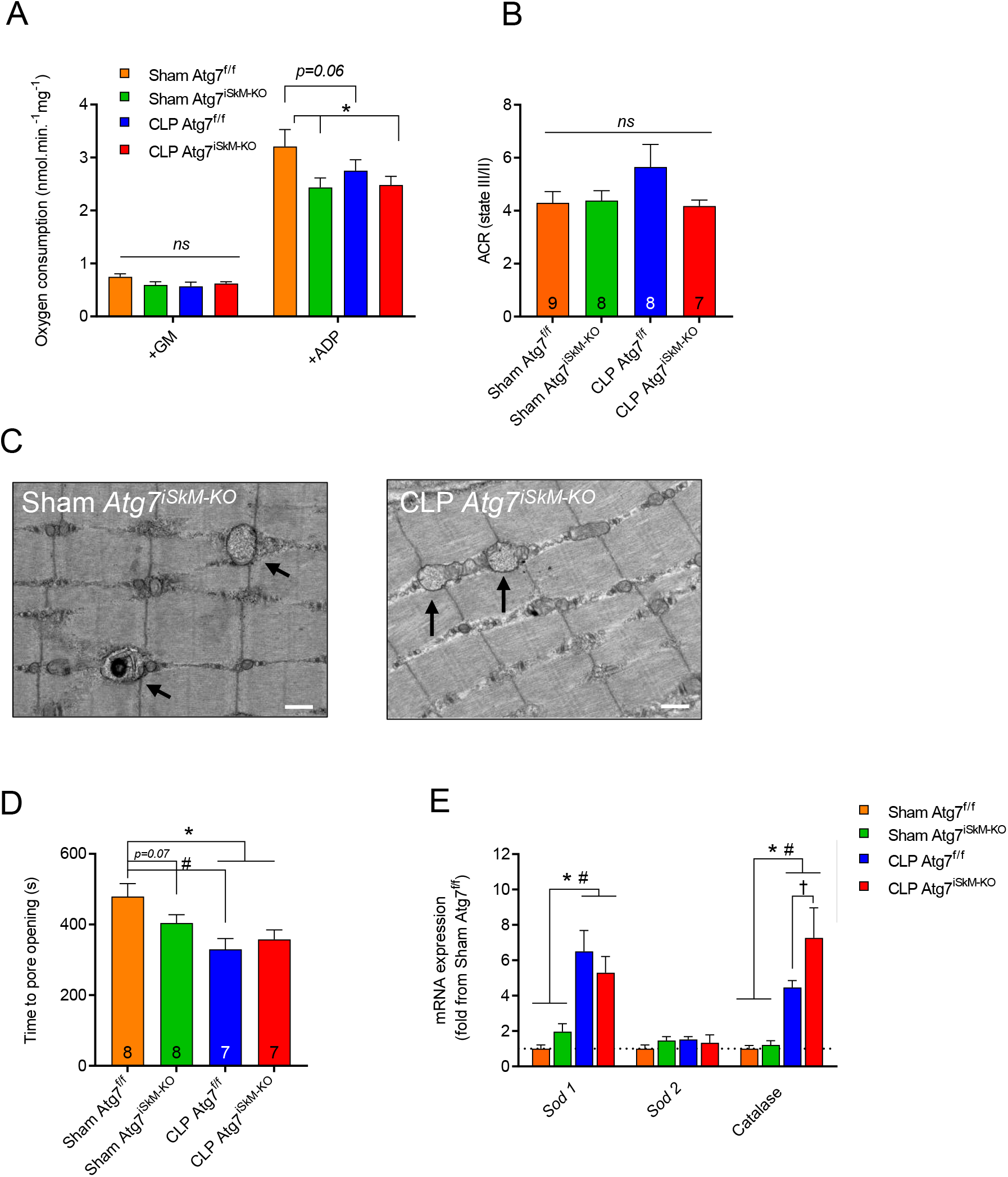
Autophagy is critical to mitochondrial homeostasis and skeletal muscle endurance. A) Mitochondrial oxygen consumption in permeabilized myofibers from GAS muscles of female Atg7^f/f^ and Atg7^iSkM-KO^ mice 48 h after sham surgery or CLP. Data normalized per mg of muscle. B) Acceptor control ratio (ACR) (index of mitochondrial coupling efficiency) in GAS muscles of female Atg7^f/f^ and Atg7^iSkM-KO^ mice 48 h after sham surgery or CLP. C) Time of opening of the mPTP in permeabilized myofibers from GAS muscles of female Atg7^f/f^ and Atg7^iSkM-KO^ mice 48 h after sham surgery or CLP. D) mRNA expressions of antioxidant defense-related genes in GAS muscles of female Atg7^f/f^ and Atg7^iSkM-KO^ mice 48 h after sham surgery or CLP, n = 6-8 per group. **(E)** Electron micrographs showing accumulation of abnormal mitochondria (black arrows) in GAS muscles of female Atg7^iSkM-KO^ mice 48 h after sham surgery or CLP. Scale bars = 0.5 µm. Data in panels A-D presented as mean ± SEM. Number of animals indicated in bars, where applicable. * p <0.05 vs sham Atg7^f/f^; # p <0.05 for sepsis effect (i.e. sham Atg7^f/f^ vs. CLP Atg7^f/f^ or sham Atg7^iSkM-KO^ vs. CLP Atg7^iSkM-KO^); † p < 0.05 for sepsis plus knockout effect (i.e. CLP Atg7^f/f^ vs. CLP Atg7^iSkM-KO^).

To test the possibility that inactivation of autophagy resulted in accumulation of dysfunctional and/or damaged mitochondria, we then looked for ultrastructural abnormalities by transmission electron microscopy in GAS muscles of Sham Atg7^iSkM-KO^ and CLP Atg7^iSkM-KO^ mice. As expected, abnormal mitochondria with disorganized cristae and major ultrastructural abnormalities were detected in autophagy deficient skeletal muscles (Fig. 4C and Supplementary Fig. S8E). This is consistent with previous observations that autophagy inactivation results in the accumulation of abnormal mitochondria and autophagic vesicles, consistent with blockage of autophagy (24–28).

Time to mitochondrial permeability transition pore (mPTP) opening was 12% shorter in muscles of sham Atg7^iSkM-KO^ relative to sham Atg7^f/f^ mice, indicating that autophagy ablation sensitized mitochondrial to mPTP opening. In Atg7^f/f^ mice, sepsis resulted in a decrease in the time to mPTP opening (Fig. 4D). In Atg7^iSkM-KO^ mice, sepsis did not further shorten the time to mPTP opening (Fig. 4D). These results indicate that both autophagy inactivation and sepsis enhance the susceptibility to mPTP opening in skeletal muscles. Yet, despite significant changes in mitochondrial respiratory capacity and mPTP opening time, there were no significant effects of autophagy inactivation nor sepsis on mitochondrial hydrogen peroxide (H2O2) production (Supplementary Fig. S8C). The degree of oxidative stress in the muscles of each group was evaluated by measuring the levels of three antioxidant enzymes (*Sod1*, *Sod2*, and *catalase*). No significant differences in *Sod1*, *Sod2*, and *catalase* mRNA levels were detected in the GAS and TA muscles of sham Atg7^f/f^ and Atg7^iSkM-KO^ mice (Fig. 4E and Supplementary Fig. S8D). In contrast, sepsis resulted in an increase in *Sod1* and *catalase* mRNA levels in Atg7^f/f^ and Atg7^iSkM-KO^ mice relative to corresponding sham mice (Fig. 4E and Supplementary Fig. S8D). *Catalase* levels were also higher in the GAS muscle of septic Atg7^iSkM-KO^ mice relative to septic Atg7^f/f^ mice (Fig. 4E and Supplementary Fig. S8D). These results confirm the view that oxidative stress develops in septic skeletal muscles. They also suggest that autophagy inactivation does not exacerbate sepsis-induced muscle oxidative stress.

### Impact of sepsis and muscle-specific inactivation of autophagy on the ubiquitin-proteasome pathway

Previous studies have shown that inactivation of autophagy in skeletal muscles triggers significant increases in 26S proteasome activity (16). To determine if this occurred in the skeletal muscles of Atg7^iSkM-KO^ mice, ubiquitin E3 ligase expressions and 11S, 19S, and 20S subunits of the 26S proteasome were measured 48 h post-surgery in the TA and GAS of sham-operated and septic animals. No significant differences in the expressions of tripartite motif containing 32 (*Trim32*), *Nedd4*, *Fbxo32*, *Fbxo30* (Musa1), and *Trim63* were observed between sham Atg7^f/f^ and Atg7^iSkM-KO^ mice, nor were any observed for proteasome 26S subunits *Psmd11* and *Psmd8* (19S) or the constitutive and inducible β subunits of 20S (Fig. 5A-G, Supplementary Fig. S9A&B). However, two 11S subunits, proteasome activator subunits 1 and 2 (*Psme1* and *Psme2*), one constitutive 20S subunits of 20S, subunit beta 5 (*Psmb5*) and one 20S β subunit, proteasome 20S subunit beta 10 (*Psmb10*), were upregulated in Atg7^iSkM-KO^ mice, relative to Atg7^f/f^ mice (Fig. 5D&G).

**Figure 5:**
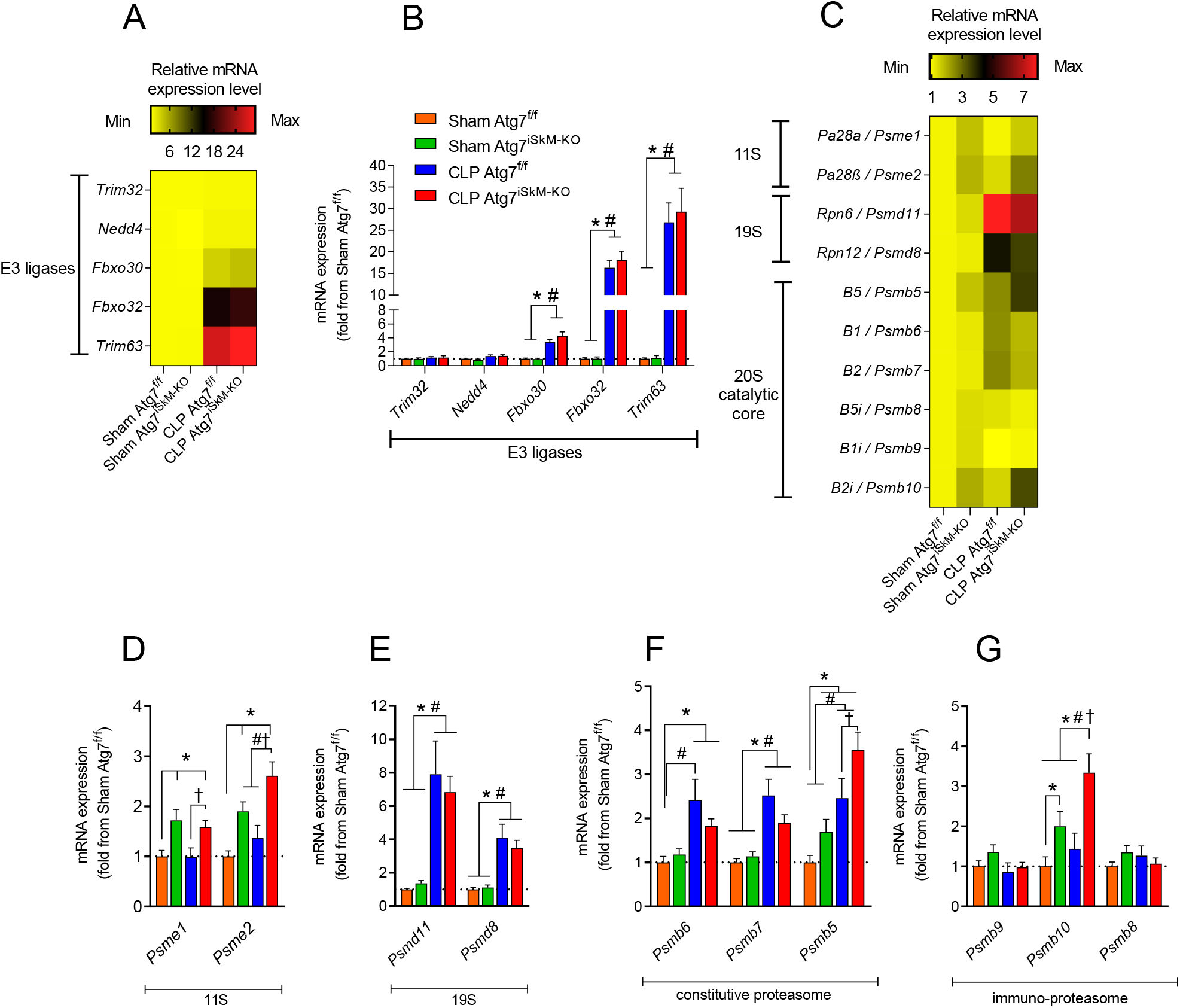
Sepsis-induced muscle proteolysis. A) Heat map summarizing relative mRNA expressions of various E3 ubiquitin ligases involved in skeletal muscle atrophy in TA muscles of female Atg7^f/f^ and Atg7^iSkM-KO^ mice 48 h after sham surgery or CLP. Colors indicate relative expression levels; red indicates high expression and yellow indicates low expression. B) mRNA expressions of E3 ubiquitin ligases in TA muscles of female Atg7^f/f^ and Atg7^iSkM-KO^ mice 48 h after sham surgery or CLP. C) Heat map summarizing relative mRNA expressions of various proteasome subunits. Colors indicate relative expression levels; red indicates high expression and yellow indicates low expression. D-G) mRNA expressions of proteasome subunits genes in TA muscles of female Atg7^f/f^ and Atg7^iSkM-KO^ mice 48 h after sham surgery or CLP. 18S levels used as a control. as fold change relative to sham Atg7^f/f^. Data in panels B and D-I presented as mean ± SEM. * p <0.05 vs sham Atg7^f/f^; # p <0.05 for sepsis effect (i.e. sham Atg7^f/f^ vs. CLP Atg7^f/f^ or sham Atg7^iSkM-KO^ vs. CLP Atg7^iSkM-KO^); † p < 0.05 for sepsis plus knockout effect (i.e. CLP Atg7^f/f^ vs. CLP Atg7^iSkM-KO^).

In sepsis, *Fbxo30*, *Fbxo32*, *Trim63*, *Psmd11, Psmd8*, and 2 constitutive 20S subunits of 20S, subunit beta 2 and 5 (*Psmb7*, *Psmb5*), were upregulated in Atg7^f/f^ and Atg7^iSkM-KO^ mice as compared to corresponding sham mice (Fig. 5A-G and Supplementary Fig. S9 A-C). Because skeletal muscle mass is regulated by the balance between protein synthesis and degradation, we measured *in vivo* protein synthesis using SUnSET method (29). No significant differences in the rate of protein synthesis were observed across all groups (Supplementary Fig. S9 D&E). Collectively, these results suggest that autophagy inactivation in skeletal muscles coincides with increased activity of specific subunits of the 26S proteasome but does not affect the expression of ubiquitin E3 ligases or protein synthesis. Sepsis, in contrast, does trigger upregulation of the E3 ligases, as well as specific subunits of the 19S and 20S proteasome.

### Sepsis and autophagy inhibition enhance proinflammatory cytokines expression in skeletal muscle

Several muscle-derived pro-inflammatory cytokines such as tumor necrosis factor (TNF-α) and interleukin 1 have been shown to negatively affect muscle function, including contractile performance. In this study, expressions of several of these cytokines and their receptors were measured in each group of mice. *Tweak*, *IL-1b*, *Tnf-α R*, and *IL-18* expressions were higher in the GAS of sham Atg7^iSkM-KO^ mice relative to sham Atg7^f/f^ mice and, in response to sepsis, *Tweak*, *Fn14*, *IL-6R*, *IL-1b*, *Tnf-α, Tnf-α R*, *IL-18*, *IL-18R,* and *IL-24* expressions were higher in septic Atg7^f/f^ and Atg7^iSkM-KO^ mice as compared to corresponding sham mice (Fig. 6A-F). No difference was observed between Atg7^f/f^ and Atg7^iSkM-KO^ mice in relation to the degree to which these cytokines and receptors were induced by sepsis. Using ELISA, we observed a trend for an increase in muscle IL-1β protein levels in sham Atg7^iSkM-KO^ mice relative to sham Atg7^f/f^ (Fig. 6G). Surprisingly, lower IL-1β protein levels were observed in CLP Atg7^iSkM-KO^ relative to the three groups. No difference in TNF-α protein level were detected across all groups (Fig. 6H). These results suggest that autophagy exerts a negative effect on muscle-derived pro-inflammatory cytokine levels under normal conditions but does not impact their patterns of expression in response to sepsis. Expressions of *mstn* (encodes myostatin protein) and the myokine fibroblast factor 21 (*Fgf21*) were also measured in the GAS: *mstn* expression was higher in sham Atg7^iSkM-KO^ mice as compared to Atg7^f/f^ mice and, in response to sepsis, *mstn* and *Fgf21* expressions were higher in Atg7^iSkM-KO^ and Atg7^f/f^ mice as compared to corresponding sham mice (Fig. 6G&H). Importantly, changes in *Fgf21* gene expression did not translate at the protein level as no differences in FGF21 protein levels were observed across groups (Fig. 6L).

**Figure 6:**
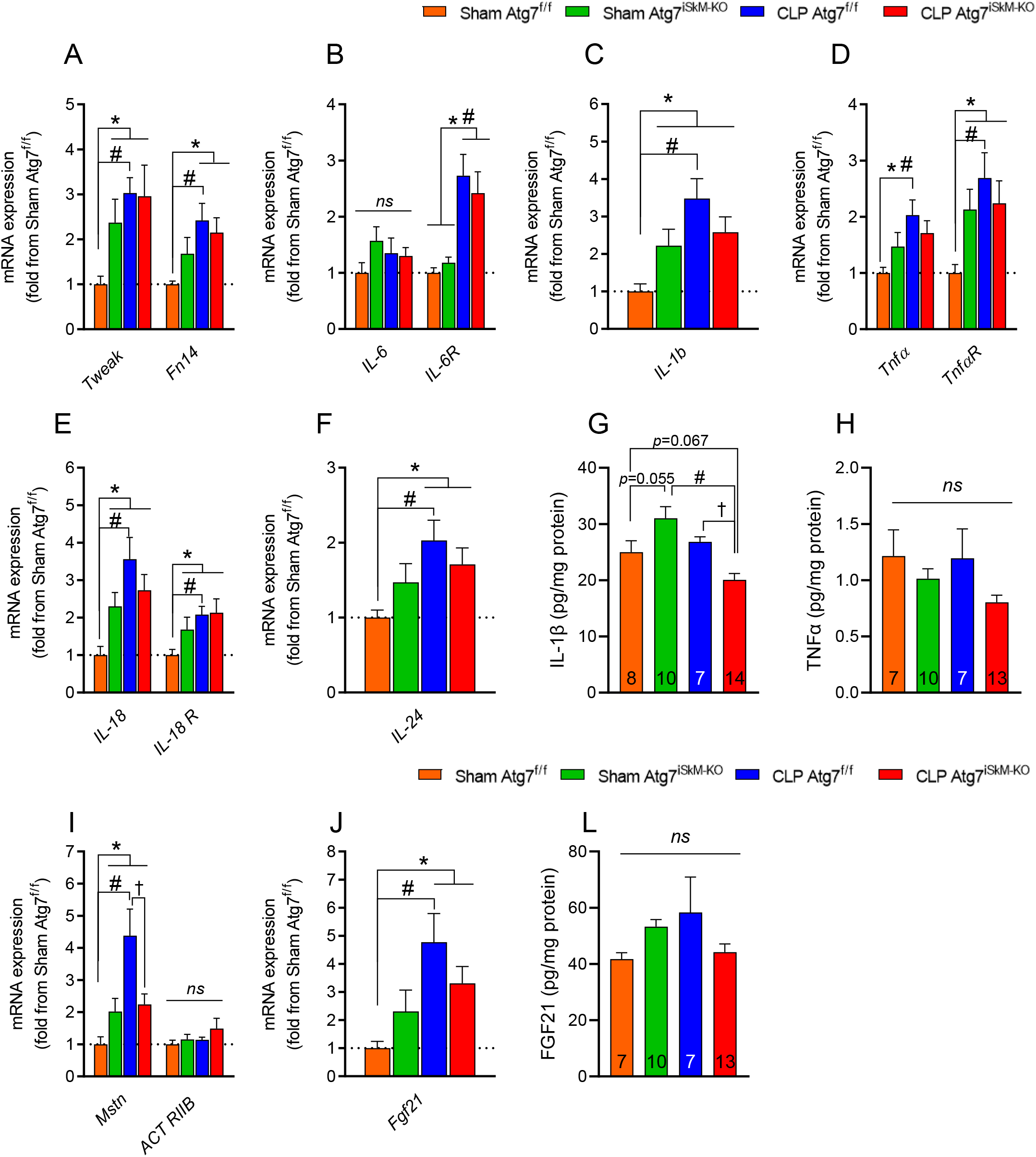
Sepsis upregulates proinflammatory cytokines and myostatin pathway. A-F) mRNA expressions of genes involved in the inflammatory response in GAS muscles of female Atg7^f/f^ and Atg7^iSkM-KO^ mice 48 h after sham surgery or CLP. G-H) IL-1β and TNFα protein level in skeletal muscle measured by ELISA. mRNA expressions of *Mtsn*, *ACT RIIB* and *Fgf21* in GAS muscles of female Atg7^f/f^ and Atg7^iSkM-KO^ mice 48 h after sham surgery or CLP. L) FGF21 protein level in skeletal muscle measured by ELISA. Data presented as fold change relative to sham Atg7^f/f^ and as mean ± SEM. Number of animals indicated in bars, where applicable. For panels showing mRNA expression data n = 6-8 per group. * p <0.05 vs sham Atg7^f/f^; # p <0.05 for sepsis effect (i.e. sham Atg7^f/f^ vs. CLP Atg7^f/f^ or sham Atg7^iSkM-KO^ vs. CLP Atg7^iSkM-KO^); † p < 0.05 for sepsis plus knockout effect (i.e. CLP Atg7^f/f^ vs. CLP Atg7^iSkM-KO^).

### Potential mechanisms underlying the prolonged effects of autophagy inactivation on sepsis-induced muscle dysfunction

Following prolonged sepsis, Atg7^iSkM-KO^ mice developed more severe limb muscle fiber atrophy than Atg7^f/f^ mice. To determine why, transcriptomic analyses were performed on TA samples from Atg7^f/f^ and Atg7^iSkM-KO^ mice six days after sepsis was induced. In Atg7^iSkM-KO^ mice, 451 genes were upregulated and 323 were downregulated in comparison to Atg7^f/f^ mice (FDR-adjusted *p*< 0.05; Fig. 7A). Gene ontology (GO) analyses of differentially expressed genes indicate that cellular processes such as catabolism, stress response, ribonucleoprotein synthesis, and intracellular protein transport were highly augmented in septic muscles Atg7^iSkM-KO^ mice (Fig. 7B). Catabolism-associated genes involved in muscle atrophy were highly upregulated, including *Trim63*, *Fbxo32*, and *Fbxo31* and *Psma3*, *Psma4*, *Psma5*, *Psma7*, *Psmb2*, *Psmb4*, *Psmb6*, *Psmc2*, *Psmc3*, *Psmc5*, *Psmd1*, and *Psmd11* (Supplementary Fig. S10A). Ingenuity pathway analysis (IPA) of differentially expressed genes demonstrated that several molecular networks were upregulated in Atg7^iSkM-KO^ mice as compared to Atg7^f/f^ mice in response to sepsis (Supplementary File S2.1). This indicates that strong activation of the ubiquitin proteasome proteolytic pathway is triggered in muscles in which the essential autophagy gene *Atg7* has been deleted, and that this activation may lead to more severe atrophy than when it has not been deleted.

**Figure 7:**
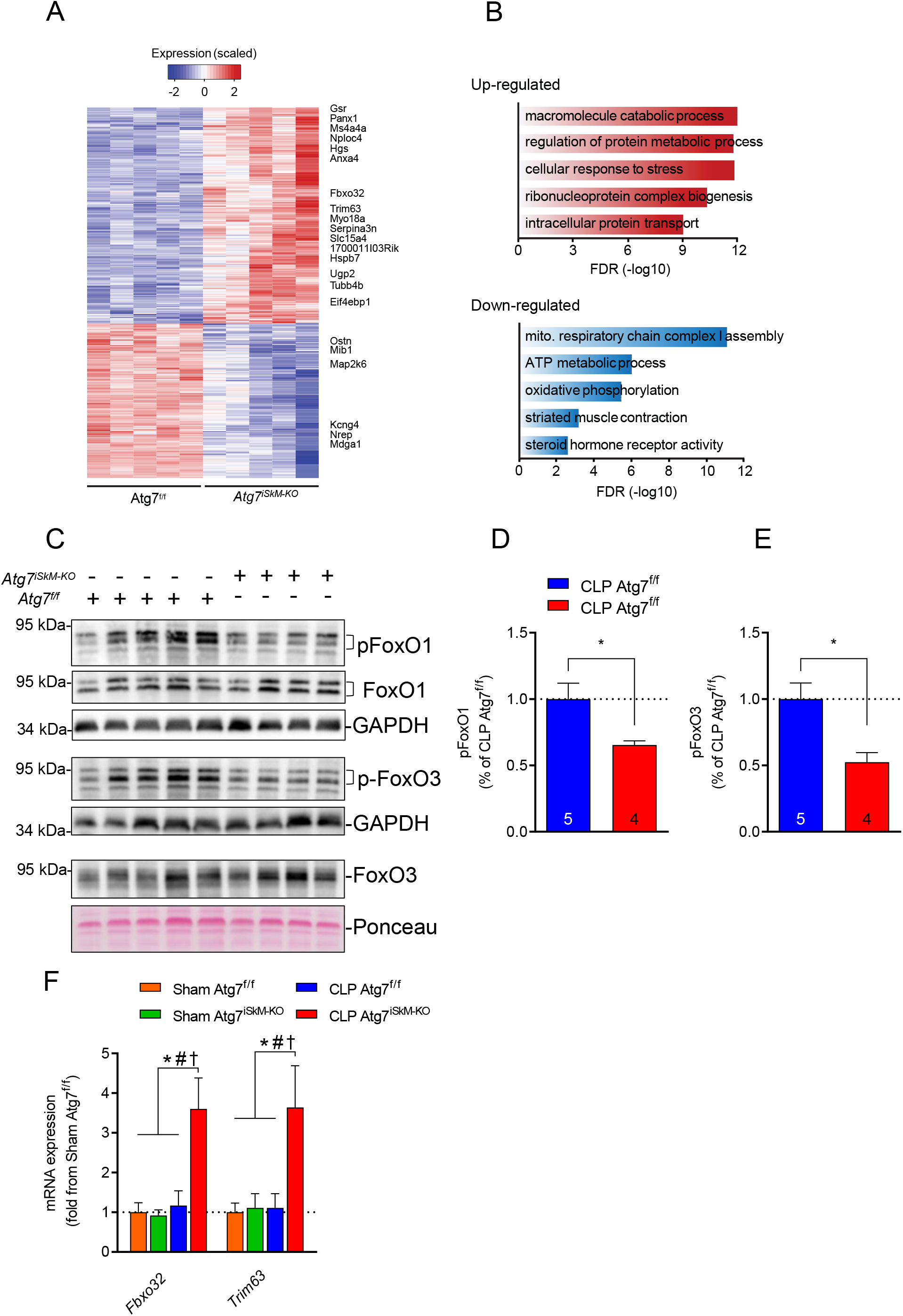
Effects of prolonged sepsis on muscle transcriptomes. A) Heatmap showing skeletal muscle gene expression signatures in Atg7^f/f^ and Atg7^iSkM-KO^ mice in response to prolonged sepsis. Colors indicate relative expression levels; red indicates high expression and blue indicates low expression. B) Top five upregulated and downregulated pathways as identified through biometric analyses. FDR=false discovery ratio. C) Representative immunoblots of pFoxO1, FoxO1, pFoxO3, and FoxO3 protein levels in skeletal muscles of female Atg7^f/f^ and Atg7^iSkM-KO^ mice 6 days after CLP showing decreased phosphorylation of FoxO1 and FoxO3 resulting from Atg7 knockout and sepsis. D-E) Densitometric analyses of pFoxO1 and pFoxO3 proteins normalized to GAPDH in Atg7^f/f^ and Atg7^iSkM-KO^ mice 6 days after CLP. GAPDH or Ponceau serves as loading control. Number of animals indicated within bars. Data presented as mean ± SEM. Knockout effect indicated by *p < 0.05, CLP Atg7^f/f^ vs. CLP Atg7^iSkM-KO^. (F) mRNA expressions of E3 ubiquitin ligases in TA muscles of female Atg7^f/f^ and Atg7^iSkM-KO^ mice 6 days after sham surgery or CLP. Data presented as fold change relative to sham Atg7^f/f^ and as mean ± SEM, n = 5-8 per group. * p <0.05 vs sham Atg7^f/f^; # p <0.05 for sepsis effect (i.e. sham Atg7^f/f^ vs. CLP Atg7^f/f^ or sham Atg7^iSkM-KO^ vs. CLP Atg7^iSkM-KO^); † p < 0.05 for sepsis plus knockout effect (i.e. CLP Atg7^f/f^ vs. CLP Atg7^iSkM-KO^).

FoxO transcription factors are major regulators of the autophagy and proteasome proteolytic pathways (18, 30). They transcribe several autophagy-related genes, ubiquitin E3 ligases, and proteasome subunits. To determine the degree to which they were activated in response to autophagy inactivation and sepsis, FOXO1 and FOXO3 phosphorylation levels were quantified in the TA muscles of Atg7^f/f^ and Atg7^iSkM-KO^ mice. In response to sepsis, the phosphorylation of both transcription factors was strongly attenuated in Atg7^iSkM-KO^ mice as compared to Atg7^f/f^ mice (Fig. 7C-E). *Fbxo32* and *Trim63* were also induced in Atg7^iSkM-KO^ mice, but not in Atg7^f/f^ mice, where levels remained like those detected in sham mice (Fig. 7F). These results show that the ubiquitin proteasome pathway is sustainably activated in response to prolonged sepsis when autophagy has been inactivated.

Transcriptome analyses showed that several cellular processes, such as mitochondrial respiratory chain complex assembly, ATP metabolism, oxidative phosphorylation, muscle contraction, and steroid hormone receptor activity, were downregulated in response to sepsis in the TA of Atg7^iSkM-KO^ mice as compared to septic Atg7^f/f^ mice (Fig. 7B). Downregulated genes associated with mitochondrial metabolism included NADH: ubiquinone oxidoreductase subunits S8, S5, S4, B7, B2, and B11 (*Ndufs8*, *Ndufs5*, *Ndufs4*, *Ndufb7*, *Ndufb2,* and *Ndufb11*), NADP: ubiquinone oxidoreductase complex assembly factors 6, 5, 9, 13, 11, and 1 (*Ndufaf6*, *Ndufaf5*, *Ndufa9*, *Ndufa13*, *Ndufa11*, and *Ndufa1*) (Supplementary Fig. 10B). A full list of differentially regulated gene and pathway analyses are listed in Dataset S2.1. Collectively, these results suggest that inactivation of autophagy intensifies the negative repercussions that sepsis exerts on a number of processes that are critical to skeletal muscle contractile function, as well as enhancing activation of the proteasome proteolytic pathway, resulting in worsened muscle fiber atrophy.

### Impact of sepsis and muscle-specific inactivation of autophagy on whole-body metabolism & muscle autophagy

To evaluate the regulatory role of skeletal muscle autophagy in relation to whole body metabolism, indirect calorimetry was performed before and after surgery. Before surgery, no differences in respiratory exchange ratio (RER), rate of oxygen consumption (VO2), heat production, feeding behavior, non-fasting blood glucose, body temperature, and whole body fatty acid oxidation were found between Atg7^f/f^ and Atg7^iSkM-KO^ mice, despite the latter having lower total movement (Supplementary Fig. S11). On Day 1 post-surgery, total VO2 decreased in response to sepsis in Atg7^f/f^ and Atg7^iSkM-KO^ mice, and recovered to pre-surgical values thereafter (Fig. 8A).

**Figure 8:**
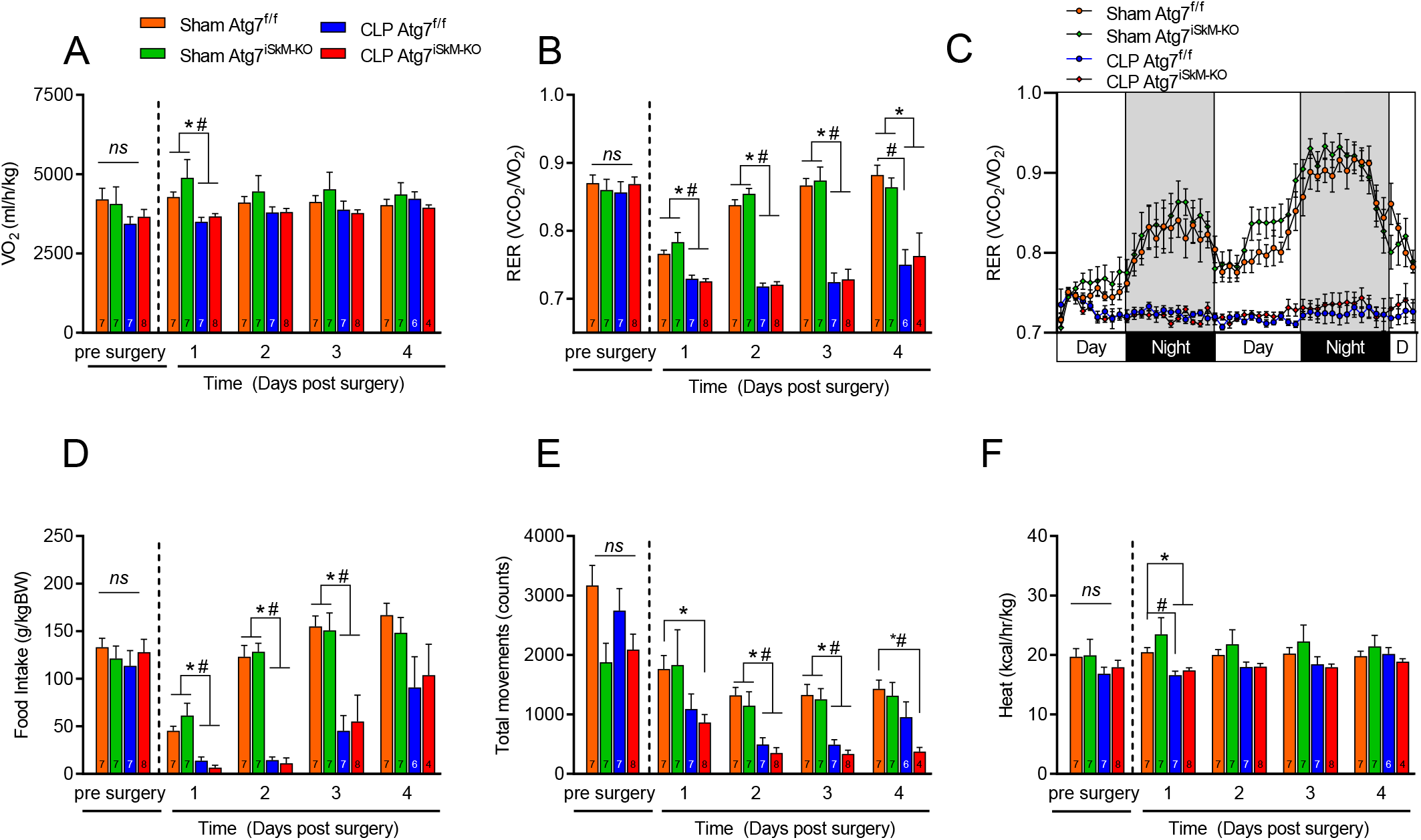
Effects of autophagy inactivation and sepsis on metabolism. A-C) Average kinetic data for whole-body oxygen consumption (VO_2_) and RER (VCO_2_/VO_2_) during day and night periods over a 48-h period after sham surgery or CLP. D) Food intake (gram of consumed food per kg of body mass per day. (E) Total movement (ambulatory activity). (F) Heat (energy expenditure) relative to daily body mass. Metabolic parameters measured by indirect calorimetry for 2 days prior to surgery and 4 days after sham surgery or CLP. Food intake and metabolic parameters expressed relative to daily body mass, given differences in body composition over time. PhenoMaster metabolic cage data measured on a dark (6:00 p.m. to 6:00 a.m.)/light (6:00 a.m. to 6:00 p.m.) cycle. Data presented as mean ± SEM. * p <0.05 vs sham Atg7^f/f^; # p <0.05 for sepsis effect (i.e. sham Atg7^f/f^ vs. CLP Atg7^f/f^ or sham Atg7^iSkM-KO^ vs. CLP Atg7^iSkM-KO^); † p < 0.05 for sepsis plus knockout effect (i.e. CLP Atg7^f/f^ vs. CLP Atg7^iSkM-KO^).

RER in sham Atg7^f/f^ and Atg7^iSkM-KO^ mice declined 1-day post-surgery, with complete recovery thereafter (Fig. 8B&C). These transient declines were coincident with decreased food intake (Fig. 8D). In response to sepsis, RERs in Atg7^f/f^ and Atg7^iSkM-KO^ mice persistently declined, suggesting increased dependence on lipid oxidation, which is consistent with reductions in food intake, although no differences in intake were observed (Fig. 8B-D). This suggests that the different rates of mass loss that occurred in these mice were not a consequence of variability in food intake (Fig. 2A&B). Persistent decrease in total movement was observed in both Atg7^f/f^ and Atg7^iSkM-KO^ mice in response to sepsis, an effect that was more severe in Atg7^iSkM-KO^ than Atg7^f/f^ mice (Fig. 8E). Heat production decreased in Atg7^f/f^ and Atg7^iSkM-KO^ mice in response to sepsis only on Day 1 post-surgery (Fig. 8F).

It has been well established that body temperature is a strong predictor of mortality during critical illness (31–33) and that skeletal muscles play a major role in thermogenesis (34, 35). To identify the mechanisms underlying mass loss and increased lethality of Atg7^iSkM-KO^ mice in response to sepsis, we monitored rectal temperature, blood glucose, and β-hydroxybutyrate levels. Autophagy ablation in sham operated mice had no impact on rectal temperature. While sepsis had a modest impact on rectal temperature in Atg7^f/f^ mice, it led to a progressive decline in rectal temperature in Atg7^iSkM-KO^ mice (Fig. 9A). Blood glucose levels in sham Atg7^f/f^ and Atg7^iSkM-KO^ mice declined on Day 1 post-surgery and remained lower than pre-surgery values for the duration of the experimental period (Fig. 9B). Blood glucose levels were lower in Atg7^f/f^ and Atg7^iSkM-KO^ mice in response to sepsis as compared to corresponding sham mice, and even lower in Atg7^iSkM-KO^ mice as compared to Atg7^f/f^ mice, up until Day 6 post-surgery (Fig. 9B). These findings are consistent with previous results demonstrating the importance of muscle autophagy in the regulation of glucose levels during fasting (36). Blood β-hydroxybutyrate levels were elevated in response to sepsis in both Atg7^f/f^ and Atg7^iSkM-KO^ mice, although to a greater degree in Atg7^iSkM-KO^ mice (Fig. 9C). These results suggest that increased ketogenesis compensates for hypoglycemia and that the absence of muscle autophagy worsens sepsis-induced hypoglycemia and hyperketonemia.

**Figure 9:**
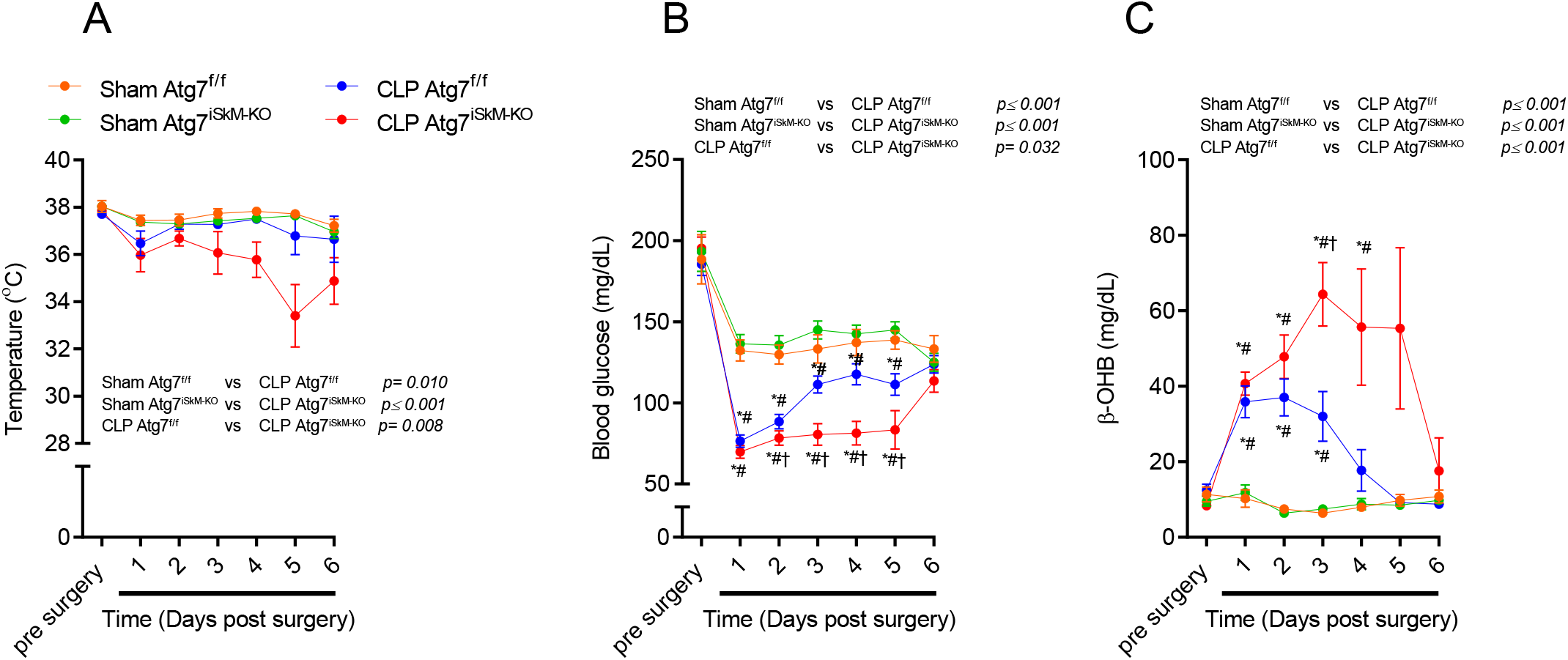
Sepsis exacerbates metabolic impairments in Atg7 knockout mice. A) Kinetics of core body temperature B) Whole blood glucose and C) β-OHB levels measured in Atg7^f/f^ and Atg7^iSkM-KO^ mice over a 6-day period between 6 and 8 a.m. Data presented as mean ± SEM. * p <0.05 vs sham Atg7^f/f^; # p <0.05 for sepsis effect (i.e. sham Atg7^f/f^ vs. CLP Atg7^f/f^ or sham Atg7^iSkM-KO^ vs. CLP Atg7^iSkM-KO^); † p < 0.05 for sepsis plus knockout effect (i.e. CLP Atg7^f/f^ vs. CLP Atg7^iSkM-KO^).

## Discussion

To date, the functional importance of autophagy in sepsis-induced skeletal muscle dysfunction has not been fully elucidated. In a recent report, we demonstrated that the molecular machinery of autophagy in skeletal muscle fibers is strongly activated by sepsis (15). In the current study, we used tamoxifen-inducible skeletal muscle-specific Atg7 knockout (Atg7^iSkM-KO^) mice to determine whether autophagy inactivation is protective of or detrimental to muscle function and atrophy in septic mice. We demonstrate for the first time how important autophagy is to the regulation of skeletal muscle integrity over the course of a critical illness like sepsis. We show that *Atg7* deletion selectively in skeletal muscles does not prevent skeletal muscle atrophy during sepsis. Rather, it accelerates losses of muscle mass. Another interesting observation is that *Atg7* deletion significantly lowers the specific force of septic muscles (force normalized to fiber cross-sectional area), indicating that in addition to the development of severe muscle atrophy, there is also more pronounced impairment in muscle contractility.

With the aim of identifying the mechanism underlying the more severe atrophy observed in Atg7^iSkM-KO^ mice in response to sepsis, we reasoned that the severity of sepsis-induced muscle atrophy might be explained by the upregulation of the proteasome proteolytic system since this system contributes to a large extent to the degradation of muscle proteins including myofilament proteins (37–40). Consistent with previous work from our laboratory which showed significant activation of the proteasome proteolytic pathway in septic skeletal muscles (15), we report that sepsis was associated with significant upregulation of two muscle-specific ubiquitin E3 ligases *Fbxo32* and *Trim63*. Interestingly, the degrees to which these two ubiquitin E3 ligases were induced 48 h post sepsis were similar in Atg7^iSkM-KO^ and Atg7^f/f^ mice suggesting that they don’t contribute to more severe muscle wasting observed at this time point of sepsis in Atg7^iSkM-KO^ mice. However, we should point out that six days post sepsis initiation, the expressions of both *Fbxo32* and *Trim63* in Atg7^f/f^ mice returned to normal levels while their expressions remained significantly elevated in septic Atg7^iSkM-KO^ mice. These findings can be partly explained by increased activities of FoxO1 and FoxO3 transcription factors, which are known to mediate the atrophy program and the induction of *Fbxo32* and *Trim63* in catabolic conditions (18, 30). In addition to ubiquitin E3 ligases, we measured the expression of constitutive and inducible subunits of 20S, 19S, and 11S components of the 26S proteasome as indicator of the activity of the proteasome proteolytic pathway. We found that the levels of two subunits of 11S (*Psme1* and *Psme2*) and one β subunit of 20S, *Psmb10*, were significantly elevated in muscles of Atg7^iSkM-KO^ mice as compared to Atg7^f/f^ mice under sham and sepsis conditions (Figure 5). These results indicate for the first time selective autophagy inactivation in normal (sham) skeletal muscles leads to compensatory increases in the expressions of various 26S proteasome subunits and that late phases of sepsis is associated substantial induction of both ubiquitin E3 ligases and proteasome subunit expression in muscles of mice with deficient autophagy.

Our results show that muscles of sham Atg7^iSkM-KO^ mice display a significant decrease in mitochondrial respiration and a trend for a decrease in the time to mPTP opening. These results therefore strengthen the literature showing that autophagy is essential for the maintenance of optimal mitochondrial function (24–28, 41). In agreement with previous studies (22, 42–45), we also report that sepsis results in mitochondrial dysfunction, characterized by reduced respiration and impaired mPTP function. The results suggest that an accumulation of dysfunctional mitochondria might have contributed to sepsis-induced muscle atrophy and weakness in Atg7^f/f^ mice. In line with this view, we recently reported that overexpressing Parkin, a protein regulating mitophagy, can attenuate sepsis-induced skeletal muscle wasting (23). Surprisingly, however, mitochondrial respiration, time to mPTP opening and H_2_O_2_ emission were not further altered 48h post sepsis in Atg7^iSkM-KO^ mice. These data suggest that further accumulation of mitochondrial dysfunction might not have contributed to the more severe atrophy and contractile dysfunction in Atg7^iSkM-KO^ mice in response to sepsis. However, it should be noted that many mitochondrial genes were still down regulated 6 days post CLP in Atg7^iSkM-KO^ mice, suggesting that further mitochondrial dysfunction might progressively develop past 48h in autophagy deficient skeletal muscle.

Another important finding of our study is that sepsis was associated with stronger signs of sickness in Atg7^iSkM-KO^ mice relative to Atg7^f/f^ mice (Fig. 2D). Indeed, Atg7^f/f^ mice exhibited several sickness behaviors in response to sepsis, including loss of appetite, reduced physical activity, and severe body mass loss, resulting in a reduced survival rates (e.g., 65% survival six days post CLP). These behavioral abnormalities associated with sepsis have been previously reported by investigators using the CLP model of sepsis (46). However, a more severe sepsis phenotype was observed in Atg7^iSkM-KO^ mice, characterized by more pronounced body mass loss, more severe signs of sickness, and a worse survival rate. Importantly, differences in food intake are unlikely to explain the more severe phenotype seen in response to sepsis in Atg7^iSkM-KO^ mice since measurements of food intake showed no differences between the two genotypes. The more likely mechanism behind pronounced sepsis-induced body mass loss in Atg7^iSkM-KO^ mice is the significantly greater skeletal muscle wasting observed in these mice relative to Atg7^f/f^ mice.

We also observed that autophagy inactivation in skeletal muscle can worsen systemic metabolic derangements associated with sepsis, as reflected by more severe hyperketonemia and hypoglycemia. Ketogenesis is normally activated in response to catabolic stress to spare glucose and to support brain function (47). Because hyperketosis can lead to ketoacidosis, which can lead to coma and death, our data, and recent observations (48), suggest elevated β-hydroxybutyrate levels may be involved in the pathogenesis of sepsis. With respect to the more pronounced and persistent hypoglycemia that was seen in Atg7^iSkM-KO^ mice, several studies have demonstrated that autophagy plays an important role in the regulation of whole-body glucose metabolism (27, 36, 49). Indeed, Kim et al. recently showed that the muscle-selective deletion of Atg7 in high fat fed mice resulted in lower glycemia and protected against insulin resistance and obesity (27). In the liver, autophagy contributes to the maintenance of glycemia through the conversion of amino acids to glucose via gluconeogenesis (49). Indeed, in mice with liver-specific deletion of Atg7, starvation was shown to result in greater decrease in blood glucose levels as compared to wild type animal (49). Based on these findings, we speculate that induction of autophagy in septic muscles may be particularly important in providing the amino acids that the liver requires to maintain glycemia and in eliminating damaged proteins and organelles in muscle cells, thereby limiting contractile dysfunctions and optimizing survival.

It has been well established that skeletal muscles produce cytokines and chemokines under normal resting conditions and that this production increases in response to intense muscle contraction. Muscle cells are also known to express innate immune receptors including Toll-like receptors 2, 4, 5, and 9 (50). In the current study, we found that autophagy inactivation in sham Atg7^iSkM-KO^ mice was associated with significant increase in the expressions of *Tweak*, *IL-1β*, *Tnf-α R*, and *IL-18* relative to sham Atg7^f/f^ mice. These results suggest that autophagy exerts a negative influence on the production of pro-inflammatory cytokines in normal skeletal muscle fibers. This proposal is supported by several studies that documented negative effects of autophagy on innate immune responses and the production of pro-inflammatory cytokines in various cells. For instance, inactivation of autophagy in Becn1^+/-^ and Lcb3^-/-^ macrophages results in increased production of inflammasome-associated *IL-1β* and *IL-8* (51). This protective effect of autophagy was mediated by the removal of dysfunctional mitochondria and decreased cytosolic translocation of mitochondrial DNA which is known to stimulate the secretion of IL-1*β* and IL8. In an another study, Zhou et al. (52) reported that inactivation of autophagy in monocytes leads to activation of the pyrin domain containing 3 (NLRP3) inflammasome as a result of accumulation of damaged reactive oxygen species (ROS) generated by dysfunctional mitochondria. In the heart, deletion of *Becn1* causes an increase in the expression of pro-inflammatory cytokines *Tnf-α*, interleukin 17 alpha (*IL-17α*), *IL-6*, and interferon gamma (*IFNγ*) in LPS model of sepsis (53). Overexpression of *Becn1* in the heart elicited opposite effects on LPS-induced cytokine production (53). It is possible that the increase in muscle-derived *Tweak*, *IL-1β*, *Tnf-α R*, and *IL-18* mRNA levels in sham Atg7^iSkM-KO^ mice might have been due to accumulation of dysfunctional mitochondria and release of mitochondrial DNA into the cytosols of muscle fibers and the consequent activation of innate immune receptors.

It should be emphasized that our findings of a protective role of autophagy against sepsis-induced skeletal muscle atrophy and contractile dysfunction and increased mortality of septic Atg7^iSkM-KO^ mice relative to Atg7^f/f^ mice are in agreement with previous reports indicating a protective role for autophagy against sepsis-induced organ and cellular failure in the liver and immune cells. Indeed, selective inactivation of autophagy in the liver resulted in increased mortality, severe mitochondrial damage, and activation of apoptosis in murine sepsis models (54, 55). Similarly, selective inactivation of autophagy in T cells was associated with enhanced T cells apoptosis and increased sepsis-induced mortality (56). Since activation of autophagy in response to sepsis plays a pivotal role in the physiological maintenance of cellular homeostasis, our study raises concerns about early nutritional and insulin therapies that may suppress autophagy in critically ill patients.

In summary, we highlight in this study that autophagy inactivation in skeletal muscles triggers significant worsening of sepsis-induced muscle atrophy and contractile dysfunction. We also provide evidence indicating that autophagy inactivation in skeletal muscles worsens sepsis-induced hypoglycemia and hyperketosis and negatively impacts survival. These results position enhancing autophagy as a potential therapeutic approach for treating sepsis-induced skeletal muscle dysfunction and metabolic derangements.

## Methods

### Animals

All mice used in this study originated from a C57BL6/J genetic background. Atg7 floxed mice (57) were crossed with mice expressing Cre-ER^T2^ driven by human skeletal Actin (HSA) promoter (58). Resultant HSA-Cre-ER^T2^ Atg7^f/f^ mice were fed *ad libitum* with a certified rodent diet containing tamoxifen citrate (400 mg/kg) (Teklad TAM400/CreER, Envigo, Madison, WI) for 10 weeks (59), then returned to a standard chow diet (Teklad global 18% protein irradiated rodent diet 2918, Envigo) for 2-3 weeks to ensure elimination of tamoxifen from the serum prior to surgery (60). Mice fed this diet (Atg7^iSkM-KO^) were compared to sex and age-matched HSA-Cre-ER^T2^ Atg7^f/f^ mice that were fed a standard chow diet only. To control for the effects of tamoxifen diet *per se*, C57BL6/J mice (Charles River Laboratories, Saint-Constant, QC, Canada) fed according to the same tamoxifen protocol as described above were used as additional controls (Supplementary Fig. S1). All mice were group housed under a standard 12:12 h light/dark cycle at 22°C. All experiments were approved by the Research Ethics Board of the Research Institute of the McGill University Health Centre and are in accordance with the principles outlined by the Canadian Council of Animal Care.

### Cecal ligation and perforation

Cecal ligation and perforation (CLP) was performed to induce polymicrobial sepsis, as described previously (15), with minor modifications. The CLP model is known to accurately mimic the clinical features of human sepsis (61). Briefly, mice were anesthetized using ∼3% isoflurane with air (Piramal Critical Care, Bethlehem, PA) and a midline abdominal incision (∼2 cm) was performed. The caecum was carefully ligated at ∼1 cm from its distal end to control for the degree of disease severity (62, 63) then perforated via a through- and-through puncture with a 25^1/2^ gauge needle in a sterile environment, leaving 2 holes. The ligated caecum was gently compressed to extrude a small amount of cecal contents through the punctured holes, then replaced in the abdominal cavity. The peritoneum was closed in two separate layers using 3-0 absorbable polyfilament interrupted sutures and the skin was closed with a 9 mm surgical staple (AutoClip® System, Fine Science Tools, Foster City, CA). All animals received subcutaneous injections of Buprenex® (0.05 to 0.2 mg/kg buprenorphine in 1 ml of 0.9% saline) immediately after surgery and every 12 h thereafter to minimize pain (0.05 mg/kg in ∼100 µL of 0.9% saline). Sham-operated mice were subjected to identical procedures, except for cecal ligation and perforation. Age- and gender-matched controls were included for all procedures. Animals were returned to their cages (3-5 mice per cage) with free access to food and water, unless stated otherwise. All animals were closely monitored by animal care staff blinded to surgical status and genotype for signs of excessive pain or distress, such as lack of movement agonized breathing, or excessive body mass loss (20%). Any mouse determined to be moribund was euthanized. Several mice were sacrificed 48 h post-surgery for use in muscle contractility and mitochondrial function tests. Mice that were used for 6-day measurements were monitored more intensively, at least 3-5 times daily. To minimize suffering, temperature was measured daily using a rectal probe. If body temperature dropped more than 4°C, animals were euthanized. A daily clinical severity score (CSS) was assigned to track the course of the disease and to minimize excessive pain or distress, as previously described (64). Grade 1 was interpreted as no signs of illness, grade 2 as mild signs (e.g. less active, slightly hunched posture), grade 3 as medium severe signs (e.g. less active, hunched, and slow movement), and grade 4 as very severe signs (e.g. lethargic, severely hunched, no movement, agonal breathing, and moribund). Mice were euthanized with CO_2_ and death was confirmed by cervical dislocation.

### *In situ* measurement of TA contractile function

To determine skeletal muscle contractile function, animals were anesthetized with an intraperitoneal injection of a ketamine-xylazine cocktail (ketamine: 130 mg/kg; xylazine: 20 mg/kg). Anesthesia was maintained with 0.05 ml supplemental doses, as needed. A Dynamic Muscle Control and Analysis Software (DMC/DMA) Suite was used for collection and data analysis (Aurora Scientific, Aurora, ON). The distal tendon of the left TA muscle was isolated and attached to the arm of a 305C-LR dual-mode muscle lever with 4.0 surgical silk, as previously described, with minor modifications (65–67). The partially exposed muscle surface was kept moist and directly stimulated with an electrode placed on the belly of the muscle. Optimal muscle length and voltage was progressively adjusted to produce maximal tension and length was measured with a microcaliper. The pulse duration was set to 0.2 ms for all tetanic contractions. Force-frequency relationship curves were determined at muscle optimal length at 10, 30, 50, 70, 100, 120, 150 and 200Hz, with 1 min intervals between stimulations to avoid fatigue. Following tetanic-force measurements, the muscle was rested for 2 min then subjected to a fatigue resistance protocol of 60 tetanic contractions (75 Hz/200 ms) every 2 s for a total of 2 min. At the end of each experiment, mice were sacrificed, and muscles were carefully dissected, weighed, and frozen in liquid nitrogen or in isopentane pre-cooled in liquid nitrogen. *In situ* muscle force was normalized to tissue cross-sectional area (expressed as Newtons/cm^2^). Muscle cross-sectional area was estimated by dividing muscle mass by the product of muscle length and muscle density (1.056 g/cm^3^).

### *In situ* measurement of mitochondrial function

#### Preparation of permeabilized muscle fibers

Mitochondrial function was determined in freshly excised GAS muscles at 2-day post-surgery. Left GAS muscles were rapidly dissected and immersed in ice-cold (4°C) stabilizing buffer A (2.77 mM CaK2 EGTA, 7.23 mM K2 EGTA, 6.56 mM MgCl2, 0.5 mM dithiothreitol (DTT), 50 mM 2-(N-morpholino) ethanesulfonic acid potassium salt (K-MES), 20 mM imidazol, 20 mM taurine, 5.3 mM Na2 ATP, and 15 mM phosphocreatine, pH 7.3). Different regions of the GAS muscle have different muscle fiber type composition and the following protocol was used to minimize variance in results. Mitochondrial respiration and H_2_O_2_ emission were performed in the oxidative red GAS while the assessment of the mitochondrial permeability transition pore sensitivity to Ca^2+^ was assessed in the glycolytic white GAS. GAS samples were weighed then separated into small fiber bundles using fine forceps and a Leica S4 E surgical dissecting microscope (Leica Microsystems, Wetzlar, Germany). Muscle fiber bundles were incubated for 30 min at low rocking speed in glass scintillation vials containing buffer A supplemented with 0.05 mg/mL saponin to selectively permeabilize the sarcolemma. Red fiber bundles used for respiration and H_2_O_2_ emission analyses were then washed 3 times for 10 min at low rocking speed in buffer Z (110 mM, 35 mM KCI, 1 mM EGTA, 3 mM MgCl_2_, and 10 mM K_2_HPO_4_, pH 7.3) at 4°C, supplemented with 5 mg/mL BSA. White fiber bundles used for CRC were washed 3 times for 10 min at low rocking speed in buffer C (80 mM K-MES, 50 mM HEPES, 20 mM taurine, 0.5 mM DTT, 10 mM MgCl2, and 10 mM ATP, pH 7.3) at 4°C. Bundles were then transferred into buffer D (800 mM KCl, 50 mM HEPES, 20 mM taurine, 0.5 mM DTT, 10 mM MgCl_2_, and 10 mM ATP, pH 7.3) for 30 min at 4°C. CRC bundles were then washed 3 times in a low-EGTA CRC buffer (250 mM sucrose, 10 mM Tris, 5 µM EGTA, and 10 mM 3-(*N-morpholino*) propane sulfonic acid (MOPS, pH 7.3) at 4°C and kept on ice until measurements were performed.

#### Mitochondrial respiration

Mitochondrial respiration was measured at 37°C in 2 mL of buffer Z using an Oxygraph O2K system (Oroboros Instruments, Innsbruck, AT). Briefly, 3 to 6 mg (wet mass) of GAS bundled fibers were weighed and added to the respiration chamber. The following substrates were added sequentially: 10 mM glutamate, 5 mM malate (G+M), and 2 mM ADP. Respiration rate was normalized as nanomoles per min per mg of wet muscle mass.

#### Mitochondrial H_2_O_2_ production

H_2_O_2_ production was measured using an Amplex Red/horseradish peroxidase (HRP) assay, a Hitachi FL-2500 fluorescence spectrophotometer (excitation/emission wavelength of 563/587 nm), and FL Solutions software. Following a period of baseline autofluorescence, 4–6 mg (wet mass) of GAS bundled fibers were weighed and inserted into a thermojacketed, magnetically stirred cuvette containing 600 μL of buffer Z, Amplex Red (5.5 μM), and HRP (1 U/ml) at 37°C. The following substrates and inhibitors were added sequentially: glutamate (5 mM) and malate (5 mM), succinate (5 mM), ADP (0.1 mM), ADP (1 mM) and AA (400μM). H_2_O_2_ production was normalized as picomoles per min per mg of wet mass. All experiments were analyzed with a custom-made software program developed using IGOR Pro software (Wavemetrics, Portland, OR).

#### Mitochondrial Ca^2+^ retention capacity (CRC)

Sensitivity to mitochondrial permeability transition pore (mPTP) opening was evaluated by determining the CRC of mitochondria. GAS bundled fibers were placed in a cuvette with 600 μL of CRC assay mix (10 mM Pi, 2.5 mM malate, 5 mM glutamate, 0.5 nM oligomycin, 1 μM calcium green) at 37°C. Fluorescence was measured using a Hitachi F2500 spectrophotometer and FL Solutions software with excitation and emission detectors set at 505 and 535 nm and progressive uptake of Ca^2+^. Ca^2+^ uptake and release were recorded over 5-10 min. The time to pore opening was analyzed as the total amount of Ca^2+^ intake by the mitochondria before Ca^2+^ release. All experiments were analyzed with a custom-made software program developed using IGOR Pro software (Wavemetrics, OR).

#### Transmission electron microscopy

Small strips prepared from GAS fibers were fixed in 2% glutaraldehyde buffer solution in 0.1 M cacodylate, pH 7.4, then post-fixed in 1% osmium tetroxide in 0.1 M cacodylate buffer. Tissues were dehydrated using a gradient of increasing concentrations of methanol to propylene oxide and infiltrated and embedded in EPON^TM^ at the Facility for Electron Microscopy Research at McGill University. Ultrathin longitudinal sections (60 nm) were cut with a Reichert-Jung Ultracut III ultramicrotome (Leica Microsystems), mounted on nickel carbon-formvar coated grids, and stained with uranyl acetate and lead citrate. Sections were imaged using a FEI Tecnai 12 transmission electron microscope at 120 kV and images were digitally captured using an AMT XR80C CCD camera system.

#### Measurement of myofiber atrophy

TA samples were mounted in tragacanth (Sigma-Aldrich # G1128) on plastic blocks and frozen in liquid isopentane cooled in liquid nitrogen and stored at −80°C. Samples were cut into 10µm cross-sections using a cryostat at −20°C then mounted on lysine coated slides (Superfrost), as described in (65, 68). Cross-sections were brought to room temperature, rehydrated with PBS (pH 7.2), then blocked with goat serum (10% in PBS). They were then incubated with primary polyclonal anti-laminin rabbit IgG antibody (Sigma-Aldrich # L9393) for 1 h at room temperature. Sections were then washed three times in PBS before being incubated for 1 h at room temperature with an Alexa Fluor® 594 goat anti-rabbit IgG antibody (A-11037, Invitrogen, Waltham, MA). Sections were then washed three times in PBS and slides were cover slipped using Prolong™ Gold (P36930, Invitrogen) as a mounting medium. Slides were imaged with a Zeiss fluorescence microscope (Zeiss Axio Imager 2). Median distribution of minimum Feret diameters of at least 300 fibers per muscle sample were analyzed using ImageJ (NIH, Bethesda, MD) (69). The degree of myofiber atrophy was calculated as percent difference in mean minimum Feret diameters, relative to muscles of sham Atg7^f/f^ mice. The means, fibers distributions, and average number of fibers analyzed per muscle per mouse are shown in Supplemental Figure 6.

#### p62/SQSTM1 positive myofibers

Sections were stained to assess p62 positive fibers as previously described (70). Muscle sections were first allowed to reach room temperature and rehydrated with PBS (pH 7.2) and then fixed in 4% PFA, permeabilized in 0.1% Triton X-100 in PBS for 15min. Slides were then washed three additional times with PBS and then blocked with goat serum (10% in PBS) for 1 hour. After, muscle sections were incubated for 1 hour at room temperature with anti-mouse p62/SQSTM1 (Novus Biologicals Inc. clone 2C11, 1:200). Sections were then washed three times in PBS before being incubated for 1 h at room temperature with an Alexa Fluor® 488 goat anti-rabbit IgG antibody (A-21131, 1:500). Myofibers positive for p62 were quantified as a percentage of the total number of fibers counted.

#### Fiber type composition

TA muscle sections were immunolabeled for myosin heavy chain (MHC) types I, IIa and IIb, as previously described (68). Cross-sections were brought to room temperature, rehydrated with PBS (pH 7.2), then blocked with goat serum (10% in PBS). They were then incubated for 1 h at room temperature with the following primary antibody cocktail: mouse IgG2b monoclonal anti-myosin heavy chain (MHC) type I (BA-F8), mouse IgG1 monoclonal anti-MHC type IIa (SC-71), mouse IgM monoclonal anti-MHC type IIb (BF-F3, 1:200), and rabbit IgG polyclonal anti-Laminin (Sigma-Aldrich # L9393). Sections were then washed three times in PBS before being incubated for 1 h at room temperature with the following Alexa Fluor® secondary antibody cocktail: Alexa Fluor® 350 goat anti-mouse IgG2b (y2b) (A-21140), Alexa Fluor® 594 goat anti-mouse IgG1 (y1) (A-21125), Alexa Fluor® 488 goat anti-mouse IgM (A-21042,1:500), and Alexa Fluor® 488 goat anti-rabbit IgG (A-11008). They were then washed three times in PBS and slides were cover slipped using Prolong™ Gold (P36930) as a mounting medium. Images were captured using a Zeiss Axio Imager M2. All MHC-targeting primary antibodies were purchased from the Developmental Studies Hybridoma Bank (DSHB) at the University of Iowa. Fiber type analyses were measured by a single observer blinded to sample identity.

### Physiological measurements

#### Body composition analysis

Baseline (prior to surgery) body mass were recorded and thereafter at various timepoints post-surgery. The body mass loss for each animal was calculated as percent difference relative to baseline body mass. In a separate experiment, lean and fat mass were measured in live animals without anesthesia or sedation using an EchoMRI™ (Model ET-040), prior to surgery and 4 days post-surgery. Lean and fat mass loss for each animal was calculated as percent difference relative to baseline values.

#### Grip strength

Forelimb grip strength was measured 3 days prior to surgery using a grip strength meter with a metal grid (Bioseb, Vitrolles, FR). Mice were held at the base of the tail then placed slightly above the grid to enable the animal to grab hold of it. They were then gently pulled away until the grid was released. Maximum force in grams exerted by each mouse was recorded 3 times with a 3-minute interval between each measurement, to prevent fatigue. Data are expressed as average grams of force divided by body mass. All measurements were performed by an investigator blinded to mouse genotype.

#### Blood metabolic parameters

Whole blood glucose and β-hydroxybutyrate (β-OHB) levels were measured between 6 and 8 a.m. in tail vein blood at various time points during the study with an Accu-Chek handheld glucometer (Roche Diagnostics, Indianapolis, IN) or a Freestyle Optium Neo ketometer (Abbott Laboratories. Saint-Laurent, QC). Data are expressed in mg/dL.

#### Body temperature

Internal body temperature (°C) was measured between 6 and 8 a.m. at various time points during the study with a Thermalert TH-5 thermometer with a flexible rectal probe (Physitemp Instruments, Clifton, NJ).

#### Whole animal metabolism, food intake, and locomotor activity

Indirect calorimetry was performed at the McGill Mouse Metabolic Platform. Animals were individually housed at room temperature (22°C-24°C) in PhenoMaster metabolic cages (TSE Systems, Chesterfield, MO). After 3 days of acclimatization, O_2_ consumption, CO_2_ production, energy expenditure (heat), respiratory exchange ratio (RER), locomotor activity (beam breaks), and caloric intake were measured. Measurements were made 2 days prior to and for 4 consecutive days after sham surgery or CLP. Average daily metabolic parameters were all normalized to daily body mass. Locomotor activity was simultaneously monitored using infrared sensor frames. Energy expenditure was calculated as (VO_2_*[3.815 + (1.232*RQ)]*4.1868). RER was calculated as CO_2_ production divided by O_2_ consumption.

### Autophagic flux quantification

Autophagic flux was monitored in 14–16 weeks old male wild-type C57BL6/J mice subjected to sham or CLP procedures. Briefly, mice received an i.p. injection of the autophagy inhibitor leupeptin (40 mg/kg dissolved in sterile PBS (Sigma-Aldrich # L2884) 20 h post sham or CLP procedures. The sham and CLP control groups received an equivalent volume of sterile PBS. Mice were euthanized 4 hours post leupeptin injection (24h post sham or CLP surgery). Muscles were then rapidly excised and frozen in liquid nitrogen and used for immunoblotting to detect LC3B proteins. Data are shown in Supplemental Figure 3E-F.

### Cytokines and myokine quantification

#### IL-1 β ELISA

The concentration of IL-1β in GAS muscle extracts at 2-day post-surgery was measured using Mouse IL-1 β ELISA Kit (Invitrogen #88-7013), according to manufacturer’s instructions. Approximately 15-30 mg of muscle tissue were homogenized in lysis buffer (50mM Tris base, 150mM NaCl, 1%, Triton X-100, 0.5% sodium deoxycholate, 0.1% SDS and 10μl/ml of a protease inhibitor cocktail (Sigma-Aldrich # P8340). The homogenate was centrifuged at 15,000g for 15 minutes at 4°C, and the supernatant was collected. The protein concentration was measured using the Bradford method. Data are expressed in pg/mg of protein.

#### Multiplex analysis

The concentrations of TNFα and FGF21 in GAS muscle extracts at 2-day post-surgery were measured using MILLIPLEX MAP Kit, Mouse Aging Magnetic Bead Panel-1 (Millipore MAGE1MAG-25K), according to manufacturer instructions. Approximately 15-30 mg of muscle tissue was homogenized in PBS based lysis buffer (0.1% Triton X-100, 10μl/ml of a protease inhibitor cocktail (Sigma-Aldrich # P8340)). The homogenate was centrifuged at 15,000g for 15 minutes at 4°C, and the supernatant was collected. The protein concentration was measured using the Bradford method. Data are expressed in pg/mg of protein.

#### *In vivo* protein synthesis measurements

*In vivo* protein synthesis in skeletal muscle was measured using the SUnSET technique (29). Briefly, mice were weighed and injected with intraperitoneally puromycin dissolved in 100 μl of sterile PBS ((0.04 μmol puromycin/ g body mass (Sigma-Aldrich # P8833)). At 30 min post puromycin injection, TA muscles were carefully removed and then frozen in liquid nitrogen and stored (−80 °C) for immunoblotting analysis using a mouse IgG2a monoclonal anti-puromycin antibody (clone 12D10, 1:2500; Millipore # MABE343).

#### Immunoblotting

Portions of frozen muscle samples (≈15-30 mg) were homogenized in an ice-cold lysis buffer (50mM HEPES, 150mM NaCl, 100mM NaF, 5mM EDTA, 0.5% Triton X-100, 0.1 mM DTT, 2 µg/ml leupeptin, 100 µg/ml PMSF, 2 µg/ml aprotinin, and 1 mg/100 ml pepstatin A, pH 7.2) with ceramic beads using Mini-Beadbeater (BioSpec Products, Bartlesville, OK) at 60Hz. Muscle homogenates were kept on ice for 30 min with periodic agitation then centrifuged at 5000 *g* for 15 min at 4°C. Supernatants were collected and pellets were discarded. The protein content in each sample was determined using the Bradford method. Aliquots of crude muscle homogenate were mixed with Laemmli (6X, SDS-sample buffer, reducing, BP-111R) buffer (Boston BioProducts, Ashland, MA) and subsequently denatured for 5 min at 95°C. Equal or equivalent amounts of protein extracts (30 ug per lane) were separated by SDS-PAGE, then transferred onto polyvinylidene difluoride (PVDF) membranes (Bio-Rad Laboratories, Saint-Laurent, QC) using a wet transfer technique. Total protein was detected with Ponceau-S solution (Sigma-Aldrich, #P3504) or with stain-free technology (Bio-Rad). Membranes were blocked in PBS + 1% Tween® 20 + 5% BSA for 1 h at room temperature then incubated overnight with specific primary antibodies at 4°C.Membranes were washed in PBST (3×5 min) then incubated with HRP-conjugated goat anti-rabbit (ab6721) or rabbit anti-mouse (ab6728) secondary antibodies (Abcam, Cambridge, MA) for 1 h at room temperature, before further washing. Immunoreactivity was detected using Pierce™ enhanced chemiluminescence substrate (Thermo Fisher Scientific, Waltham, MA), a ChemiDoc™ Imaging System, and ImageLab software (Bio-Rad). Optical densities (OD) of protein bands were normalized to a loading control. Immunoblot data are presented relative to sham Atg7^f/f^.

#### RNA extraction and real-time PCR

Total RNA was extracted from frozen muscle samples using a PureLink™ RNA Mini Kit (Invitrogen Canada, Burlington, ON). Quantification and purity of RNA was assessed using the A260/A280 absorption method. Total RNA (2 μg) was reverse transcribed using a Superscript II® Reverse Transcriptase Kit and random primers (Invitrogen Canada, Burlington, ON). Reactions were incubated at 42°C for 50 min and at 90°C for 5 min. Real-time PCR detection of mRNA expression was performed using a 7500 Sequence Detection System (Applied Biosystems, Foster City, CA). Specific primers (10 μM) were designed to quantify various mRNAs (Table 1). 18S were used as endogenous control transcripts. Each primer (3.5 μl) was combined with reverse-transcriptase reagent (1 μl) and SYBR® Green master mix (25 μl) (Qiagen, Valencia, CA). The thermal profile was as follows: 95°C for 10 min; 40 cycles each of 95°C for 15 s; 57°C for 30 s; and 72°C for 33 s. For each target gene, cycle threshold (C_T_) values were obtained. Relative mRNA level quantifications of target genes were determined using the threshold cycle (ΔΔC_T_) method.

#### Microarrays

Total RNA was isolated from TA muscle samples (20-25 mg) that were obtained 6 days post-surgery. A total of 21,981 mouse genes were included in the Affymetrix Mouse Clariom S Assay (Affymetrix, Santa Carla, CA) and all steps were performed at the McGill University and Génome Québec Innovation Centre (Montréal, QC). Raw data were pre-processed using Transcriptome Analysis Console (TAC) 4.0.1 software. A limma R software package (71) was used to identify expression level differences between Atg7^f/f^ and Atg7^iSkM-KO^ muscles. Nominal p-values were corrected for multiple testing using the Benjamini-Hochberg method. Genes that showed an FDR of <0.05 were considered statistically significant and differentially expressed (DE). The Canadian Centre for Computational Genomics (C3G) assisted with bioinformatic analyses. Gene Ontology (GO) was used to broadly identify annotation categories of significantly affected molecular function and biological processes.

#### Data Analysis and Statistics

GraphPad Prism software version 8.0 was used for statistical analyses, unless otherwise indicated. Data are reported as mean ± SEM. Differences between experimental groups were analyzed with one-way analysis of variance (ANOVA) or two-way ANOVA with corrections for multiple comparisons using the Benjamini, Krieger, and Yekutili two-stage method for controlling for the false discovery rate q<0.05 and p<0.05 were considered statistically significant). Comparisons of survival curves were performed using both Mantel-Cox and Gehan-Breslow-Wilcoxon tests. *P* values of < 0.05 were considered statistically significant. The exact number of animals in each experiment is indicated in figure legends.

## Abbreviations

Atg7: Autophagy related 7
Bnip3: BCL2 interacting protein 3
Bnip3L: BCL2 interacting protein 3 like
CLP: Cecal ligation and perforation
Cre: Cre recombinase
ER^T2^: Triple mutant form of human estrogen receptor
Fn14: TNF receptor superfamily member 12A
Fgf21: Fibroblast growth factor 21
FOXO1: Forkhead box O1
FOXO3: Forkhead box O3
Gabarapl1: GABA type A receptor associated protein like 1
GAS: Gastrocnemius
LC3B: Microtubule associated protein 1 light chain 3 beta
IL-1b: Interleukin 1
IL-18: Interleukin 18
IL-24: Interleukin 24
mPTP: mitochondria permeability transition pore
Park2: Parkin RBR E3 ubiquitin protein ligase
PCR: Polymerase chain reaction
RER: Respiratory exchange ratio
Sod1: Superoxide dismutase 1
Sod2: Superoxide dismutase 2
Sqstm1: Sequestome 1
Tweak: TNF superfamily member 12
TA: Tibialis anterior
Tnf-α: Tumor necrosis alpha
Tnf-αR: Tumor necrosis alpha receptor

## Acknowledgements

The authors are grateful to Ms Anne Gatensby for editing the manuscript and to Drs. M. Kokoeva and Xiaohong Liu for their assistance in using the Mouse Metabolic Phenotyping Platform, Research Institute of McGill University Health Centre. We also thank Génome Québec Innovation Centre for generating microarray data and Alain Sarabia Pacis (Canadian Center for Computational Genomics (C3G) Montréal Node, McGill University, Montréal, QC, Canada) for his bioinformatics support. The C3G is supported by the Canadian government through Génome Canada. We finally thank Jeannie Mui at the Facility for Electron Microscopy Research (FEMR, McGill University, Montréal, QC, Canada) for her support and expertise.

## References

1. Khan J, Harrison TB, Rich MM, and Moss M. Early development of critical illness myopathy and neuropathy in patients with severe sepsis. Neurology. 2006;67(8):1421–5.

2. Tennila A, Salmi T, Pettila V, Roine RO, Varpula T, and Takkunen O. Early signs of critical illness polyneuropathy in ICU patients with systemic inflammatory response syndrome or sepsis. Intensive Care Med. 2000;26(9):1360–3.

3. De Jonghe B, Sharshar T, Hopkinson N, and Outin H. Paresis following mechanical ventilation. Curr Opin Crit Care. 2004;10(1):47–52.

4. de Jonghe B, Lacherade JC, Sharshar T, and Outin H. Intensive care unit-acquired weakness: risk factors and prevention. Crit Care Med. 2009;37(10 Suppl):S309-15.

5. Cheung AM, Tansey CM, Tomlinson G, Diaz-Granados N, Matté A, Barr A, et al. Two-Year Outcomes, Health Care Use, and Costs of Survivors of Acute Respiratory Distress Syndrome. American journal of respiratory and critical care medicine. 2006;174(5):538–44.

6. Tiao G, Fagan JM, Samuels N, James JH, Hudson K, Lieberman M, et al. Sepsis stimulates nonlysosomal, energy-dependent proteolysis and increases ubiquitin mRNA levels in rat skeletal muscle. J Clin Invest. 1994;94(6):2255–64.

7. Chai J, Wu Y, and Sheng ZZ. Role of ubiquitin-proteasome pathway in skeletal muscle wasting in rats with endotoxemia. Crit Care Med. 2003;31(6):1802–7.

8. Callahan LA, and Supinski GS. Sepsis-induced myopathy. Crit Care Med. 2009;37(10 Suppl):S354–67.

9. Supinski GS, and Callahan LA. Calpain activation contributes to endotoxin-induced diaphragmatic dysfunction. Am J Respir Cell Mol Biol. 2010;42(1):80–7.

10. Supinski GS, and Callahan LA. Caspase activation contributes to endotoxin-induced diaphragm weakness. J Appl Physiol (1985). 2006;100(6):1770-7.

11. Supinski GS, Wang W, and Callahan LA. Caspase and calpain activation both contribute to sepsis-induced diaphragmatic weakness. J Appl Physiol (1985). 2009;107(5):1389-96.

12. Hobler SC, Williams A, Fischer D, Wang JJ, Sun X, Fischer JE, et al. Activity and expression of the 20S proteasome are increased in skeletal muscle during sepsis. Am J Physiol. 1999;277(2):R434–40.

13. Tiao G, Lieberman M, Fischer JE, and Hasselgren PO. Intracellular regulation of protein degradation during sepsis is different in fast- and slow-twitch muscle. Am J Physiol. 1997;272(3 Pt 2):R849-56.

14. Hobler SC, Tiao G, Fischer JE, Monaco J, and Hasselgren PO. Sepsis-induced increase in muscle proteolysis is blocked by specific proteasome inhibitors. Am J Physiol. 1998;274(1):R30–7.

15. Stana F, Vujovic M, Mayaki D, Leduc-Gaudet JP, Leblanc P, Huck L, et al. Differential Regulation of the Autophagy and Proteasome Pathways in Skeletal Muscles in Sepsis. Crit Care Med. 2017;45(9):e971–e9.

16. Masiero E, Agatea L, Mammucari C, Blaauw B, Loro E, Komatsu M, et al. Autophagy is required to maintain muscle mass. Cell metabolism. 2009;10(6):507–15.

17. Sandri M. Autophagy in skeletal muscle. FEBS letters. 2010;584(7):1411–6.

18. Mammucari C, Milan G, Romanello V, Masiero E, Rudolf R, Del Piccolo P, et al. FoxO3 controls autophagy in skeletal muscle in vivo. Cell Metab. 2007;6(6):458–71.

19. Doyle A, Zhang G, Abdel Fattah EA, Eissa NT, and Li YP. Toll-like receptor 4 mediates lipopolysaccharide-induced muscle catabolism via coordinate activation of ubiquitin-proteasome and autophagy-lysosome pathways. FASEB J. 2011;25(1):99–110.

20. Jamart C, Gomes AV, Dewey S, Deldicque L, Raymackers JM, and Francaux M. Regulation of ubiquitin-proteasome and autophagy pathways after acute LPS and epoxomicin administration in mice. BMC Musculoskelet Disord. 2014;15:166.

21. Maes K, Stamiris A, Thomas D, Cielen N, Smuder A, Powers SK, et al. Effects of controlled mechanical ventilation on sepsis-induced diaphragm dysfunction in rats. Crit Care Med. 2014;42(12):e772–82.

22. Mofarrahi M, Sigala I, Guo Y, Godin R, Davis EC, Petrof B, et al. Autophagy and skeletal muscles in sepsis. PLoS One. 2012;7(10):e47265.

23. Leduc-Gaudet J-P, Mayaki D, Reynaud O, Broering FE, Chaffer TJ, Hussain SNA, et al. Parkin Overexpression Attenuates Sepsis-Induced Muscle Wasting. Cells. 2020;9(6):1454.

24. Carnio S, LoVerso F, Baraibar MA, Longa E, Khan MM, Maffei M, et al. Autophagy impairment in muscle induces neuromuscular junction degeneration and precocious aging. Cell Rep. 2014;8(5):1509–21.

25. Masiero E, Agatea L, Mammucari C, Blaauw B, Loro E, Komatsu M, et al. Autophagy is required to maintain muscle mass. Cell metabolism. 2009;10(6):507–15.

26. Wu JJ, Quijano C, Chen E, Liu H, Cao L, Fergusson MM, et al. Mitochondrial dysfunction and oxidative stress mediate the physiological impairment induced by the disruption of autophagy. Aging (Albany NY). 2009;1(4):425–37.

27. Kim KH, Jeong YT, Oh H, Kim SH, Cho JM, Kim YN, et al. Autophagy deficiency leads to protection from obesity and insulin resistance by inducing Fgf21 as a mitokine. Nat Med. 2013;19(1):83–92.

28. Lim JA, Zare H, Puertollano R, and Raben N. Atg5(flox)-Derived Autophagy-Deficient Model of Pompe Disease: Does It Tell the Whole Story? Molecular therapy Methods & clinical development. 2017;7:11–4.

29. Schmidt EK, Clavarino G, Ceppi M, and Pierre P. SUnSET, a nonradioactive method to monitor protein synthesis. Nat Methods. 2009;6(4):275–7.

30. Sandri M, Sandri C, Gilbert A, Skurk C, Calabria E, Picard A, et al. Foxo transcription factors induce the atrophy-related ubiquitin ligase atrogin-1 and cause skeletal muscle atrophy. Cell. 2004;117(3):399–412.

31. Pittet D, Thievent B, Wenzel RP, Li N, Auckenthaler R, and Suter PM. Bedside prediction of mortality from bacteremic sepsis. A dynamic analysis of ICU patients. Am J Respir Crit Care Med. 1996;153(2):684–93.

32. Cauvi DM, Song D, Vazquez DE, Hawisher D, Bermudez JA, Williams MR, et al. Period of irreversible therapeutic intervention during sepsis correlates with phase of innate immune dysfunction. The Journal of biological chemistry. 2012;287(24):19804–15.

33. Nemzek JA, Xiao HY, Minard AE, Bolgos GL, and Remick DG. Humane endpoints in shock research. Shock. 2004;21(1):17–25.

34. Block BA. Thermogenesis in muscle. Annual review of physiology. 1994;56:535–77.

35. Rowland LA, Bal NC, and Periasamy M. The role of skeletal-muscle-based thermogenic mechanisms in vertebrate endothermy. Biological reviews of the Cambridge Philosophical Society. 2015;90(4):1279–97.

36. Karsli-Uzunbas G, Guo JY, Price S, Teng X, Laddha SV, Khor S, et al. Autophagy is required for glucose homeostasis and lung tumor maintenance. Cancer discovery. 2014;4(8):914–27.

37. Gomes MD, Lecker SH, Jagoe RT, Navon A, and Goldberg AL. Atrogin-1, a muscle-specific F-box protein highly expressed during muscle atrophy. Proceedings of the National Academy of Sciences of the United States of America. 2001;98(25):14440–5.

38. Bodine SC, Latres E, Baumhueter S, Lai VK, Nunez L, Clarke BA, et al. Identification of ubiquitin ligases required for skeletal muscle atrophy. Science. 2001;294(5547):1704–8.

39. Sartori R, Schirwis E, Blaauw B, Bortolanza S, Zhao J, Enzo E, et al. BMP signaling controls muscle mass. Nature genetics. 2013;45(11):1309–18.

40. Milan G, Romanello V, Pescatore F, Armani A, Paik J-H, Frasson L, et al. Regulation of autophagy and the ubiquitin–proteasome system by the FoxO transcriptional network during muscle atrophy. Nature Communications. 2015;6:6670.

41. Lo Verso F, Carnio S, Vainshtein A, and Sandri M. Autophagy is not required to sustain exercise and PRKAA1/AMPK activity but is important to prevent mitochondrial damage during physical activity. Autophagy. 2014;10(11):1883–94.

42. Rooyackers OE, Kersten AH, and Wagenmakers AJ. Mitochondrial protein content and in vivo synthesis rates in skeletal muscle from critically ill rats. Clin Sci (Lond). 1996;91(4):475–81.

43. Callahan LA, and Supinski GS. Sepsis induces diaphragm electron transport chain dysfunction and protein depletion. Am J Respir Crit Care Med. 2005;172(7):861–8.

44. Crouser ED, Julian MW, Blaho DV, and Pfeiffer DR. Endotoxin-induced mitochondrial damage correlates with impaired respiratory activity. Crit Care Med. 2002;30(2):276–84.

45. Protti A, Carre J, Frost MT, Taylor V, Stidwill R, Rudiger A, et al. Succinate recovers mitochondrial oxygen consumption in septic rat skeletal muscle. Crit Care Med. 2007;35(9):2150–5.

46. Crowell KT, Soybel DI, and Lang CH. Restorative Mechanisms Regulating Protein Balance in Skeletal Muscle During Recovery From Sepsis. Shock. 2017;47(4):463–73.

47. Cahill GF. Fuel Metabolism in Starvation. Annu Rev Nutr Annual Review of Nutrition. 2006;26(1):1–22.

48. Zhou C, and Byard RW. Septic Ketoacidosis-A Potentially Lethal Entity with Renal Tubular Epithelial Vacuolization. J Forensic Sci. 2017;62(1):122–5.

49. Ezaki J, Matsumoto N, Takeda-Ezaki M, Komatsu M, Takahashi K, Hiraoka Y, et al. Liver autophagy contributes to the maintenance of blood glucose and amino acid levels. Autophagy. 2011;7(7):727–36.

50. Boyd JH, Divangahi M, Yahiaoui L, Gvozdic D, Qureshi S, and Petrof BJ. Toll-like receptors differentially regulate CC and CXC chemokines in skeletal muscle via NF-kappaB and calcineurin. Infect Immun. 2006;74(12):6829–38.

51. Nakahira K, Haspel JA, Rathinam VA, Lee SJ, Dolinay T, Lam HC, et al. Autophagy proteins regulate innate immune responses by inhibiting the release of mitochondrial DNA mediated by the NALP3 inflammasome. Nat Immunol. 2011;12(3):222–30.

52. Zhou R, Yazdi AS, Menu P, and Tschopp J. A role for mitochondria in NLRP3 inflammasome activation. Nature. 2011;469(7329):221-5.

53. Sun Y, Yao X, Zhang Q-J, Zhu M, Liu Z-P, Ci B, et al. Beclin-1-Dependent Autophagy Protects the Heart During Sepsis. Circulation. 2018;138(20):2247–62.

54. Oami T, Watanabe E, Hatano M, Teratake Y, Fujimura L, Sakamoto A, et al. Blocking Liver Autophagy Accelerates Apoptosis and Mitochondrial Injury in Hepatocytes and Reduces Time to Mortality in a Murine Sepsis Model. Shock. 2018;50(4):427–34.

55. Thiessen SE, Derese I, Derde S, Dufour T, Pauwels L, Bekhuis Y, et al. The Role of Autophagy in Critical Illness-induced Liver Damage. Sci Rep. 2017;7(1):14150.

56. Oami T, Watanabe E, Hatano M, Sunahara S, Fujimura L, Sakamoto A, et al. Suppression of T Cell Autophagy Results in Decreased Viability and Function of T Cells Through Accelerated Apoptosis in a Murine Sepsis Model. Critical care medicine. 2017;45(1):e77–e85.

57. Komatsu M, Waguri S, Ueno T, Iwata J, Murata S, Tanida I, et al. Impairment of starvation-induced and constitutive autophagy in Atg7-deficient mice. J Cell Biol. 2005;169(3):425–34.

58. Schuler M, Ali F, Metzger E, Chambon P, and Metzger D. Temporally controlled targeted somatic mutagenesis in skeletal muscles of the mouse. Genesis. 2005;41(4):165–70.

59. Kiermayer C, Conrad M, Schneider M, Schmidt J, and Brielmeier M. Optimization of spatiotemporal gene inactivation in mouse heart by oral application of tamoxifen citrate. Genesis. 2007;45(1):11–6.

60. Robinson SP, Langan-Fahey SM, Johnson DA, and Jordan VC. Metabolites, pharmacodynamics, and pharmacokinetics of tamoxifen in rats and mice compared to the breast cancer patient. Drug metabolism and disposition: the biological fate of chemicals. 1991;19(1):36–43.

61. Buras JA, Holzmann B, and Sitkovsky M. Animal models of sepsis: setting the stage. Nat Rev Drug Discov. 2005;4(10):854–65.

62. Ruiz S, Vardon-Bounes F, Merlet-Dupuy V, Conil J-M, Buléon M, Fourcade O, et al. Sepsis modeling in mice: ligation length is a major severity factor in cecal ligation and puncture. Intensive care medicine experimental. 2016;4(1):22.

63. Toscano MG, Ganea D, and Gamero AM. Cecal ligation puncture procedure. J Vis Exp. 2011(51).

64. Gonnert FA, Recknagel P, Seidel M, Jbeily N, Dahlke K, Bockmeyer CL, et al. Characteristics of clinical sepsis reflected in a reliable and reproducible rodent sepsis model. J Surg Res. 2011;170(1):e123–34.

65. Leduc-Gaudet J-P, Reynaud O, Hussain SN, and Gouspillou G. Parkin overexpression protects from ageing-related loss of muscle mass and strength. The Journal of Physiology. 2019;0(0).

66. Mofarrahi M, McClung JM, Kontos CD, Davis EC, Tappuni B, Moroz N, et al. Angiopoietin-1 enhances skeletal muscle regeneration in mice. Am J Physiol Regul Integr Comp Physiol. 2015;308(7):R576–89.

67. Gouspillou G, Godin R, Piquereau J, Picard M, Mofarrahi M, Mathew J, et al. Protective role of Parkin in skeletal muscle contractile and mitochondrial function. The Journal of physiology. 2018;596(13):2565–79.

68. Leduc-Gaudet JP, Picard M, St-Jean Pelletier F, Sgarioto N, Auger MJ, Vallee J, et al. Mitochondrial morphology is altered in atrophied skeletal muscle of aged mice. Oncotarget. 2015;6(20):17923–37.

69. Briguet A, Courdier-Fruh I, Foster M, Meier T, and Magyar JP. Histological parameters for the quantitative assessment of muscular dystrophy in the mdx-mouse. Neuromuscul Disord. 2004;14(10):675–82.

70. Dulac M, Leduc-Gaudet JP, Reynaud O, Ayoub MB, Guerin A, Finkelchtein M, et al. Drp1 knockdown induces severe muscle atrophy and remodelling, mitochondrial dysfunction, autophagy impairment and denervation. The Journal of physiology. 2020;598(17):3691–710.

71. Ritchie ME, Phipson B, Wu D, Hu Y, Law CW, Shi W, et al. limma powers differential expression analyses for RNA-sequencing and microarray studies. Nucleic Acids Res. 2015;43(7):e47.

