## Supplementary Figures for "Role of autophagy in sepsis-induced skeletal muscle dysfunction, whole-body metabolism, and survival"

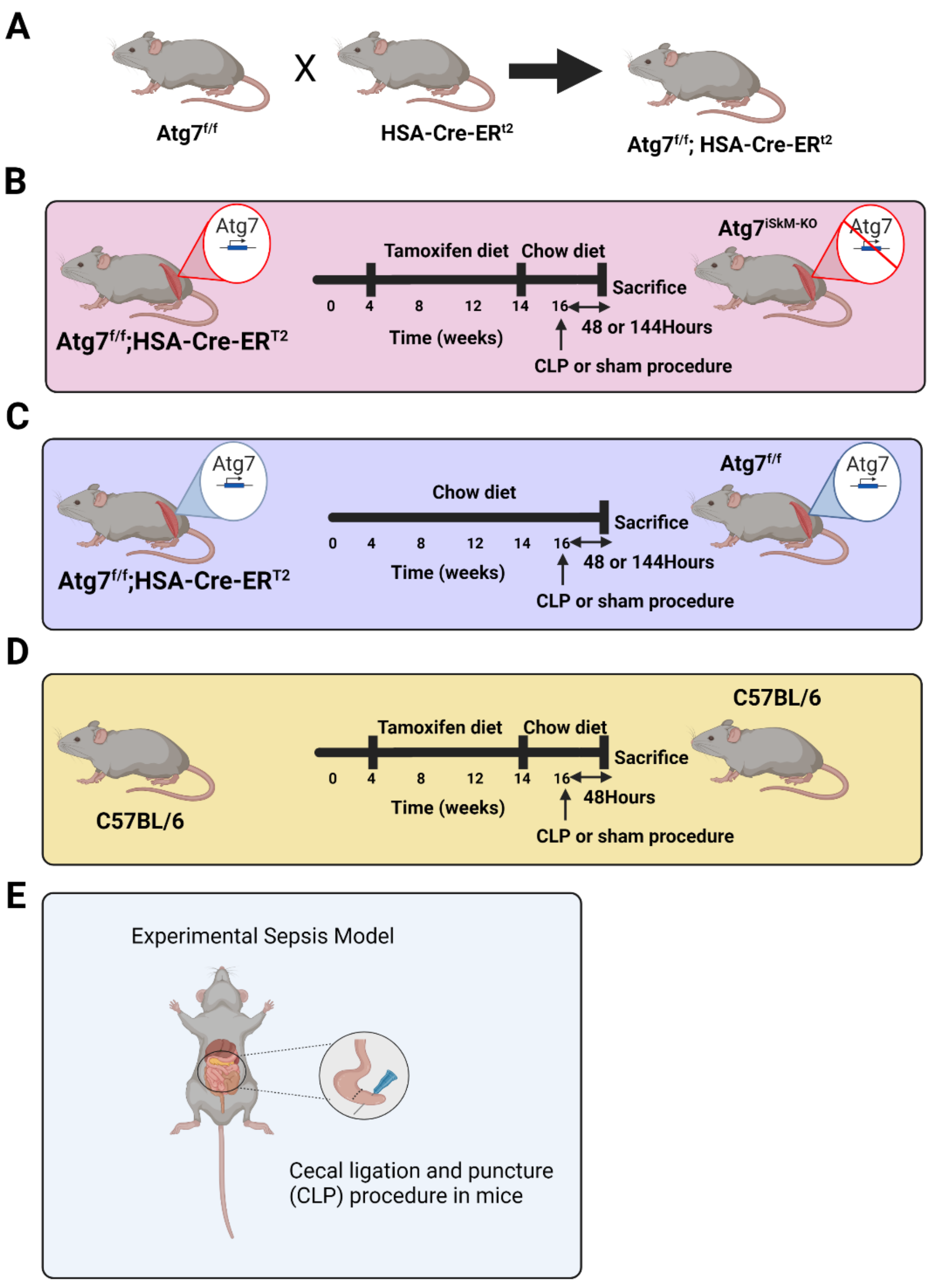


**Figure S1: Generation of skeletal muscle-specific conditional Atg7 knockout mice.**
**A)** Schematic diagram of the experimental design showing breeding, diet, and sepsis induction strategies used to delete Atg7 in murine skeletal muscles. **B)** We found that tamoxifen diet for 10 weeks follow by a washout period of 2-3 weeks is the most suitable to generate mice with skeletal muscle-specific conditional Atg7 knockout (Atg7$^{iSkM\text{-}KO}$). **C)** Sex and age-matched HSA-Cre-ERT2 Atg7$^{f/f}$ mice fed a standard chow diet are referred to as Atg7$^{f/f}$ mice. **D)** To control for the effects of tamoxifen diet, C57BL6/J were also studied. No effect of tamoxifen diet on any of our outcome measurements, including muscle mass, muscle contractility and mitochondrial respiration, could be evidenced (data not shown). **E)** Cecal ligation and perforation (CLP) mouse model.

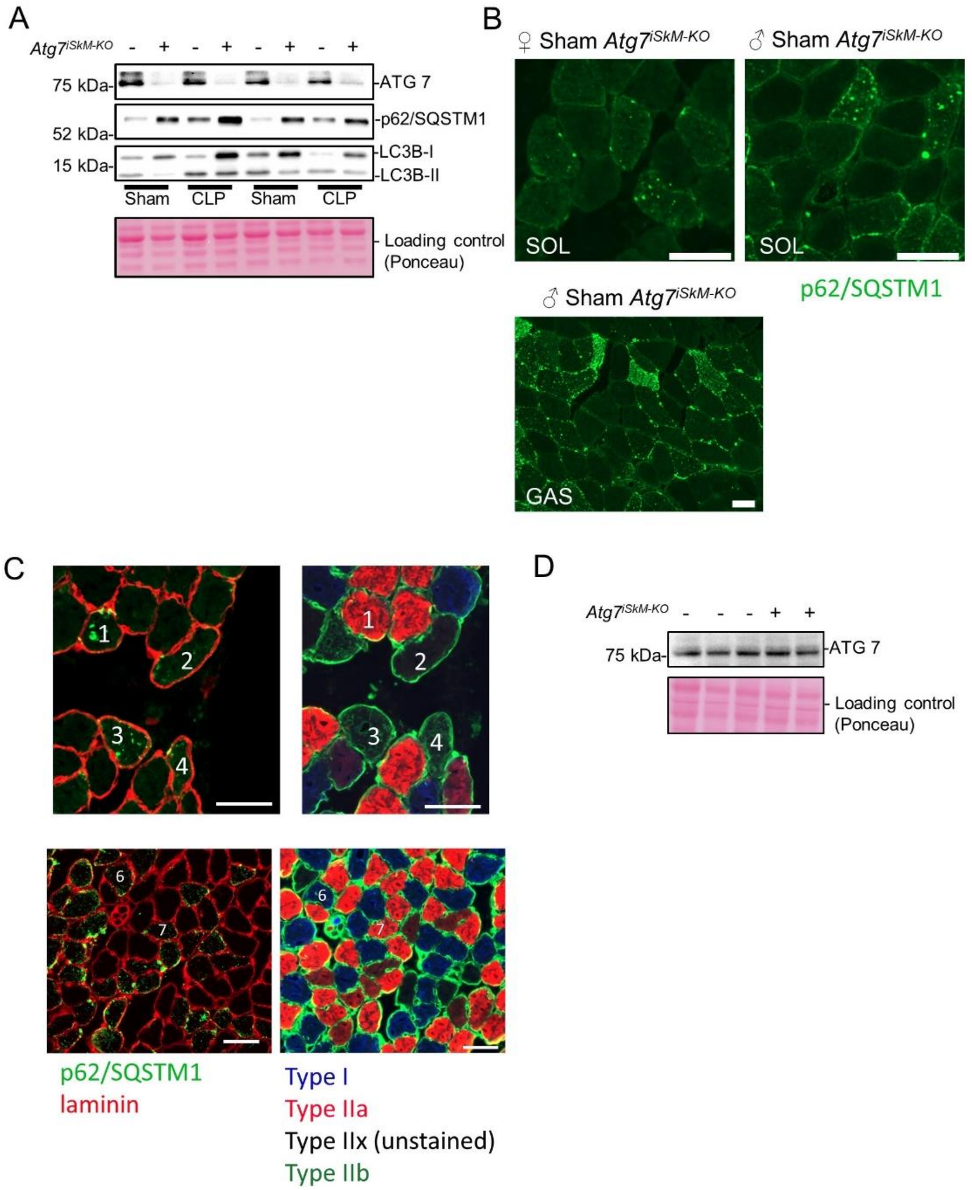


**Figure S2: Confirmation of autophagy inactivation in skeletal muscles and regulation of autophagy-related genes.**

**A)** Representative immunoblots showing ATG7, p62, and LC3 proteins in GAS muscles of female Atg7$^{f/f}$ (intact) and Atg7$^{iSkM-KO}$ (knockout) mice 48 h after sham surgery or CLP showing that sepsis activates autophagy in Atg7$^{f/f}$ mice and that LC3 lipidation is impaired in Atg7$^{iSkM-KO}$ mice. Ponceau serves as loading control. **B)** Immunostaining for p62/SQSTM1 showing accumulation of aggregates in oxidative (SOL) and glycolytic (GAS) muscles. **C)** Left panels show laminin (red) and p62/SQSTM1 (green) immunostaining of SOL muscles of female Atg7$^{iSkM-KO}$ 48 h after sham surgery. The panels on the right show the corresponding MHC (red = MHCIIa; green = MHCIIb; and black (unstained) = MHCIIx) and laminin (green) immunostain obtained on a serial cross-section. Numbers provided in this panel highlight the same fibers on muscle cross-sections. p62/SQSTM1 positive staining, revealing p62/SQSTM1 accumulation, was found in all fiber types. Scale bar=50 mm. **D)** Representative immunoblot showing that Atg7 remains unaffected in cardiac muscle.

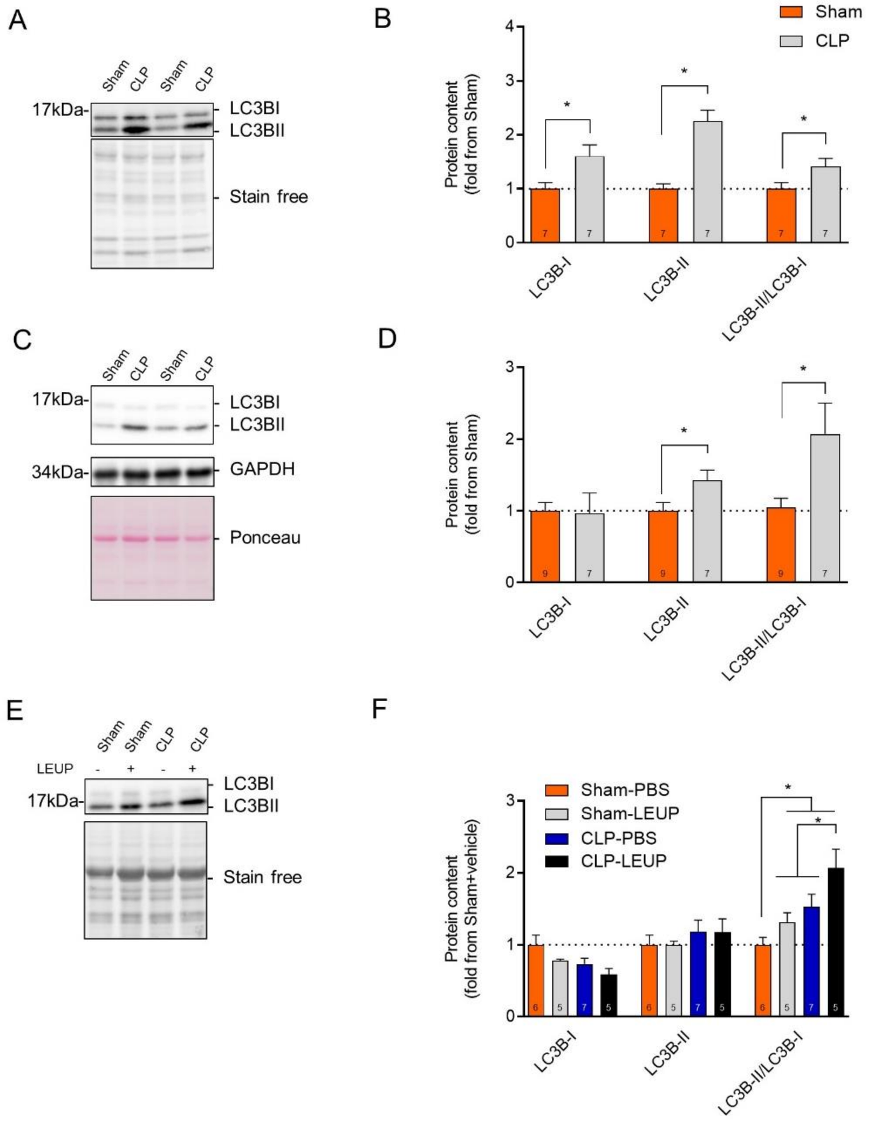


**Figure S3: The impact of sepsis on skeletal muscle autophagy.**

**A-D)** Representative LC3 immunoblots and densitometric analyses from diaphragm (A-B) and TA (C-D) muscles of C57BL6/J mice 48 hours post sham or CLP surgery. **E-F)** Representative LC3 immunoblots and quantification of the autophagic flux in TA muscle from C57BL6/J mice 24 hours post sham or CLP surgery. The lysosomal protease inhibitors leupeptin (LEUP, 40 mg/kg IP) was used to block the autophagic flux. Mice were sacrificed 4 hours later. Control sham and CLP groups received an equivalent volume of sterile PBS. Mice were divided into four groups: Sham-PBS (sham mice injected with PBS); Sham-LEUP (sham mice injected with leupeptin); CLP-PBS (CLP mice injected with PBS); CLP-LEUP (CLP mice injected with leupeptin). Ponceau S or stain-free stains were used as loading controls. The number of animals for each group is indicated within bars. Data are presented as fold change relative to Sham-PBS and are expressed as mean ± SEM. * $p < 0.05$

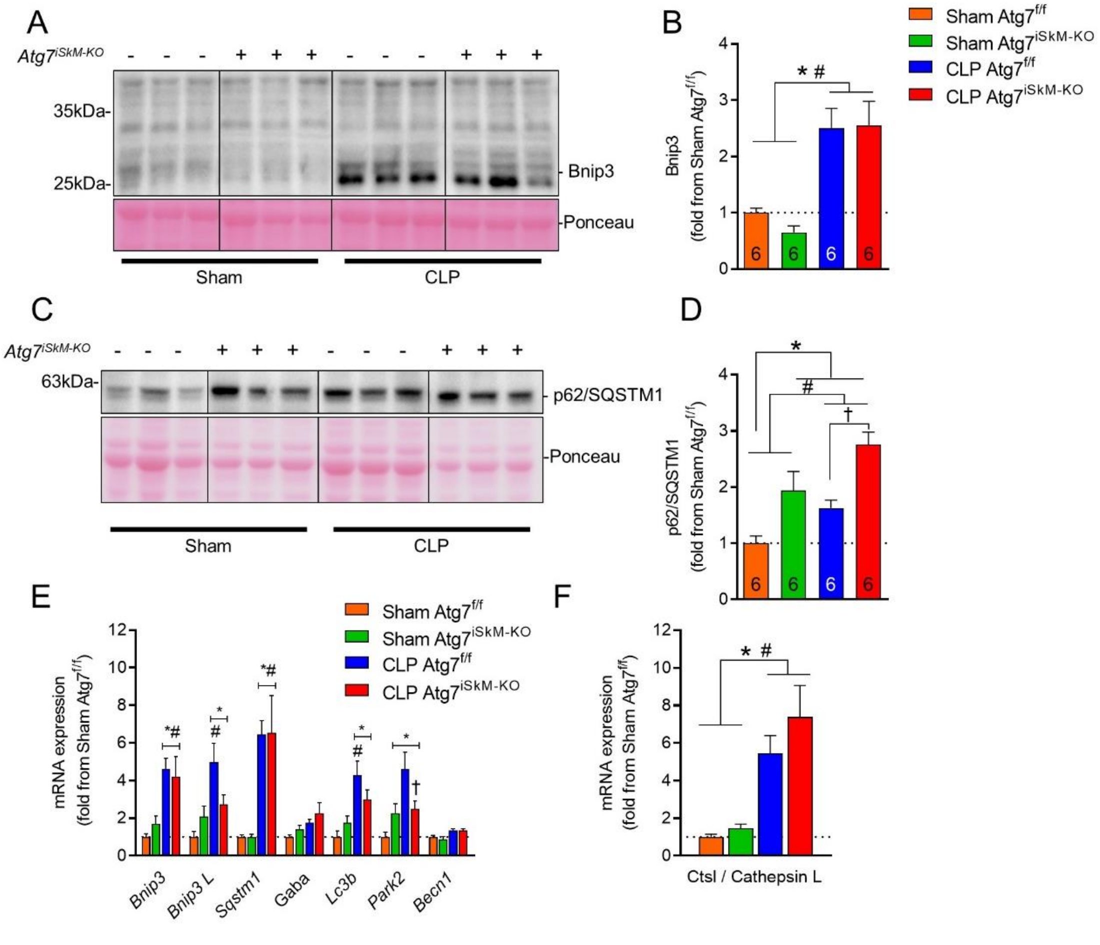


**Figure S4: Autophagy inactivation in skeletal muscles and regulation of autophagy-related genes.**

**A-D)** Representative immunoblots and densitometric analyses of Bnip3 and p62/SQSTM1 levels in GAS muscles of female Atg7$^{f/f}$ and Atg7$^{iSkM\text{-}KO}$ mice 48 h after sham surgery or CLP. Both proteins are significantly upregulated by sepsis. Ponceau S were used as loading control. Number of animals indicated within bars. **E-F)** mRNA expressions of various autophagy-related genes and lysosomal cathepsin in GAS muscles of female Atg7$^{f/f}$ and Atg7$^{iSkM\text{-}KO}$ mice 48 h after sham surgery or CLP showing significant upregulation in response to sepsis**.** Gaba refers to Gabarapl1 and Becn1 refers to Beclin1. Data presented as fold change relative to sham Atg7$^{f/f}$ and as mean ± SEM. * $p < 0.05$ vs sham Atg7$^{f/f}$; # $p < 0.05$ for sepsis effect (i.e. sham Atg7$^{f/f}$ vs. CLP Atg7$^{f/f}$ or sham Atg7$^{iSkM\text{-}KO}$ vs. CLP Atg7$^{iSkM\text{-}KO}$); † $p < 0.05$ for sepsis plus knockout effect (i.e. CLP Atg7$^{f/f}$ vs.CLP Atg7$^{iSkM\text{-}KO}$).

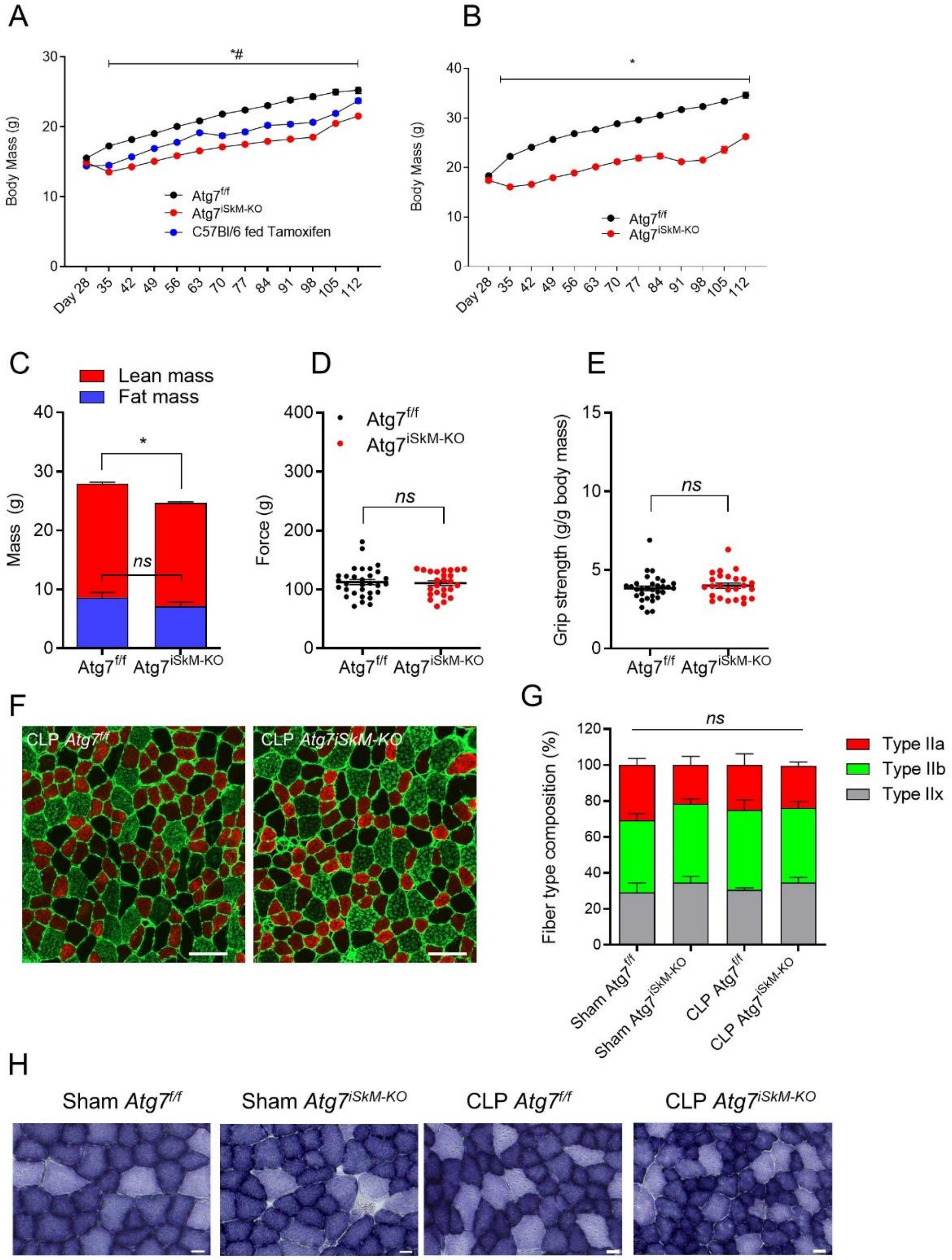


**Figure S5: Changes in body mass, muscle fiber size and fiber composition.**
**A-B)** Body mass of female Atg7$^{f/f}$ and Atg7$^{iSkM\text{-}KO}$ mice and age-matched tamoxifen-fed C57Bl/6 mice (panel A) and male Atg7$^{f/f}$ and Atg7$^{iSkM\text{-}KO}$ mice (panel B) as a function of age (in days). Animals were weighed weekly and a minimum of 25 mice were tracked per group. In both sexes, significant reductions in body mass were observed in Atg7$^{iSkM\text{-}KO}$ mice as compared to Atg7$^{f/f}$ mice from the age of 35 days on, independent of tamoxifen diet. Data presented as mean ± SEM. Knockout effect indicated by $^{*}p < 0.05$, Atg7$^{f/f}$ vs. Atg7$^{iSkM\text{-}KO}$. Comparison of Atg7 mice to C57Bl/6 mice indicated by $^{\#}p < 0.05$, Atg7$^{f/f}$ vs. C57Bl/6 fed Tamoxifen. **C)** EchoMRI-measured

body composition of female Atg7$^{f/f}$ (n = 28) and Atg7$^{iSkM\text{-}KO}$ (n = 24) mice 3 days prior to sham surgery or CLP. Atg7 knockout results in significant reductions of lean mass. Fat mass is unaffected. **D-E)** No differences in forelimb grip strength were observed between Atg7$^{f/f}$ (n = 31) and Atg7$^{iSkM\text{-}KO}$ (n = 25) mice in terms of absolute force (D) or force normalized to lean mass (E). Knockout effect indicated by *$p < 0.05$. *ns* = no significant difference. **F)** Representative images of MHC-labeled TA muscles (red = MHCIIa; green = MHCIIb; and black (unstained) = MHCIIx) from female CLP Atg7$^{f/f}$ and CLP Atg7$^{iSkM\text{-}KO}$ mice 2 days after sham surgery. Scale bars = 100 µm. **G)** Percent fiber type composition of TA muscles. **H)** Representative images of succinate dehydrogenase (SDH) staining in TA muscles of female Atg7$^{f/f}$ and Atg7$^{iSkM\text{-}KO}$ mice showing no significant differences in oxidative capacity or glycolytic potential as a consequence of Atg7 knockout or sepsis, although myofiber size decreases in both knockout muscles and septic muscles. Scale bar = 20µm.

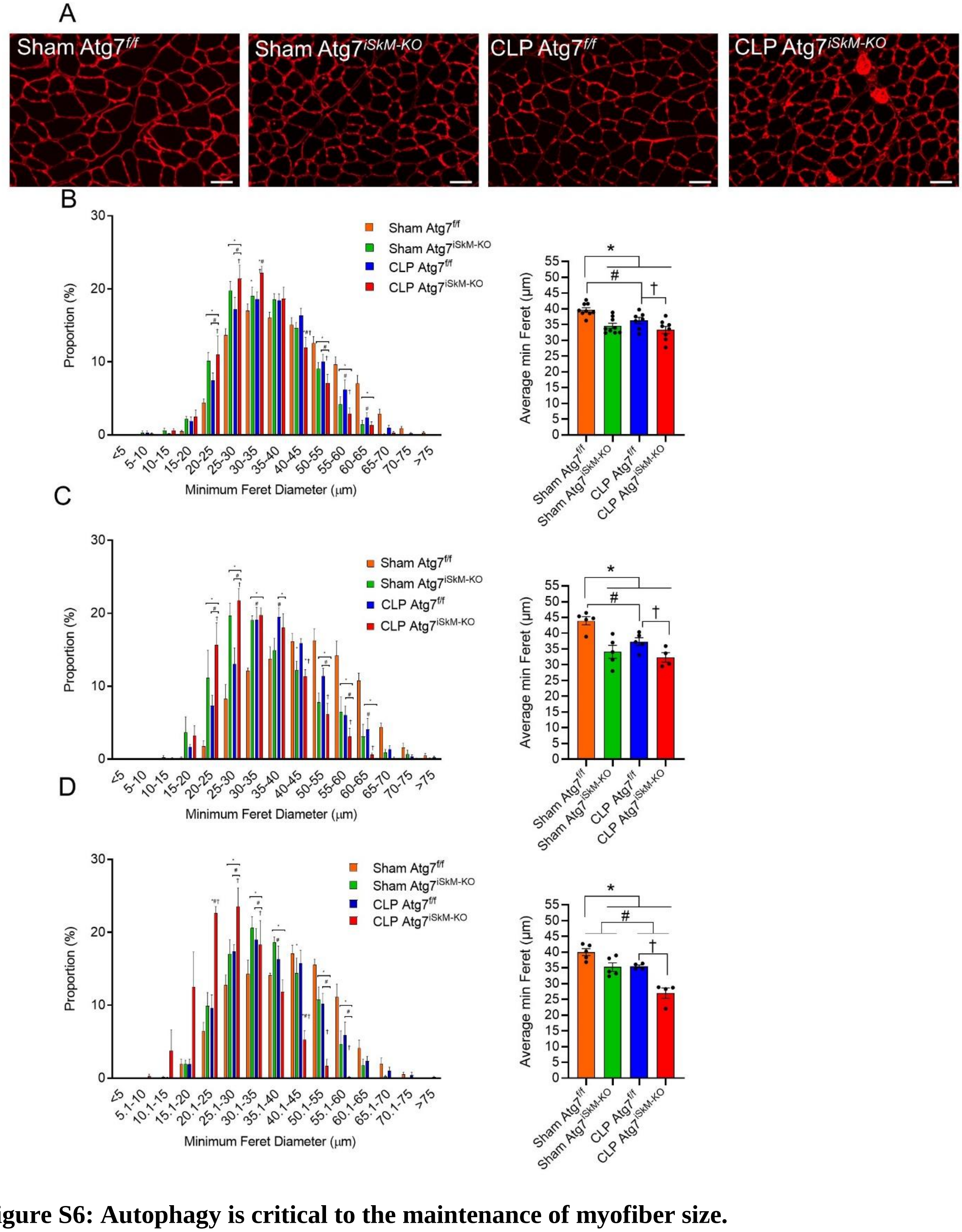


**Figure S6: Autophagy is critical to the maintenance of myofiber size.**

A) Representative laminin-stained TA muscle cryosection image from female Atg7$^{f/f}$ and Atg7$^{iSkM-KO}$ mice 48 h after sham surgery or CLP. B) Distribution of myofiber diameter in TA muscles of female Atg7$^{f/f}$ and Atg7$^{iSkM-KO}$ mice 48 h after sham surgery or CLP by minimum Feret diameter (left panel) and average minimum Feret diameter (right panel). Sham Atg7$^{f/f}$ (n = 9; 433 ± 25 fibers per mice were counted), sham Atg7$^{iSkM-KO}$ (n = 9; 420 ± 35 fibers per mice were counted), CLP Atg7$^{f/f}$ (n = 7; 390 ± 21 fibers per mice were counted), and CLP Atg7$^{iSkM-KO}$ (n = 8; 432 ± 32 fibers per mice were counted). C) Distribution of myofiber diameter in TA muscles of male Atg7$^{f/f}$ and Atg7$^{iSkM-KO}$ mice 48 h after sham surgery or CLP by minimum Feret diameter (left panel) and average minimum Feret diameter (right panel). Sham Atg7$^{f/f}$ (n = 5; 339 ± 31 fibers per mice were counted), sham Atg7$^{iSkM-KO}$ (n = 5; 427 ± 26 fibers per mice were counted), CLP Atg7$^{f/f}$ (n = 5; 336 ± 10 fibers per mice were counted), and CLP Atg7$^{iSkM-KO}$ (n = 8; 345 ± 18 fibers per mice were counted). D) Distribution of myofiber diameter in TA muscles of female Atg7$^{f/f}$ and Atg7$^{iSkM-KO}$ mice 6 days after sham surgery or CLP by minimum Feret diameter (left panel) and average minimum Feret diameter (right panel). Sham Atg7$^{f/f}$ (n = 5; 414 ± 21 fibers per mice were counted), sham Atg7$^{iSkM-KO}$ (n = 5; 491 ± 28 fibers per mice were counted), CLP Atg7$^{f/f}$ (n = 4; 440 ± 68 fibers per mice were counted), and CLP Atg7$^{iSkM-KO}$ (n = 4; 603 ± 101 fibers per mice were counted). Data presented as mean ± SEM. * $p < 0.05$ vs sham Atg7$^{f/f}$; # $p < 0.05$ for sepsis effect (i.e. sham Atg7$^{f/f}$ vs. CLP Atg7$^{f/f}$ or sham Atg7$^{iSkM-KO}$ vs. CLP Atg7$^{iSkM-KO}$); † $p < 0.05$ for sepsis plus knockout effect (i.e. CLP Atg7$^{f/f}$ vs.CLP Atg7$^{iSkM-KO}$).

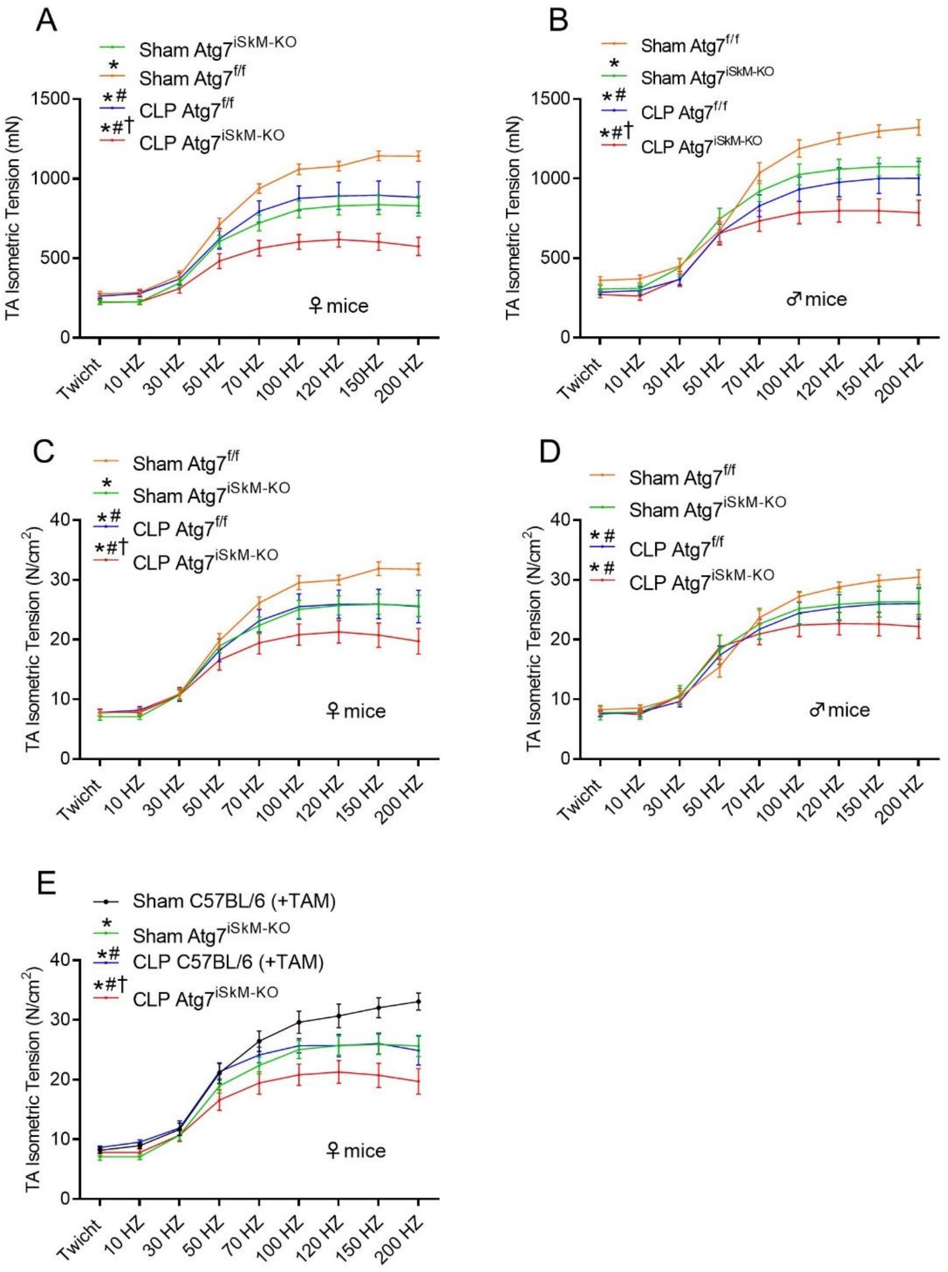


**Figure S7: Autophagy inactivation and sepsis impair muscle contractility.**

**A-D)** Absolute (panels A&B) and specific force (panels C&D) of TA muscles from female (panels A&C) and male (panels B&D) Atg7$^{f/f}$ and Atg7$^{iSkM-KO}$ mice measured *in situ* over a range of stimulation frequencies 48 h after sham surgery or CLP, showing that muscles from septic knockout mice are the most severely impaired. **E)** Specific force of TA muscles from female Atg7$^{f/f}$ and age-matched tamoxifen-fed C57BL/6 mice over a range of stimulation frequencies measured 48 h after sham surgery or CLP. Data in panels A-E presented as mean ± SEM. Sample sizes in Figures A, C, and E as follows: sham Atg7$^{f/f}$ (n = 12); sham Atg7$^{f/f}$HSA-Cre-ER$^{T2}$ (+TAM) (also referred to as sham Atg7$^{iSkM-KO}$) (n = 12); CLP Atg7$^{f/f}$ (n = 10); CLP Atg7$^{f/f}$HSA-Cre-ER$^{T2}$ (+TAM) (also referred to as CLP Atg7$^{iSkM-KO}$) (n = 12); sham C57BL/6 (+TAM) (n = 8); and CLP C57BL/6 (+TAM) (n = 5). Sample sizes in Figures B and D as follows: sham Atg7$^{f/f}$ (n = 8); sham

$Atg7^{-/-}$ (n = 8); CLP $Atg7^{f/f}$ (n = 8) and CLP $Atg7^{-/-}$ (n = 11). * $p < 0.05$ vs sham $Atg7^{f/f}$ (panels A-D) and * $p < 0.05$ vs sham C57Bl/6 (+TAM) (panel E). Sepsis effect indicated by #$p < 0.05$, sham $Atg7^{f/f}$ vs. CLP $Atg7^{f/f}$ or sham $Atg7^{-/-}$ vs. CLP $Atg7^{-/-}$ (panels A-D) and #$p < 0.05$, sham C57BI/6 (+TAM) vs. CLP C57Bl/6 (+TAM) (panel E). Sepsis plus knockout effect indicated by †$p < 0.05$, CLP $Atg7^{f/f}$ vs.CLP $Atg7^{-/-}$ (panels A-D) or CLP C57Bl/6 (+TAM) vs. CLP $Atg7^{f/f}$HSA-Cre-$ER^{T2}$+TAM (panel E).

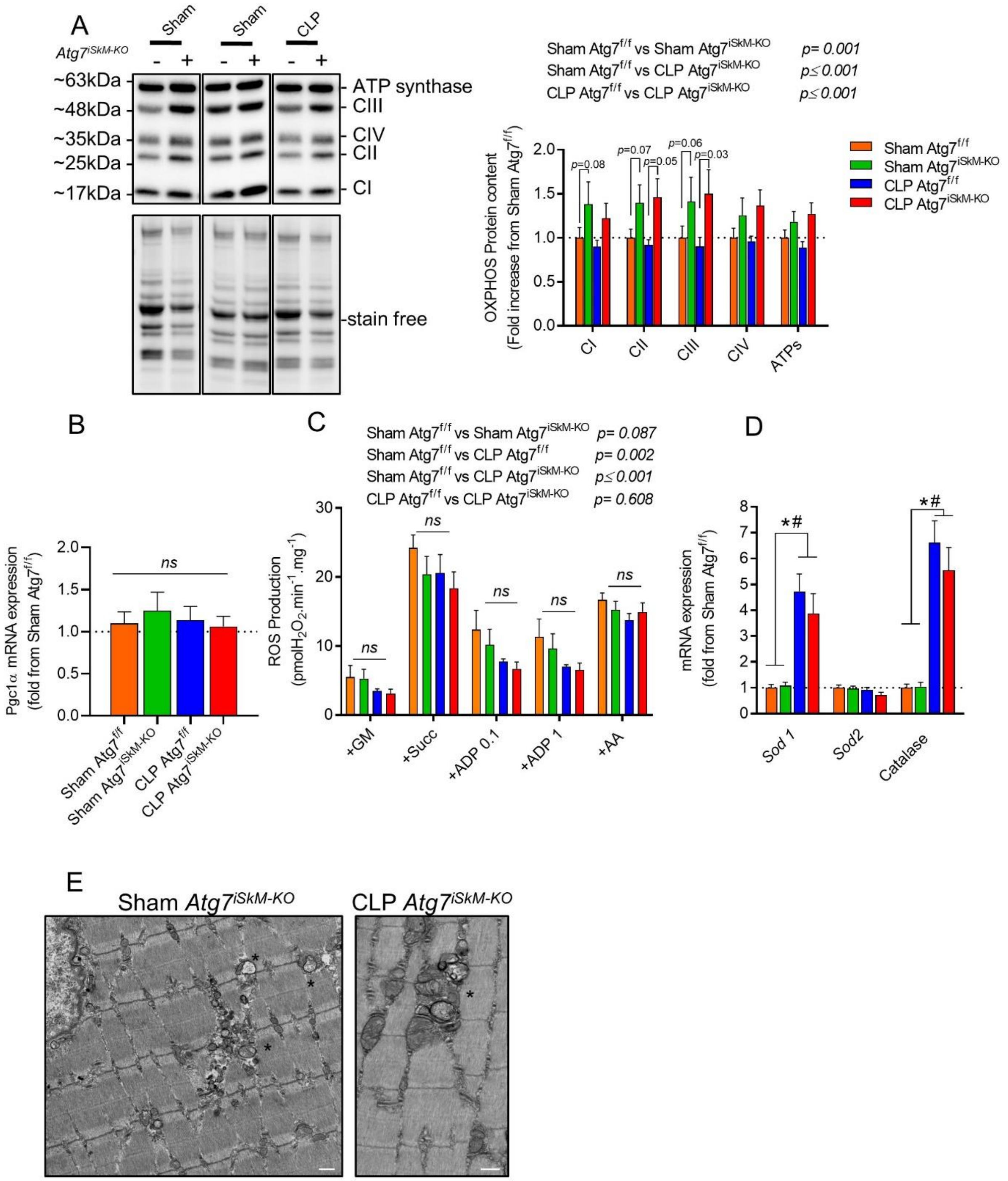


**Figure S8. Autophagy inactivation impairs mitochondria but has no effect on mitochondrial ROS production.**

**A)** Representative immunoblots of mitochondrial respiratory chain subunits (OXPHOS) in GAS muscles of female Atg7$^{f/f}$ and Atg7$^{iSkM-KO}$ mice 48 h after sham surgery or CLP (n = 8 for each sham group, n = 5 each CLP group). Stain-free technology was used to normalize mitochondrial respiratory chain subunits. Significant increases in sham and septic knockout mice suggest accumulation of impaired mitochondria. **B)** mRNA expression of PGC1$\alpha$ in TA muscles of male Atg7$^{f/f}$ and Atg7$^{iSkM-KO}$ mice 48 h post sham or CLP surgery (n = 6-8 per group). **C)** Mitochondrial

ROS production in GAS muscles of female Atg7$^{f/f}$ and Atg7$^{iSkM\text{-}KO}$ mice 48 h after sham surgery or CLP (n = 8 for each sham group, n = 7 for each CLP group) showing that ROS production is unaffected by autophagy inactivation or sepsis. **D)** mRNA expressions of antioxidant defense-related genes in TA muscles of female Atg7$^{f/f}$ and Atg7$^{iSkM\text{-}KO}$ mice 48 h after sham surgery or CLP (n = 6-8 for each group). **E)** Electron micrographs of GAS muscles from Atg7$^{iSkM\text{-}KO}$ mice 48 h after sham or CLP surgery (n = 2 per group*)* showing accumulation of abnormal mitochondria (black asterisks). Scale bars: 0.5 μm. Data presented as mean ± SEM. *ns* = not significant. * p <0.05 vs sham Atg7$^{f/f}$; # p <0.05 for sepsis effect (i.e. sham Atg7$^{f/f}$ vs. CLP Atg7$^{f/f}$ or sham Atg7$^{iSkM\text{-}KO}$ vs. CLP Atg7$^{iSkM\text{-}KO}$).

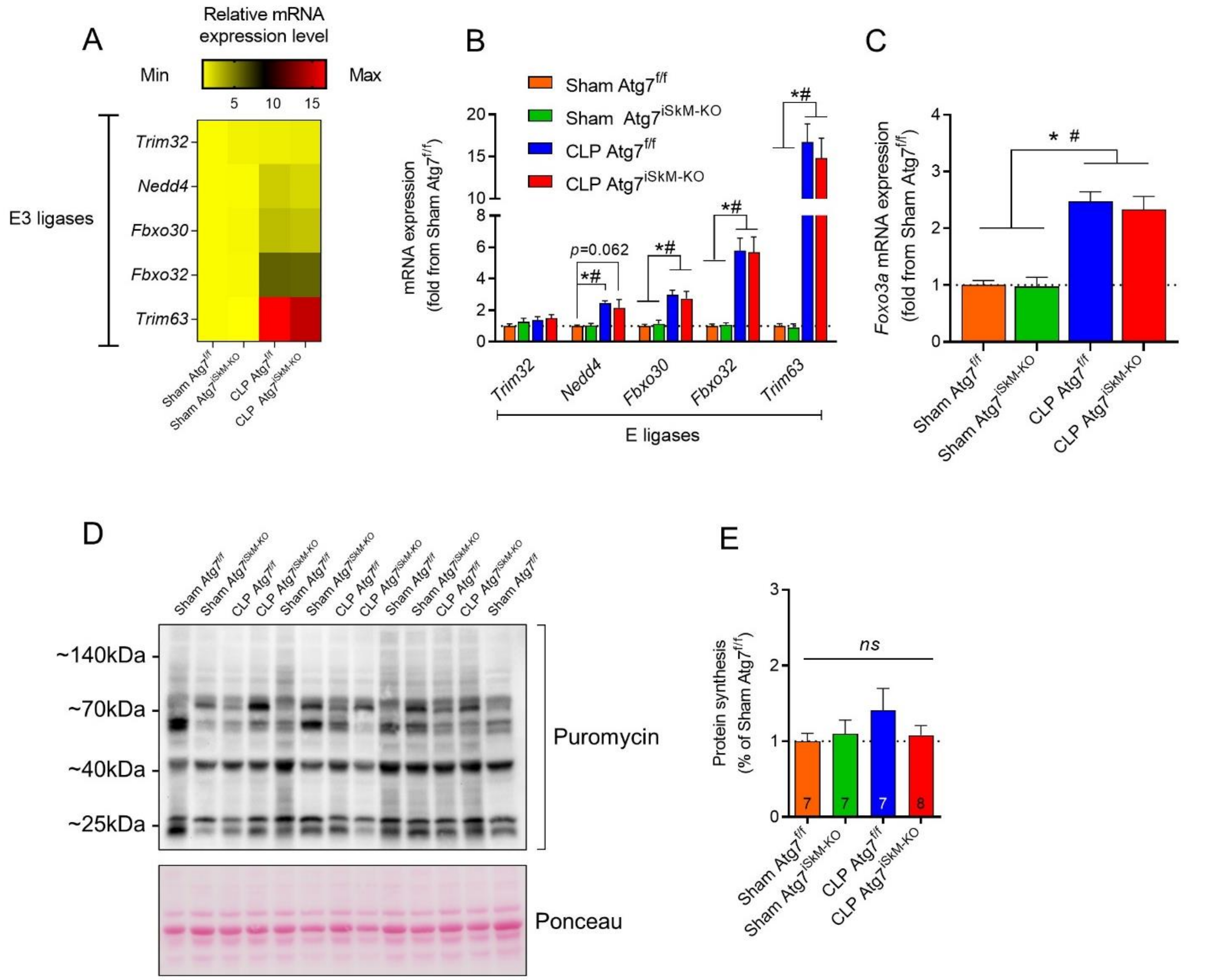


**Figure S9: Autophagy inactivation and sepsis exert differential effects on muscle proteolysis but not on protein synthesis.**

**A)** Heat map summarizing relative mRNA expressions of various E3 ubiquitin ligases involved in skeletal muscle atrophy. Colors indicate relative expression levels, red equals high expression and yellow equals low expression. **B)** mRNA expressions of E3 ubiquitin ligases in GAS muscles of female Atg7$^{f/f}$ and Atg7$^{iSkM-KO}$ mice 48 h after sham surgery or CLP (n = 7-8 for each sham group, n = 6-8 for each CLP group). 18S levels used as a control. **C)** mRNA expression of *Foxo3a* in GAS muscles of female Atg7$^{f/f}$ and Atg7$^{iSkM-KO}$ mice 48 h after sham surgery or CLP (n = 8 for each sham group, n = 7 for each CLP group). **D-E)** Representative immunoblot and corresponding quantification of puromycin incorporation in the TA muscle of male Atg7$^{f/f}$ and Atg7$^{iSkM-KO}$ mice 48 h post sham or CLP surgery using the *in vivo* SUnSET technique. Data in panels B, C & E presented as fold change relative to sham Atg7$^{f/f}$ and as mean ± SEM. * p <0.05 vs sham Atg7$^{f/f}$; # p <0.05 for sepsis effect (i.e. sham Atg7$^{f/f}$ vs. CLP Atg7$^{f/f}$ or sham Atg7$^{iSkM-KO}$ vs. CLP Atg7$^{iSkM-KO}$). *ns* = not significant.

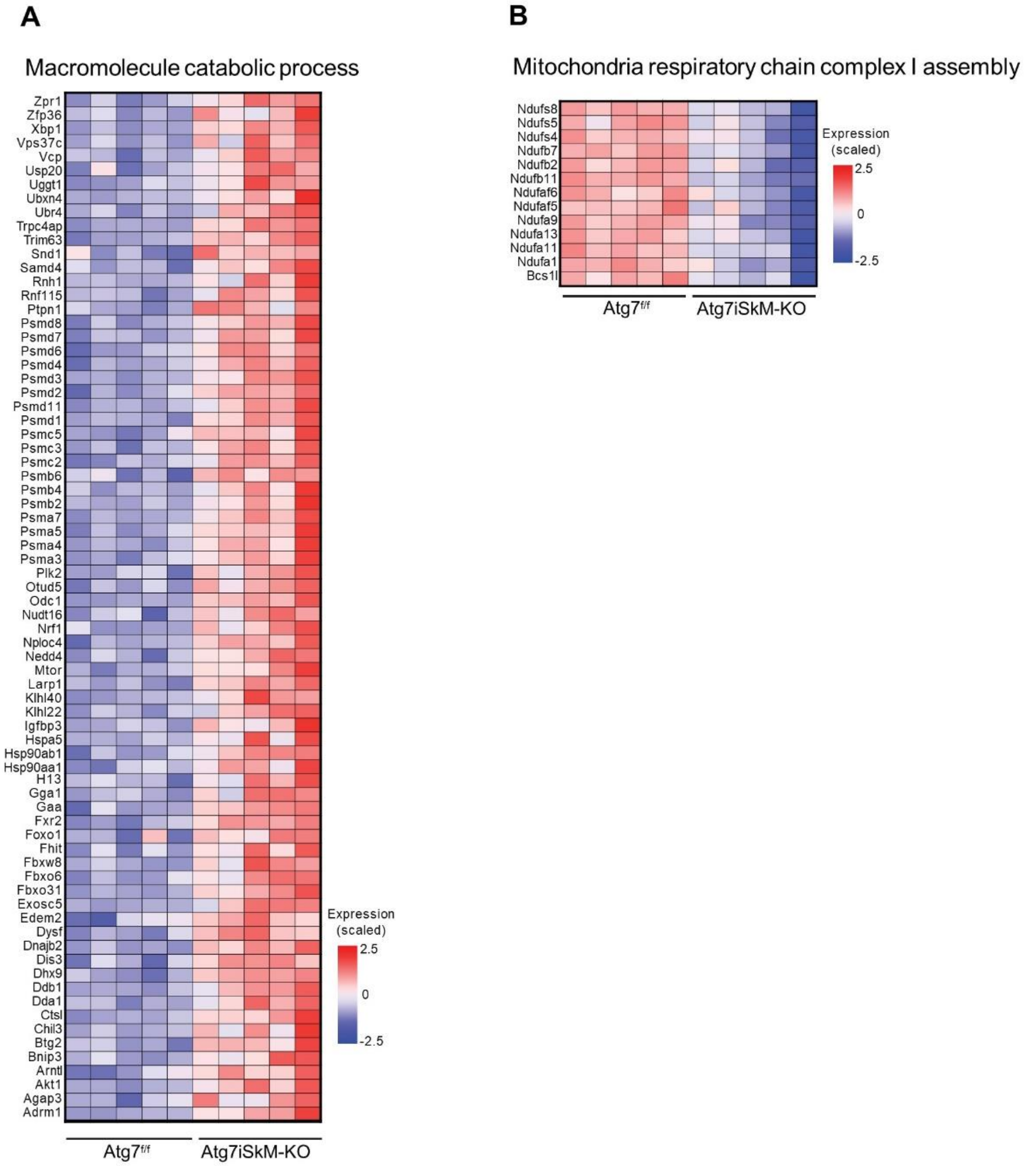


**Figure S10: Autophagy inactivation alters the transcriptional signature of septic muscles.**

**A-B)** Heatmaps showing relative expressions of genes involved in macromolecule catabolism (panel A) and relative expressions of genes involved in the assembly of mitochondrial respiratory chain complex I in muscles of female mice 6 days post CLP surgery (panel B). Colors indicate relative expression levels, red corresponds to high expression while blue corresponds to low expression. n = 5 for each group of CLP mice.

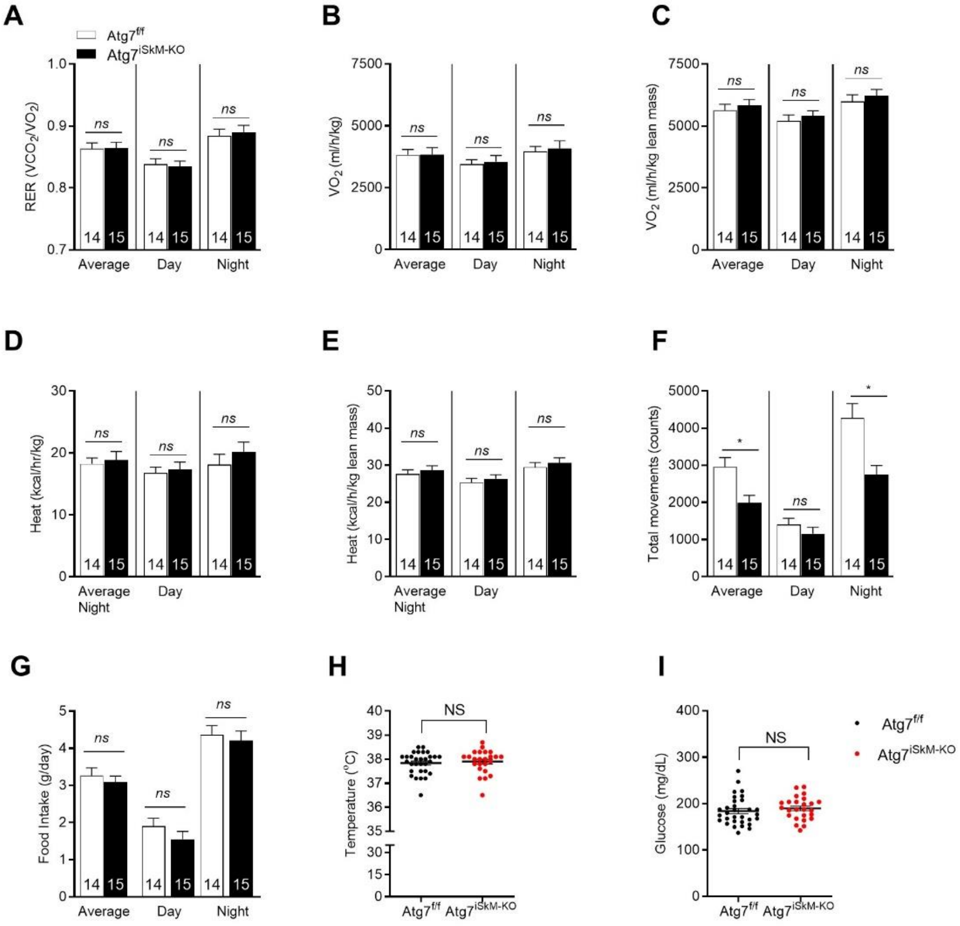


**Figure S11. Autophagy inactivation prior to surgery or CLP has no effect on whole-body metabolism.**

**A)** Respiratory exchange ratio ($VCO_2/VO_2$). **B)** Whole-body oxygen consumption ($VO_2$). **C)** Oxygen consumption. ($VO_2$) relative to lean mass. **D)** Whole body heat (energy expenditure). **E)** Heat (energy expenditure) relative to lean mass. **F)** Total movement (ambulatory activity). **G)** Food intake (g/day). Metabolic cage (PhenoMaster) parameters (A-G) measured over the immediate 2-day period prior to surgery on a dark (6:00 p.m. to 6:00 a.m.)/light (6:00 a.m. to 6:00 p.m.) cycle. Number of animals indicated within bars (lower panels). **H-I)** Non-fasting body temperature (H) and blood glucose levels (I) of female $Atg7^{f/f}$ and $Atg7^{iSkM\text{-}KO}$ mice (n = 30, $Atg7^{f/f}$, n = 24, $Atg7^{iSkM\text{-}KO)}$. Measured between 6-8 a.m. Data presented as mean ± SEM. Knockout effect indicated by *$p < 0.05$, sham $Atg7^{f/f}$ vs. sham $Atg7^{iSkM\text{-}KO}$. *ns* = not significant.
