## Supplementary Table for "Role of autophagy in sepsis-induced skeletal muscle dysfunction, whole-body metabolism, and survival"

### Table of antibodies for western blotting

| Antibody | Source / Product no. | Dilution | Analysis |
|---|---|---|---|
| rabbit anti-Atg7 | Cell signaling #8558 | 1/1000 | WB |
| rabbit anti-LC3 | Cell signaling #12741 | 1/1000 | WB |
| rabbit anti-GAPDH | Cell signaling # 2118 | 1/2500 | WB |
| rabbit anti-phospho-Fox01 (Ser256) | Cell signaling # 9461 | 1/750 | WB |
| rabbit anti-total FoxO1 | Cell signaling # 9454 | 1/750 | WB |
| rabbit anti-phospho-FoxO3a (Ser253) | Cell signaling # 9466 | 1/750 | WB |
| rabbit anti-total FoxO3a | Cell signaling # 2497 | 1/750 | WB |
| mouse anti-Bnip3 | Sigma-Aldrich #B7931 | 1/1000 | WB |
| p62/SQSTM1 | Novus Biologicals Inc. clone 2C11 | 1/1000 | WB |
| Anti-puromycin, clone 12D10 | Millipore # MABE343 | 1/2500 | WB |
| OXPHOS | Abcam #110413 | 1/1000 | WB |
| Goat anti mouse IgG | Abcam # Ab6728 | 1/5000 | WB |
| Goat anti rabbit IgG | Abcam # Ab6721 | 1/5000 | WB |

**Table of antibodies used for immunofluorescence studies.**

| Antibody | Source / Product no. | Dilution | Analysis |
|---|---|---|---|
| rabbit IgG polyclonal anti-laminin | Sigma-Aldrich # L9393 | 1 :750 | IF |
| mouse IgG2b monoclonal anti-MHC type I | DSHB # BA-F8 | 1:25 | IF |
| mouse IgG1 monoclonal anti-MHC type IIa | DSHB # SC-71 | 1 :200 | IF |
| mouse IgM monoclonal anti-MHC type IIb | DSHB # BF-F3 | 1 :200 | IF |
| p62/SQSTM1 | Novus Biologicals Inc. clone 2C11 | 1/200 | IF |
| Alexa Fluor 350 IgG2b (y2b) goat anti-mouse | Invitrogen, A-21140 | 1:500 | IF |
| Alexa Fluor 488 IgG goat anti-rabbit | Invitrogen, A-11008 | 1:500 | IF |
| Alexa Fluor 488 IgM goat anti-mouse | Invitrogen, A-21042 | 1:500 | IF |
| Alexa Fluor® 488 IgG2a goat anti-mouse | ThermoFisher, A-21131 | 1:500 | IF |
| Alexa Fluor 594 IgG1 (y1) goat anti-mouse | Invitrogen, A-21125 | 1:100 | IF |
| Alexa Fluor 594 IgG goat anti-rabbit | Invitrogen, A-11037, | 1:500 | IF |
| Alexa Fluor 568 IgG goat anti-rabbit | ThermoFisher, A-11011 | 1:500 | IF |

*All primary antibodies targeting MHCs were purchased from the Developmental Studies Hybridoma Bank (DSHB, University of Iowa, IA).

### Table of primers for qPCR

| | Forward primer (5'-3') | Reverse primer (3'-5') |
|---|---|---|
| *18S* | TGCGGTTTAGCGTCGGTGTC | CCAAGTGGCCAAAGCGTA |
| *Atg7* | TGGCTGCTACTTCTGCAATGATGT | CAGGACAGAGACCATCAGCTCCAC |
| *LC3B* | CGATACAAGGGGGAGAAGCA | ACTTCGGAGATGGGAGTGGA |
| *Bnip3* | TTCCACTAGCACCTTCTGATGA | GAACACCGCATTTACAGAACAA |
| *Bnip3L* | TTGGGGCATTTTACTAACCTTG- | TGCAGGTGACTGGTGGTACTAA |
| *p62/Sqstm1* | GCACCTGTCTGAGGGCTTCT | GCTCCAGTTTCCTGGTGGAC |
| *GabarapL* | GAGGACCACCCCTTCG | CGGAGGGCACAAGGTACTTC |
| *Park2* | TCTTCCAGTGTAACCACCGTC | GGCAGGGAGTAGCCAAGTT |
| *Beclin1* | CTTGGAGGAGGAGAGGCTGA | TGTGGAAGGTGGCATTGAAG |
| *Ctsl/Cathepsin L* | CGGGTTGCCTAGAAGGACAG | ACAGCCCTGATTGCCTTGAT |
| *Foxo3a* | CCGGCTCCTTCAACAGTACC | TGAAGCAAGCAGGTCTTGGA |
| *Sod1* | GGCAATGTGACTGCTGGAAA | GCAATCCCAATCACTCCACA |
| *Sod2* | CCACACATTAACGCGCAGAT | AGGGCTCAGGTTTGTCCAGA |
| *Catalase* | CTGGAGTCTTCGTCCCGAGT | CTTCCTGCCTCTCCAACAGG |
| *Trim32* | CAGAGTGAGGTGCTGGTTGC | GGCTCCAAGGAAGCTTAGCA |
| *Nedd4* | CATAGGCCTGGCCAAGAAAG | GCTGTGGAAGGACCCTGAAC |
| *Fbxo30* | TCGTGGAATGGTAATCTTGC | CCTCCCGTTTCTCTATCACG |
| *Fbxo32* | TGGGTGTATCGGATGGAGAC | TCAGCCTCTGCATGATGTTC |
| *Trim63* | TGCTTGGCACTTGAGAGGAA | AGAAGCTGGGCTTCATCGAG |
| *Pa28a/Psme1* | GCGGAAGAAGCAACAGGAG | GGTTTTAGGCGTTGCAGGAG |
| *Pa28ß / Psme2* | ATTTTGGGGTGGCAATTCAG | GCTTCATCTCGCTCATGCAC |
| *Rpn6 / Psmd11* | CATTTGGCCAAGCTGTACGA | TCGGCCTTGGAGAGTTTGAT |
| *Rpn12 / Psmd8* | GAAATCGCAGGATGCATTGA | GAGGGGATGGTGCTGTCTTC |
| *B5 / Psmb5* | GTACAAAGGCATGGGGCTGT | CGGTCCCAGAGATCCTGTTC |
| *B1 / Psmb6* | GCAGTTCACTGCCAATGCTC | CAACGTGGCAATGGTGAACT |
| *B2 / Psmb7* | TTGTCGCAGGAATGCTGTCT | CAGCAACAACCATCCCTTCA |
| *B5i / Psmb8* | TACCTGCTTGGCACCATGTC | CGTTCCCCATTCCGAAGATA |
| *B1i / Psmb9* | GGACGGAAGAAGTCCACACC | GTGCAGAGGGGAGAGCTTGT |

| | | |
|---|---|---|
| *B2i / Psmb10* | GCTGCGGACACTGAGATGAC | TTGGTACCGGAAAAGCGTCT |
| *Tweak* | CAGGAGGAGCTGACAGCAGA | AGGCCGGACTAGTTGTTCCA |
| *Fn14* | GGATTCGGCTTGGTGTTGAT | GCAGAAGTCGCTGTGTGGTC |
| *IL-6* | CACGGCCTTCCCTACTTCAC | TGCAAGTGCATCATCGTTGT |
| *IL-6R* | CACCTGCCAACCTTGTGGTA | AGGTCGGTATCGAAGCTGGA |
| *IL-1ß* | TCGCAGCAGCACATCAACAA | TGGAAGGTCCACGGGAAAGA |
| *Tnfa* | ACTGGCAGAAGAGGCACTCC | CTCCAGCTGCTCCTCCACTT |
| *Tnfa-R* | ACCGTGTGTAACTGCCATGC | ATTTGCAAGCGGAGGAGGTA |
| *IL-18* | TGCTTGCCAAAAGGAAGATG | CACAAACCCTCCCCACCTAA |
| *IL-18R* | ATAGGCGCATAGCGGAAAGA | GACCCTGGGTAACGTCTCCA |
| *IL-24* | GCACTGGCCCTTTCTTCAAC | CAGAAGGCCTCCCACAGTTC |
| *Mstn* | GTGGAAAAAGAGGGGCTGTG | GATCAGTTCCCGGAGTGGAG |
| *ACT RIIB* | GGCCACAAGCCTTCTATTGC | GCCAACCTGTCCATGGGTAT |
| *Fgf21* | TCCTGGGTGTCAAAGCCTCT | ATCCTGGTTTGGGGAGTCCT |
| *Pgc1-$\alpha$* | TTGCTAGCGGTCCTCACAGA | GTCAGGCATGGAGGAAGGAC |

### Software and Algorithms

| **Software and Algorithms** | **Source** | **Identifier** |
|---|---|---|
| GraphPad Prism 8 software | N/A | https://www.graphpad.com/scientific-software/prism/ |
| Igor Pro (Version 6.37) | N/A | WaveMetrics |
| ImageJ Software | N/A | https://imagej.nih.gov/ij/ |
| TSE PhenoMaster software | N/A | https://www.tse-systems.com/ |
| Affymetrix® Transcriptome Analysis Console (TAC) Software | N/A | Thermo Fisher Scientific |
| ImageLab software | N/A | Bio-Rad Laboratories |
| R package for Microarray Data (Limma) | (1) | https://www.bioconductor.org/packages/release/bioc/html/limma.html |

1. Ritchie ME, Phipson B, Wu D, Hu Y, Law CW, Shi W, et al. limma powers differential expression analyses for RNA-sequencing and microarray studies. *Nucleic Acids Res.* 2015;43(7):e47.
