## Supplemental file S2 for "Role of autophagy in sepsis-induced skeletal muscle dysfunction, whole-body metabolism, and survival"

### Supplemental file S2.1: Networks predicted by Ingenuity Pathway Analysis in Atg7 deficient skeletal muscle of septic

| **Top Diseases and Functions and Network Map** | **Molecules in Network** | **Score** | **Focus Molecules** |
|---|---|---|---|
| Cell Cycle, Cell Death and Survival, Cell-To-Cell Signaling and Interaction<br>**Network Map 1** | ANXA4,ARF6,ATP6V0C,BYSL,CYP19,DDA1,FHIT,FLII,G0S2,GFER,GNL2,GTPBP4,HIF3A,IKK (complex),LTV1,NFkB (complex),NOP14,PIAS4,PIM3,PLAA,PPM1G,PPP4C,RAB31,RIN3,RNF31,SLC2A1,STK40,TAX1BP1,TNFSF18,TPMT,USP11,USP20,VHL,WWP2,ZFAND6 | 31 | 32 |
| Cellular Compromise, Cellular Function and Maintenance, Nutritional Disease<br>**Network Map 2** | AR,ATF3,Calmodulin,COPA,COPG1,DHCR7,DNA-methyltransferase,DNAJB9,EIF2A,GBE1,HDL,HSP90B1,HSPA5,JMJD1C,KCNQ5,LRIG1,LRIG3,MAP3K14,MAT2A,NFKB2,NPC1,NPLOC4,RAB2A,RNF123,SEC16A,SEC61A1,SLC37A4,SMYD1,TACC2,TBRG1,TNFRSF12A,UBA1,UBR4,URI1,XBP1 | 31 | 32 |
| Post-Translational Modification, Cancer, Organismal Injury and Abnormalities<br>**Network Map 3** | 20s proteasome,26s Proteasome,ADRM1,ATF6B,ATG7,BAG3,BNIP3L,DYSF,FKHR,IBTK,KEAP1,LGR4,MAP1LC3,MAP1LC3B,MYOD1,MYOG,NBR1,NDUFA13,NR4A2,PSMB1,PSMC2,PSMC3,PSMC4,PSMC5,PSMC6,PSMD1,PSMD14,PSMD2,PSMD4,PSMD6,PSMD8,SQSTM1,STAT3,UBFD1,USP14 | 29 | 31 |
| Connective Tissue Development and Function, Organ Morphology, Organismal Development<br>**Network Map 4** | ARRDC3,CPE,DDIT4,DDX21,ESR1,Filamin,FOSL2,GHR,GPAM,Growth hormone,HNRNPF,HSF2BP,HSPA13,INPPL1,IPO5,KPNB1,MAP1B,MIER1,MINOS1,MYO1C,PDCD6,PDCD6IP,PP1 protein complex group,PPARGC1B,PSMB4,PTTG1IP,SEC31A,Shc,SMAD6,TNPO1,TPD52L2,TTC9,WDR77,WWOX,ZNF367 | 29 | 31 |
| Cell Signaling, Post-Translational Modification, Protein Synthesis<br>**Network Map 5** | Alpha tubulin,ANXA11,CDK4/6,DAXX,ERRFI1,Gamma tubulin,Histone h3,IPO4,KLHDC10,LAMB2,NDUFA5,NDUFA9,NDUFAF1,NDUFAF5,NDUFAF6,NDUFAF7,NDUFB4,NDUFB6,NDUFB7,NDUFC1,NDUFC2,NDUFS3,NDUFV1,NDUFV3,RASSF1,RPS9,SPOP,TUBA1C,TUBB,TUBB2A,TUBB4B,UQCRC1,WDR5,WLS,ZNF423 | 29 | 31 |
| Lipid Metabolism, Small Molecule Biochemistry, Neurological Disease<br>**Network Map 6** | ADM,AMPK,APOD,ARL4A,BRI3,Complement component 1,CREM,HGS,HNRNPU,HRAS,IDH3A,IFIT3,IL6ST,KCNK5,KLF11,MAPT,MDH1,MDH2,NCL,NT5C1A,PA2G4,PABPN1,PHB2,POLR1E,PRELID1,RPL3,RRP9,SDHA,SDHAF2,SERPINA3,STAU1,TCF,TCR,VPS37C,YWHAZ | 29 | 31 |

| Top Diseases and Functions and Network Map | Molecules in Network | Score | Focus Molecules |
|---|---|---|---|
| Cell Cycle, Connective Tissue Development and Function, Cell Death and Survival **Network Map 7** | BAG1,Cbp/p300,CDKN1A,CSTB,CSTF1,Cyclin B,Cyclin D,DDB1,DDX24,DLAT,E2F4,EP400,FDFT1,GPS2,H2AFZ,HP,HSPA4,LIN9,LPIN1,MED15,MGMT,NDUFAB1,NFE2L1,NFIL3,NOC2L,Nr1h,PDZD2,PPP1R13B,PRDX3,PSAT1,PSMA3,RRM1,Smad,SRXN1,TFDP1 | 27 | 30 |
| Cell Death and Survival, Cell Cycle, Cellular Development **Network Map 8** | Actin,ACTL6A,AIMP2,caspase,CD14,CD3,CDK4,CEBPD,CENPO,Cyclin A,DHPS,E2f,EEF1E1,ENDOG,ETS2,FAM83D,FAS,ITGAM,JUND,KRAS,LMNA,MARS,PDE4A,PDE4B,PLA2G16,PSD3,PSMA7,RASSF9,RB1,RYR1,SMARCC1,SOD2,TFDP2,VLDLR,ZNF385B | 27 | 30 |
| RNA Post-Transcriptional Modification, Cancer, Developmental Disorder **Network Map 9** | ACHE,CCDC59,CCNL2,CLIC4,COQ9,EBAG9,EGR1,EPDR1,EXOSC1,FOXK1,GID4,H2AFX,HEATR1,KIF5B,Lfa-1,MAP2K1/2,MTA1,P-TEFb,PCNA,PLCD4,PSMD7,RANBP10,RANBP9,RNA polymerase II,Rnr,RRP12,SBDS,SDAD1,SND1,TMA16,TRAFD1,TSR1,WDR43,WDR46,ZNF318 | 27 | 30 |
| Cellular Development, Cellular Growth and Proliferation, Connective Tissue Development and Function **Network Map 10** | Alpha catenin,BCR,CDH4,Cdk,CNTNAP2,CYTH2,DPAGT1,EIF3B,FBXO32,GADD45A,GCLM,Gcn5l,Hdac,HSPH1,IFRD1,JUP,KLF6,KLF9,LAMP2,MAX,MYC,NAP1L1,ODC1,PARP10,RFFL,Rock,SERINC3,SETD7,SLC25A19,ST3GAL3,TAF1A,thymidine kinase,TMEM126A,UTP18,ZMIZ2 | 26 | 29 |
| Hereditary Disorder, Organismal Injury and Abnormalities, Skeletal and Muscular Disorders **Network Map 11** | ACP5,ADGRG1,AES,BCKDK,CD68,CDH19,CG,CH25H,EGLN,EMP1,estrogen receptor,FAM117B,FLNC,FSH,G-protein beta,GPRC5B,IGFBP3,ING1,Lh,MAPK6,MPI,NOL3,PDLIM3,PEX11A,PSMD3,RAB27A,RAB33B,STC1,TGFB3,THBS1,TIMP4,TLN1,USF2,VIPAS39,XPR1 | 26 | 29 |
| Cancer, Neurological Disease, Organismal Injury and Abnormalities **Network Map 12** | ADAM10,Akt,AMOT,ANGPT1,ANKRD1,BTG2,CD151,CDC42EP4,CHRNA4,CNKSR1,Fgfr,FLOT1,GSR,HPCAL1,Integrin alpha 3 beta1,NOLC1,NRF1,PITPNA,POR,PREP,PRR5,Rac,Rar,RARB,Ras homolog,RNF7,Rxr,SNF8,TGM1,THOC5,TMED10,TRIM16,TSPAN12,UQCRC2,USP4 | 24 | 28 |

| **Top Diseases and Functions and Network Map** | **Molecules in Network** | **Score** | **Focus Molecules** |
|---|---|---|---|
| Cell Death and Survival, DNA Replication, Recombination, and Repair, Cancer **Network Map 13** | ALKBH3,ARHGAP35,BCL2L1,BNIP3,Caspase3/7,CDC42EP3,CPEB4,EIF3,EIF3A,EIF3C,EIF3G,EIF3M,ERCC1,ERCC4,FOXO4,MCPH1,NEU2,PARK7,PARP,PEPCK,Pkc(s),PSMD11,RAD52,RHEB,S100A8,S100A9,SH3KBP1,SIM2,SLAMF7,SNRPB,SRPK1,TFIIH,U1 snRNP,VDAC1,XPA | 24 | 28 |
| Cell-To-Cell Signaling and Interaction, Cellular Growth and Proliferation, Hematological System Development and Function **Network Map 14** | ARNT,ATF4,BPGM,CA9,CERS2,Ck2,CXCL10,CYP1B1,DNAJC15,DNAJC3,FCGR2B,GPD2,Ifn gamma,IFN type 1,Iga,IL12 (complex),IL12A,IL15,JAK,JUN,MERTK,MYD88,N-cor,PGAM2,PGD,PGK1,TAB2,TCOF1,Tgf beta,TGFBR2,TIRAP,TNFSF10,TXN,ZFP36,ZNF503 | 22 | 27 |
| Cancer, Organismal Injury and Abnormalities, Cellular Development **Network Map 15** | ACVR1,ALKBH8,APLP2,BCR (complex),CCNK,CCT2,CCT4,CD44,CTTNBP2NL,Cyclin E,DNAPK,DYNC1H1,HIGD1A,HSP90AA1,IFITM2,Igm,JUNB,KLF4,LOX,mediator,PLCL1,RGCC,Secretasegamma,SMAD1/5,Smad2/3,SMARCD3,SNAI2,SNAPC5,STAC3,STRN4,TCP1,TGFBR1,TRMT112,TUBB6,UCP2 | 22 | 27 |
| Cell Death and Survival, Amino Acid Metabolism, Post-Translational Modification **Network Map 16** | 14-3-3,ACACB,AKT1,ARHGDIA,BACH1,calpain,CLK2,Collagen Alpha1,Cpla2,EPM2A,GRB14,Gsk3,Hsp27,HTATIP2,ILF3,JINK1/2,LARP1,MAP3K6,MTOR,MYBBP1A,Notch,NSUN2,PDPK1,PI3K(family),PKM,SSH2,SSRP1,TNKS,TRIM35,TTC23,UVRAG,Vegf,YWHAB,YWHAH,ZNF598 | 19 | 25 |
| Developmental Disorder, Hereditary Disorder, Neurological Disease **Network Map 17** | ATP6V0D1,CALR,CANX,CARM1,DDX20,DHX9,EMID1,ENPP2,FBXO6,GEMIN5,GIMAP4,Hsp70,Hsp90,HSP90AB1,IFN Beta,IgG,Immunoglobulin,Interferon alpha,KCNH2,LAP3,MHC Class I (complex),MRPL18,MUS81,MYH14,PBX1,PI3K p85,PRG4,SERPINB1,SNRPF,SYNPO,Tap,Ubiquitin,UBXN4,VCP,YBX2 | 19 | 25 |
| Cell Death and Survival, Organismal Injury and Abnormalities, Connective Tissue Development and Function **Network Map 18** | ABCB6,ABCC1,ANXA6,AQP4,ATP synthase,ATP5MD,c-Src,C1QBP,CDC42EP2,Creb,cytochrome-c oxidase,DAPP1,DNM1L,ERK,F Actin,GPNMB,IGF1R,IMPDH2,INHA,INSR,IRS2,LCN2,MAOA,Mitochondrial complex 1,Mlc,NDUFB11,NDUFS1,NDUFS4,P glycoprotein,PI3K (complex),PIM1,Pka,PTPN1,UGCG,UQCRFS1 | 18 | 24 |

| **Top Diseases and Functions and Network Map** | **Molecules in Network** | **Score** | **Focus Molecules** |
|---|---|---|---|
| Cell Morphology, Cellular Movement, Gastrointestinal Disease **Network Map 19** | ATP1A2,CCL11,EDNRB,EIF4B,EIF4EBP1,ERK1/2,Focal adhesion kinase,GRM1,Hif1,Integrin,ITGA4,ITGB7,MAP4,MKNK2,MTORC1,P2RY1,p70 S6k,PIM,Pkg,Pld,PP2A,PPIA,PSMB10,PTPN12,Rap1,RAP1B,RAPGEF1,RASGRP3,RASSF4,Rsk,SDHD,SHC3,SLC36A1,TSPYL2,VEGFA | 17 | 23 |
| Cell Death and Survival, Organismal Injury and Abnormalities, Renal Necrosis/Cell Death **Network Map 20** | Ap1,CFL1,Collagen(s),CTGF,CYCS,DUSP10,EGLN3,FOXO1,GPX3,HAX1,HSPB1,HYOU1,IER3,ITCH,ITGB6,Jnk,LDL,MAFF,Mapk,MCL1,Mek,PDGF BB,PLK2,Ppp2c,PPP2R2D,PRKAA,Raf,RAS,Rb,RHOB,RXRG,SLC25A33,TAF4B,TXNRD1,UCN | 17 | 23 |
| Cellular Function and Maintenance, Molecular Transport, Small Molecule Biochemistry **Network Map 21** | ABCB7,Aconitase,ARF4,CD3 group,FBXO31,GAA,GBA,HLA-DR group,IL1,IL12 (family),IL23,ISCA1,MAC,MAP2K6,MYL12B,NFS1,NPHS1,NUBP1,P38 MAPK,PLA2G6,PLA2G7,PLC gamma,RHOC,RRAD,SAA,SH3BP2,SIRT3,SLC7A5,SRC (family),succinate dehydrogenase,SYK/ZAP,TFRC,TLR4,TRIB1,UBE2E1 | 15 | 22 |
| Cardiovascular Disease, Hereditary Disorder, Metabolic Disease **Network Map 22** | AS3MT,BAG5,CCDC134,CERK,COA5,CROT,CYB5B,NUP50,NUPR1,PEX12,RAB29,RANBP6,RTN4IP1,SERTAD2,SPAG5,USP31,ZPR1 | 14 | 15 |
| Organismal Survival, Embryonic Development, Organismal Development **Network Map 23** | ALDOA,BHLHE41,CASP4,CNTFR,FKBP10,GPNMB,HDAC4,hemoglobin,Hif1,HIF1A,HP,IL1A,INHBA,INHBB,IRS2,LCN2,LMOD2,LOX,MB,MEF2C,mir-199,mir-302,NOV,PIR,PLOD2,RETREG1,SERPINA3,SLC16A4,SLC2A1,SREBF1,STRBP,SUCLG1,TGFB3,TMEM128,TRIM63 | 13 | 20 |
| Cellular Movement, Endocrine System Disorders, Gastrointestinal Disease **Network Map 24** | ADI1,ANKRD28,ANLN,ANXA7,AURK,BCAR1,BTG2,CMBL,Cofilin,COL10A1,COL1A1,COL24A1,EDA2R,GLI1,GNAS,JUNB,miR-296-5p (miRNAs w/seed GGGCCCC),MMP14,MTMR1,NRAP,NREP,PDGFRA,PSTPIP2,PVR,RHOC,RUNX2,SEMA4D,TCHP,TRAM2,TTC13,VEGFA,WDR74,WT1,WWOX,ZMYND10 | 13 | 20 |
| Cellular Function and Maintenance, Cell Morphology, Cellular Assembly and Organization **Network Map 25** | AGTR1,AHI1,APOLD1,ARL3,CRCT1,EIF6,EMD,EXTL2,GPATCH4,HDLBP,HECTD1,KIF3A,LBR,LCE1A,LLGL1,LMCD1,LMNB1,MLLT11,MRPS35,MYH10,MYO18A,NCKAP1,NUPR1,P4HB,PDE6D,PLEKHA5,PMP22,PRKCI,PRMT1,RAB11FIP5,RAB28,RNH1,SRF,SURF4,YWHAG | 13 | 20 |

### Network Map 1

### Cell Cycle, Cell Death and Survival, Cell-To-Cell Signaling and Interaction

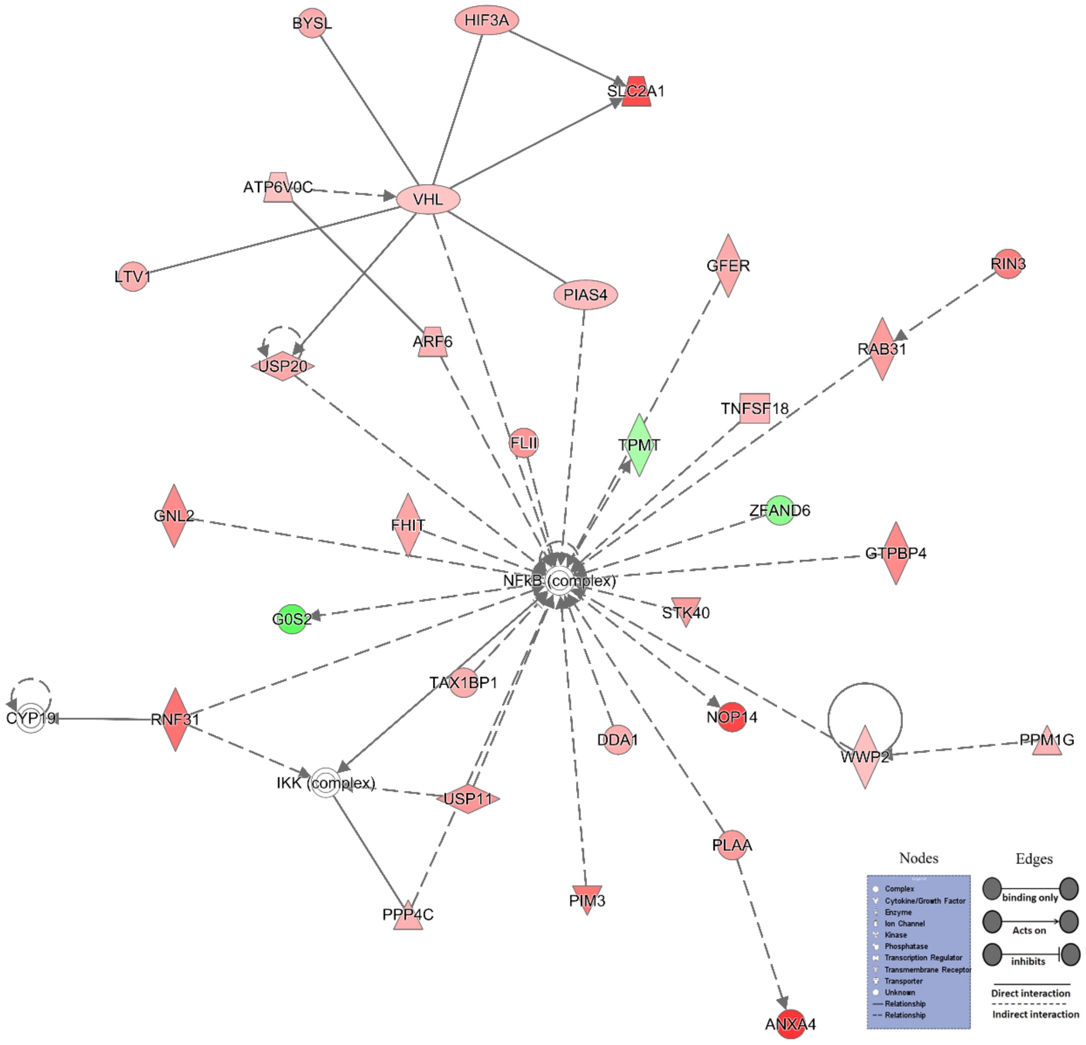


Representation of network generated by IPA clearly showing the upregulation of group of genes associated with cell death. Nodes in red indicates upregulation, whereas nodes in green color shows downregulation.

**Network Map 2**

**Cellular Compromise, Cellular Function and Maintenance, Nutritional Disease**

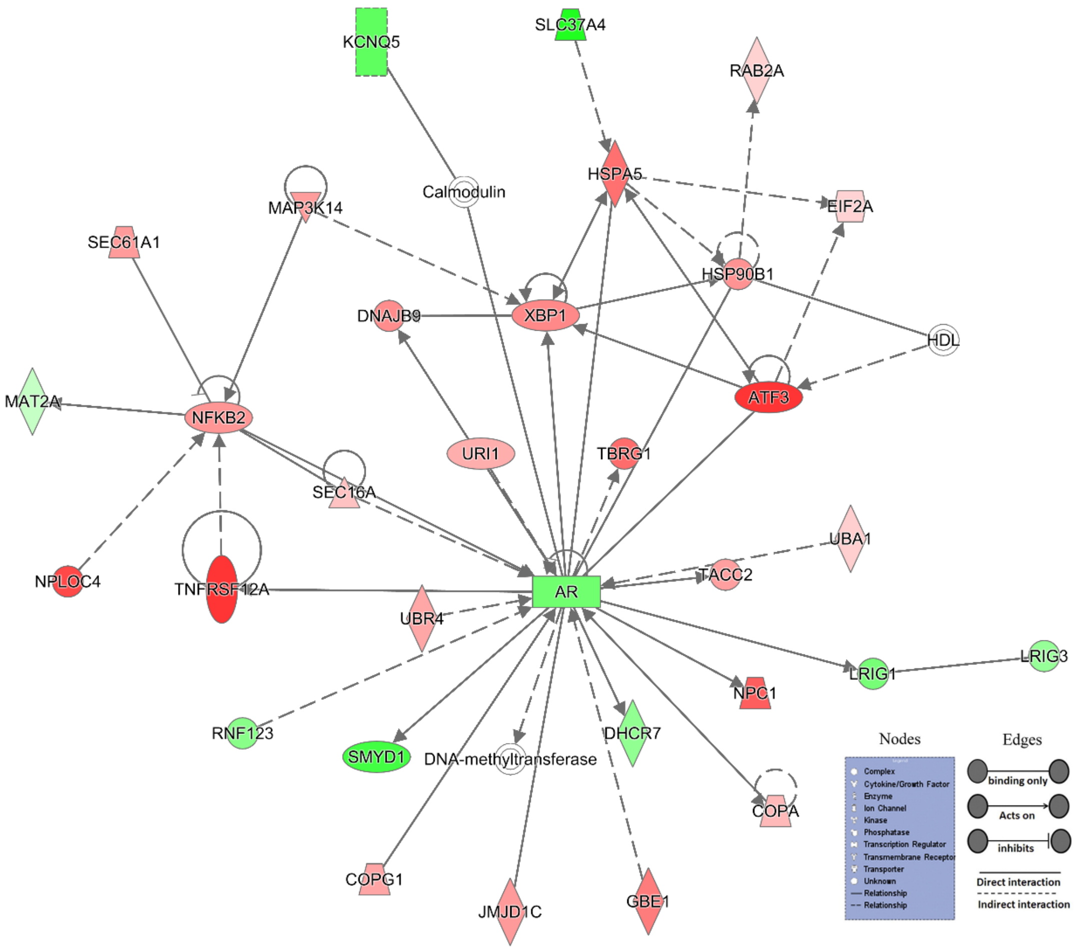


Representation of network generated by IPA clearly showing the upregulation of group of genes associated with cellular function and maintenance. Nodes in red indicates upregulation, whereas nodes in green color shows downregulation.

### Network Map 3

### Post-Translational Modification, Cancer, Organismal Injury and Abnormalities

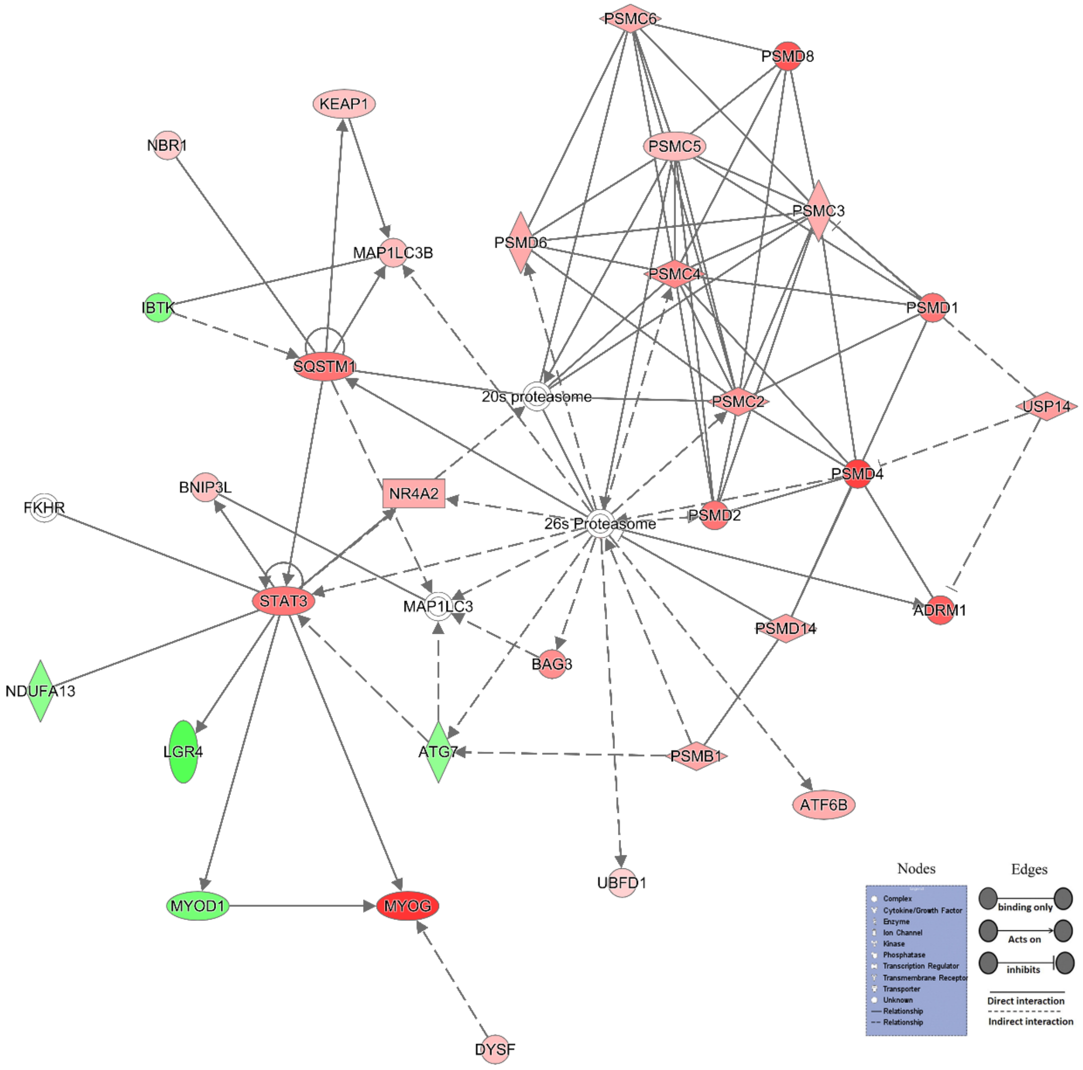


Representation of network generated by IPA clearly showing the upregulation of group of genes associated with post-translational modification. Nodes in red indicates upregulation, whereas nodes in green color shows downregulation.

**Network Map 4**

**Connective Tissue Development and Function, Organ Morphology, Organismal Development**

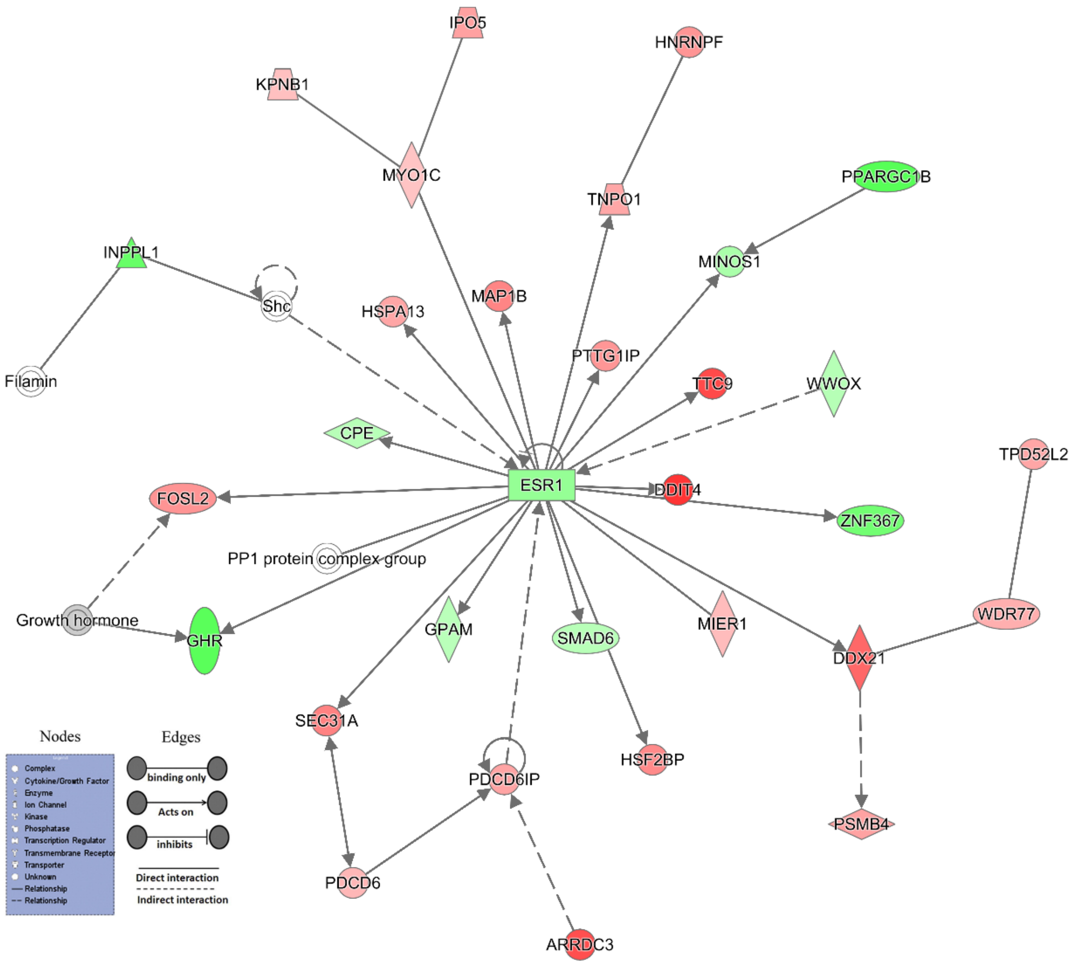


Representation of network generated by IPA clearly showing the downregulation of group of genes associated with connective tissue development and function. Nodes in red indicates upregulation, whereas nodes in green color shows downregulation.
